# Molecular Tracking and Cultivation Reveal Ammonia-Oxidizing Archaea as Emerging Commensals of the Human Skin Microbiome

**DOI:** 10.1101/2024.08.05.606590

**Authors:** Alexander Mahnert, Maximilian Dreer, Ülkü Perier, Michael Melcher, Stefanie Duller, Adina Lehnen, Theodora Goessler, Daniela Brunner, Thomas Graier, Peter Wolf, Rafael I. Ponce-Toledo, Logan H. Hodgskiss, Melina Kerou, Christine Moissl-Eichinger, Christa Schleper

## Abstract

**Background:** Ammonia-oxidizing archaea (AOA) have repeatedly been detected on human skin, yet their persistence, physiological traits, and adaptations remain poorly understood. Here, we identify *Nitrosocosmicus* species as integral members of the healthy skin microbiome using two complementary approaches.

**Results:** Through cultivation, we enriched two autotrophic strains, *Candidatus Nitrosocosmicus epidermidis* and *Ca. Nitrosocosmicus unguis*, from human skin samples. Genomic analyses revealed specific adaptations for skin colonization, including genomic islands and expanded gene families linked to interactions with host proteins and signaling pathways, distinguishing these AOA from their soil-dwelling relatives.

Parallel molecular profiling in cross-sectional and longitudinal cohorts (n=47) consistently detected *Nitrosocosmicus* particularly in sebaceous areas. Co-occurrence patterns with specific bacterial taxa reinforce their role as stable components of the skin microbiome.

**Conclusions:** These findings indicate that *Nitrosocosmicus* species are emerging commensals, evolutionarily capable of transitioning from soil to human skin, where they likely play a critical role in the skin ecosystem.

## Background

The human body is an ecosystem populated by spatially organized microbial communities composed of bacteria, archaea, viruses/phages and small eukaryotes, with diverse metabolic activities and interactions: the microbiome(Berg et al., 2020). In recent years, numerous studies have led to a better understanding of the mechanisms involved in the complex interplay between the host and its commensals and have revealed the fundamental role of the human microbiome in health and disease (Fan and Pedersen, 2020; Flowers and Grice, 2020; Gomaa, 2020). However, most studies on human microbial communities have focused exclusively on the bacterial microbiome, while other integral components, such as eukaryotes/fungi, archaea and viruses have remained largely overlooked (Bang and Schmitz, 2015; Borrel et al., 2020; Fan et al., 2023; Gaci et al., 2014; Nayfach et al., 2021). A systematic metagenomic study has recently revealed the extent of diversity and abundance of archaea in the human gut microbiome (Chibani et al., 2022), where the increased presence of methanogenic archaea affects short-chain fatty acid production and the overall host physiology (Kumpitsch et al., 2021).

The skin microbiome is similarly complex and harbors various bacterial species, including *Staphylococcus, Streptococcus, Corynebacterium, and Propionibacterium* (now *Cutibacterium*), and fungi such as *Malassezia* (Grice and Segre, 2011; Kong, 2011). The composition and diversity of the microbiome varies across different body sites (e.g. oily versus dry skin) and between individuals (Oh et al., 2016). As in the gut, microorganisms play essential roles in maintaining skin health as they provide protection against pathogens by competing for nutrients and space, producing antimicrobial compounds and modulating the skin’s immune response (Byrd et al., 2018; Ross et al., 2017).

Human skin is also colonized by archaea (Moissl-Eichinger et al., 2017; Probst et al., 2013), in particular by representatives of the ammonia oxidizing archaea (AOA), formerly Thaumarchaeota, now class Nitrososphaeria and phylum Thermoproteota (Parks et al., 2022, 2020; Rinke et al., 2021). Although rather rare in abundance, archaeal signals were found in human subjects analyzed by 16S rRNA gene sequencing and detection of archaeal *amoA*, the gene for subunit A of the key enzyme ammonia monooxygenase used for ammonia oxidation (Probst et al., 2013). AOA signatures were detected on all human subjects and were closely related to those found earlier in cleanrooms and intensive care units (Moissl-Eichinger, 2011). Their presence was confirmed by fluorescence *in situ* hybridization (Probst et al., 2013), 454 pyro-sequencing (Moissl-Eichinger et al., 2017), and infrared hyperspectral imaging (Moissl-Eichinger et al., 2017). Although the physiological relevance of AOA to the skin is still subject to speculation, its potential role is intriguing considering that urea and ammonia are components of human sweat and are the main energy sources of ammonia oxidizers. Also, nitric oxide (NO), an important signaling molecule involved in diverse processes in humans, is a byproduct of the ammonia oxidation process (Jessica A. Kozlowski et al., 2016).

Detection of archaea in human microbiomes is generally hampered by numerous methodological obstacles, such as challenging cultivation (Alves et al., 2019; Klein et al., 2022; Sun et al., 2019) and molecular detection issues (summarized in (Mahnert et al., 2018)), because of the bacteria-centric orientation in sampling, cell lysis, DNA extraction protocols, PCR primers, and classification databases. Consequently, archaeal signals often remain close to or below the detection limit, making it difficult to understand the role of archaea on human skin (Umbach et al., 2021). Such obstacles and the lack of cultivated strains of skin AOA have left many questions unanswered, with respect to i) abundance, ii) longitudinal stability, iii) association with human health, iv) physiology and metabolism, v) interaction with the bacterial microbiome, and vi) specific adaptations to the human skin.

In this manuscript, we present the results of two complementary, but independent approaches designed to investigate the persistence and characteristics of AOA on human skin: extensive cultivation efforts and molecular analyses targeting AOA in both healthy and diseased individuals.

These efforts yielded stable, highly enriched cultures of strains affiliated with the genus *Candidatus* Nitrosocosmicus from human skin. Their genomes reveal features that could facilitate colonization of the skin environment, but also highlight the genomic repertoire shared among all members of the genus *Ca*. Nitrosocosmicus, a widespread environmental group of AOA. Molecular profiling further tracked *Ca.* Nitrosocosmicus signatures across cohorts of healthy volunteers and patients, identifying correlations with physical parameters and bacterial co-occurrence patterns.

Together, these findings suggest that specific *Ca.* Nitrosocosmicus species or clades have evolved distinct traits potentially enabling their transition from soil to specialized niches on human skin, supported by genomic adaptations and their stable integration into the skin microbiome.

## Methods

### Initial cultivation

Prior to enrichment setups, subjects not working in an experimental laboratory were pre-screened for the presence of *Nitrososphaerales* by targeted 16S rRNA gene amplification. We used 3 different primer pairs, two of which targeted 16S rRNA (Cren 771F/957R (Ochsenreiter et al., 2003) and Arch 109F/1492R (Großkopf et al., 1998; Nicol et al., 2008) and one targeted the *amoA* gene (19F/629R, (Arce et al., 2018; Pester et al., 2012)). In dilution tests with template DNA primers Arch 109F/1492R turned out to be the most sensitive and were thus used to screen individuals for the presence of AOA. All tested individuals (n=41) were positive for AOA. In a first round of cultivation attempts diverse swabs and brushes were tested to identify tools that are compatible with AOA growth. Sterile polyurethane single sponge swabs (BD BBL™ CultureSwab™ EZ, Becton Dickinson, USA) were identified as most suitable for enrichments, since they promoted growth of *Nitrososphaera viennensis* in pre-tests, while several other materials turned out to be inhibitory for growth.

For enrichment cultures, sterile swabs premoistened with sterile freshwater medium (FWM (De La Torre et al., 2008; Tourna et al., 2011)) were used to sequentially rub the face, neck, upper body, back, arms, and legs to obtain as much biomass as possible. All following procedures were performed under sterile conditions (laminar flow). The polyurethane swab was removed from the plastic shaft and directly added to FWM (Enrichments T1S, X2B and Z3A). For enrichment of R2S, free edges of a subjects’ fingernails (two to three clippings each obtained with a sterilized scissor) were directly added to FWM (Enrichment R). Earlier studies had shown the suitability to use fingernail microbiomes to assess the skin microbiome, due to scratching (Rayan and Flournoy, 1987).

All initial enrichments were set up in sterile 30 ml polystyrene screw cap tubes (Greiner) containing 20 ml FWM supplemented with 0.5 mM NH4Cl, 20 µl non-chelated trace element solution, 7.5 µM FeNaEDTA, 20 µl vitamin solution as well as 25 µg/ml carbenicillin and 25 µg/ml kanamycin (for detailed medium composition see table 1 in (Reyes et al., 2020; Tourna et al., 2011)).

**Table 1:** Genomic and growth features of *Ca.* Nitrosocosmicus epidermidis, *Ca.* Nitrosocosmicus unguis compared to other cultivated representatives of the genus *Ca.* Nitrosocosmicus

| Organism<sup>#</sup> | Status | Growth temp (°C) | pH | Genome size (Mb) | No. of contigs | Completeness [%] | Contamination [%] | DNA GC % | protein coding genes | 16S/23S/5S rRNA gene copies | reference |
| --- | --- | --- | --- | --- | --- | --- | --- | --- | --- | --- | --- |
| *Nitrosocosmicus unguis* R2S | Enriched | 28 | 7 | 3.21 | 1 | 97.27 | 0.61 | 33.77 | 3597 | 3/3/1 | this study |
| *Nitrosocosmicus epidermidis* T1S | Enriched | 28 | 7 | 3.07 | 1 | 99.94 | 2.2 | 33.55 | 3401 | 3/3/1 | this study |
| *N. oleophilus* MY3 | Pure | 30 | 6.5-7 | 3.43 | 1 | 99.76 | 0.54 | 34.14 | 3722 | 3/3/1 | Jung et al. 2016 |
| *N. franklandianus* C13 | Pure | 40 | 7 | 2.84 | 1 | 99.22 | 0.15 | 34.07 | 3025 | 2/2/1 | Lehtovirta-Morley et al. 2016 |
| *N. arcticus* Kfb | Enriched | 28 | 6 | 2.65* | 23 | 98.67 | 0.19 | 34 | 2970 | 3/3/1 | Alves et al. 2019 |
| *N. hydrocola* G61 | Enriched | 33 | 8 | 2.99 | 1 | 99.76 | 0.48 | 33.94 | 3162 | 2/2/1 | Sauder et al. 2017 |
| *N. agrestis* SS | Enriched | 37 | 6.5-7 | 3.22** | 43 | 96.1 | 2.91 | 33.42 | 3524 | 2/2/1 | Liu et al. 2021 |

<sup>#</sup> all organisms have Candidatus status

Variations between initial enrichments included the carbon source (2 mM Na2CO3 or NaHCO3), pH (7 or 8.5), temperature (28°C or 32°C) supplementation with organic substrates (0.5 mM acetate and 0.5 mM pyruvate or none) and filter sterilized spent growth medium of *Nitrososphaera viennensis* EN76 (Tourna et al., 2011) and *Ca.* Nitrosocosmicus franklandianus C13 (Lehtovirta-Morley et al., 2016) (2.5% (v/v) (Supplementary Table S1). Cultures were incubated aerobically for up to several months without shaking and in the dark. First transfer of cultures was performed after 3 months and *amoA* sequences and 16S rRNA gene sequences were determined for the initial enrichments that showed nitrite production as well as for their first transfers (see Supplementary Table S2). The sequences were identical in the original cultures and the corresponding transfer. For T1S enrichments 6 different donors were sampled and 12 cultures (2 different growth conditions varying in organic acid addition) were set up. Out of 12 enrichments one culture (and its subculture) was positive for ammonia oxidation and after further passages yielded T1S. In the case of R2S, one culture out of 10 setups (2 donor individuals) was positive. For the enrichments of X2B and Z3A three individuals were sampled and 6 initial setups yielded two positive cultures after 2 and 2.5 months respectively. Sequences of 16S rRNA and *amoA* showed only few point mutations to the sequences of *Ca.* N. epidermidis.

Positive enrichments were routinely transferred (transfer volume 20% v/v) upon reaching nitrite concentrations of approximately 450 µM. After transferring subcultures R2S and T1S into a medium containing 10 µg/ml chloramphenicol, 10 µg/ml novobiocin and 10 µg/ml streptomycin twice, no bacterial contaminants were detected, leaving the cultures with yeast contaminants. Both Bacteria and fungi signatures were found in X2B and Z3A.

### Routine culture maintenance

Enrichments were routinely cultivated in sterile 30 ml polystyrene screw cap tubes containing 18 ml FWM supplemented with 0.6 mM NH4Cl, 2 mM Na2CO3 (R) or NaHCO3 (T1S, X2B, or Z3A), 20 µl non-chelated trace element solution without molybdenum (Na2MoO4·2H2O), 7.5 µM FeNaEDTA, 20 µl vitamin solution as well as 25 µg/ml carbenicillin and kanamycin and were transferred every two to four weeks upon reaching nitrite concentrations of approximately 450 µM. The inoculation volumes were decreased to 10% (v/v).

The medium of enrichment R2S additionally contained 10 mM MOPS ((3-(N-morpholino) propanesulfonic acid), titrated to pH 7 with HCl) and had a final pH of 7. Ammonium and nitrite concentrations were measured colorimetrically by the indophenol method and Griess reagent respectively, as described previously (Reyes et al., 2020; Tourna et al., 2011). Aliquots used for ammonium measurements were frozen at -20°C and measured jointly at a later time point. Cultures were incubated aerobically at 28°C without shaking in the dark. Solutions were autoclaved (FWM, NH4Cl) or filter sterilized with a 0.2 µm filter (trace element solution, vitamin solution, FeNaEDTA, antibiotics) prior to use (Supplementary Fig. 1: Enrichment Scheme).

### Microscopy

Cultures were 10x concentrated before being imaged with phase contrast light microscopy using 60x and 100x oil immersion objectives. F420 autofluorescence was imaged using an F420 filter cube. For scanning electron microscopy six ml of each enrichment at late exponential phase were concentrated by centrifugation (21000 g, 4°C, 30 min). Pellets were fixed in 2.5% glutaraldehyde for 2 h, washed three times with 1x PBS and resuspended in 100 µl 1x PBS. 50 µl of the fixed cell suspension were spotted onto 0.01% poly-L-Lysine coated glass slides (5 mm) and sedimented for 2 h, before being dehydrated in an ethanol series (30-100%, 5 min each). The dehydrated cells were critical point dried in liquid carbon dioxide and subsequently coated with gold in a sputter coater. Samples were imaged in a JEOL IT 300 scanning electron microscope at 20 kV (CIUS imaging facility, Univ. of Vienna).

### DNA extraction of cultures

High molecular weight (HMW) DNA of enrichment cultures was extracted by a combination of chemical lysis based on a modified lysis buffer for *Sulfolobus acidocaldarius* (Meulenbroek et al., 2013) and freeze-thawing, followed by a phenol chloroform extraction.

Between 0.3 to 0.5 L of each skin enrichment culture were concentrated by filtration (0.22 µm MF-Millipore™ mixed cellulose ester (MCE) membrane filter), washed off the filter with FWM and pelleted by centrifugation (21000 g, 4°C, 30 min). Pellets were resuspended in a modified TENST buffer (20 mM Tris pH 8, 1 mM EDTA, 100 mM NaCl, 5% N-Lauroylsarcosine, 0.12% Triton X-100) and incubated at 50°C for 1-2 hours. Samples were subsequently subjected to three rounds of freeze-thawing (5 min at -70°C followed by 5 min at 70°C). Proteinase K was added to a final concentration of 100 µg/ml and samples were incubated at 65°C for 1-2 hours. After lysis HMW DNA was extracted by a standard phenol chloroform extraction (2x Phenol:Chloroform:Isoamylalcohol 25:24:1, 2x Chloroform:Isoamylalcohol 24:1) with low-speed centrifugation (4500 g, 4°C, 3 min) between steps. DNA was precipitated overnight at 4°C by adding 1µl glycogen as a carrier and 2x volume of a PEG6000 solution (30% PEG6000 (v/v), 1.6 M NaCl). The precipitated DNA was washed twice with 4°C cold 70% EtOH, air-dried at room temperature for 15-30 min, resuspended in 50-60°C nuclease free water followed by an incubation at 55°C for up to two hours. The extracted HMW DNA was quantified fluorometrically with a Qubit™ 2.0 Fluorometer (Invitrogen) and stored at -20°C.

### Quantitative PCR

Archaeal *amo*A genes were quantified in triplicate using primers *amoA*19F and *amoA*629R as previously (Abby et al., 2018; Alves et al., 2019) with some modifications: 20 µl reactions contained 10 µl Luna® Universal qPCR Master Mix, 3 µl nuclease free water, 0.5 µm of each primer and 5 µl DNA template. DNA was amplified in a CFX Real Time PCR System (Bio-Rad) with following cycling parameters: 95°C for 1 min, 40x cycles of 95°C 15 s denaturation, 60°C 30 s joint annealing-extension and 60°C 30 s fluorescence measurement. Standard dilutions prepared from DNA of each enrichment respectively using primers *amoA*19F and *amoA*629R ranged between 10^2^ and 10^8^ copies µl ^-1^. The efficiency of qPCR assays was between 86.06 and 93.04% with an R^2^ of ≥ 0.996.

### Sanger and NGS sequencing of marker genes and genomes

In order to confirm purity and identity of cultures, 16S rRNA genes were amplified by PCR with the archaea specific primer A109f (Großkopf et al., 1998), the universal primer Univ1492R (Lane, 1991), the crenarchaea specific primer pair 771F/957R (Ochsenreiter et al., 2003) and the fungal ITS region-specific primer pair ITS1F and TW13 (Taylor and Bruns, 1999). Amplification of the *amoA* genes was performed by PCR using the primers *amoA*19F (Tourna et al., 2008) and *amoA*629R (Arce et al., 2018). PCR products were purified using the Monarch® PCR & DNA Cleanup Kit from New England Biolabs before being sequenced.

### Genome sequencing, assembly and comparative genomics

DNA of our enrichments was shotgun sequenced using the NovaSeq 6000 (paired-end, 150 bp) platform at Novogene. Reads were trimmed and Illumina adapters were removed using Trimmomatic (SLIDINGWINDOW:5:20 LEADING:5 TRAILING:5 MINLEN:50 HEADCROP:6) (Bolger et al., 2014). To obtain the full genomic sequences of the enriched archaeal communities from the selected cultures, Nanopore sequencing on MinION Mk1C (Oxford Nanopore Technologies plc., UK) was performed. For library preparation, extracted genomic DNA was repaired with the NEBNext Companion Module (New England Biolabs GmbH, GER), and then prepared for sequencing on a chemistry version 14 flow cell (R10.4.1, FLO-MIN114) following the Ligation sequencing gDNA – Native Barcoding Kit 24 V14 (SQK-NBD114.24) and flow cell (FLO-MIN114) protocols.

Nanopore sequencing was performed according to the following configurations: run-time of the MinION Mk1C device was set for 72 hours, with a pore scan frequency of 1.5 hours, a minimum read length of 200 bp, high-accuracy base calling, and active channel selection, reserved pores, read splitting, trimming barcodes and mid-read barcode filtering enabled. The following software versions were used: MinKNOW v22.10.7, Bream v7.3.5, Configuration v5.3.8, Guppy v6.3.9 and MinKNOW Core v5.3.1. Resulting sequencing data was analyzed with the following specifications: First, simplex base calling was carried out on a GPU node at the Life Science Compute Cluster (LiSC) University of Vienna (Austria), using guppy-gpu v6.4.2 and the dna_r10.4.1_e8.2_260bps_sup.cfg model for superior base calling. Secondly, duplex base calling was achieved at a GPU Node at the LiSC University of Vienna (Austria), following the recommended tutorial on: https://github.com/nanoporetech/duplex-tools using dorado v0.1.1 (https://github.com/nanoporetech/dorado) with the dna_r10.4.1_e8.2_260bps_fast@v4.0.0 fast model, followed by finding duplex pairs with duplextools v0.2.20-3.10 (https://github.com/nanoporetech/duplex-tools) and dorado stereo/duplex base calling with the dna_r10.4.1_e8.2_260bps_sup@v4.0.0 superior model. Merged simplex and duplex reads were then investigated by performing quality control using NanoPlot nanocomp v1.20.0 (De Coster et al., 2018) followed by filtering with filtlong v0.2.1 (https://github.com/rrwick/Filtlong) (--min-length 200; --keep_percent 90, and --target_bases 20 million). Filtered reads were then assembled using flye v2.9.1 in --nano_hq and --meta mode with a target genome size of 3 Mbpm (Kolmogorov et al., 2020). Reads were mapped with minimap2 v2.24 (Li, 2021), and then polished with racon v1.5.0 (https://github.com/isovic/racon) using the following settings (-m -8 -x -6 -g -8 -w -500). Finally, consensus contigs were obtained with medaka v1.7.2 according to the r1041_e82_260bps_sup_g632 model (https://github.com/nanoporetech/medaka). Basic genome statistics were calculated with genometools by calling gt seqstat (Gremme et al., 2013), and classified the genomes using gtdbtk v2.1.1 with the --full_tree option enabled (Chaumeil et al., 2020). Genome completeness and contamination were estimated with checkm2 v1.0.1 including the --allmodels option (Chklovski et al., 2022). After genome comparison using drep v3.4.2 (Olm et al., 2017), genomes were finally annotated with eggnog-mapper v2.1.9 (Buchfink et al., 2015; Cantalapiedra et al., 2021; Eddy, 2011; Huerta-Cepas et al., 2019; Hyatt et al., 2010; Steinegger and Söding, 2017) including prodigal for gene identification. Marker genes were identified with Metaxa2 2.2 (Bengtsson-Palme et al., 2013) and the remaining contaminants of enrichments were estimated with kraken2 2.1.2 and bracken 2.8 according to the PlusPFP database (Lu and Salzberg, 2020; Wood et al., 2019) (https://benlangmead.github.io/aws-indexes/k2).

Gene prediction and annotation of the MAGs assembled in this study was performed automatically using the Microscope annotation platform from Genoscope (Vallenet et al., 2020), followed by extensive manual curation. Annotations were supplemented with arCOG assignments from the archaeal Clusters of Orthologous Genes database (2018 release) (Makarova et al., 2015), Carbohydrate-active enzymes assignments (CAZymes) from dbCAN2 (v7.0) (Zhang et al., 2018) and TCDB family assignments from the Transporter Classification Database (Saier et al., 2016), using scripts from (Dombrowski et al., 2020).

Orthologous protein families from a dataset including *Ca*. N. franklandianus C13, *Ca.* N. oleophilus MY3, *Ca.* N. hydrocola G61, *Ca.* N. agrestis SS, and *Ca.* N. arcticus Kfb and the two newly sequenced *Ca.* Nitrosocosmicus isolates R2S and T1S were constructed using Orthofinder (v.2.5.4) with standard settings (Emms and Kelly, 2019).

### Study-cohorts for molecular-based analyses

Forty-seven healthy individuals (group A1: female=11, male=10, age=20-40 years; group A2: f=15, m=11, age=60-85 years) were recruited for an in-depth analysis of their skin microbiome, and a subgroup of high vs. low skin archaeal carriers was selected based on quantitative (*amoA* gene and 16S rRNA gene qPCR universal and archaea specific primers; data types: Cquant1, Cquant2, and Cquant3; Supplementary Fig. 2) and qualitative (16S rRNA gene amplicons universal and archaea specific primers; data types: Cqual1, Cqual2, and Cqual4; Supplementary Fig. 2) observations for longitudinal investigations (group B1 (from A1): f=2, m=4; group B2 (from A2): f=5, m=1) over the period of almost two years (6 time points; data type: L). The sampling sites for A1/A2 were hand, outer arm, crook, armpit, forehead, part, décolleté and back (n=7), and for B1/B2: forehead, décolleté, arms and back (n=4). Supplementary Table S3 gives an overview on the demographic characteristics of all recruited subjects.

Each subject completed a questionnaire with different categories, e.g., height and weight, skin type, and detailed questions about their respective lifestyle (e.g., eating habits, smoking, alcohol consumption, skin care, medications, animal contact, and hobbies).

### Skin measurements and sampling

The forehead and the part, as well as both hands, forearms, crooks and armpits were swabbed with pre-moistened (0.9 % (v/v) NaCl) swabs (one swab for each body site; BD BBL ™ Culture Swab ™ EZ; Becton Dickinson, USA). NaCl was baked at 260°C for 24 h before solving in PCR-grade water to remove DNA traces. The remaining body sites, including décolleté and back (approximately 300 cm^2^), were sampled with a pre-moistened surface-sampling cellulose-sponge kit (VWR, 300-0229P). For the longitudinal sampling (cohort B1/B2, tp2-6), only the forehead was sampled with the swab. After skin sampling, samples were immediately frozen at -80°C until further processing. All skin sampling events were accomplished in a lab that never handled AOA, and controls from the environment, sampling procedure, and sample extraction were processed in parallel.

Measurements of the skin physiology was performed using the Cutometer(R) MPA580 (Courage + Khazaka, Germany), using the Tewameter (to determine the barrier function by measuring the water loss; 30s, 3 times), the pH-meter, the Sebumeter (to determine the sebum-content, 30s), and the Corneometer (to determine skin moisture). Each skin site was measured three times and the mean of the measurements is reported in Supplementary Tables S4 and S5. Further, the surface temperature of each region was determined using an infrared thermometer (Future Founder, China).

The interpretation of skin parameters was done based on the recommended specifications by the manufacturer, which are provided in Supplementary Table S6.

### DNA extraction, controls, amplicon sequencing, and qPCR of skin samples

The DNA was extracted from swabs and sponges using the Purelink ™ Microbiome DNA Purification Kit (Rectal or Environmental Swab Samples; Thermo Fisher Scientific, USA). For the extraction from sponges (liquid sample extract from décolleté and back) one ml of the sample was concentrated by centrifugation (20817 x g, 4°C; 10 min). The largest proportion of the supernatant was removed, and the remaining 70 µl was used for bead beating with 800 µl of the S1 lysis buffer. Material and process controls were included in each step to control for the DNA contamination of the used reagents. The DNA was stored at -20°C until further downstream analyses.

Various targets were assessed using amplicon-based analyses: i) the archaeal 16S rRNA gene region (nested PCR approach as described in (Pausan et al., 2019) and (Probst et al., 2013)), the ii) *amoA* targeted approach (described in (Arce et al., 2018; Leininger et al., 2006; Tourna et al., 2011)), and the iii) “universal” approach, targeting the microbial 16S rRNA gene region (Walters et al., 2011). All primer sequences and PCR protocols are provided in Supplementary Table S7.

Library construction and next-generation sequencing (Illumina MiSeq®) was performed at the Core Facility Molecular Biology at the Center for Medical Research (Medical University of Graz, Austria). In a first step, the DNA concentrations of the generated amplicons were normalized using a SequalPrepTM normalization plate (Invitrogen) and subsequently each sample was indexed with a unique barcode sequence by 8 cycles of indexing PCR. These indexed Samples were pooled and purified by gel cuts before the library was run on an Illumina MiSeq® instrument and the MiSeq® Reagent Kit v3 with 602 cycles (2 x 301 cycles).

### Bioinformatics and data analysis

Resulting demultiplexed fastq reads were processed with QIIME2 (Bolyen et al., 2018) versions 2018.11 through 2023.2. After quality control, denoising of reads was achieved with DADA2 (Callahan et al., 2016) and truncation of forward reads at a length of 230bp and 120bp for reverse reads, was only used for the archaea-specific amplicon (519F-806R), while no truncation was set for all other amplicon constructs (universal: 515F-926R; and *amoA*: 19F-629R). Resulting feature tables and representative sequences were deposited on our Github repository (https://github.com/CME-lab-research/SkinArchaeome). Potential contaminants were identified with decontam v.1.12 (Davis et al., 2018) using the prevalence mode with a threshold of 0.5 and removed from downstream analysis. Representative sequences were classified by a pre-trained Naïve-Bayes classifier based on the curated and trimmed SILVA138 database (Klindworth et al., 2013) using RESCRIPt (Robeson et al., 2021) and the feature-classifier classify-sklearn plugin (Bokulich et al., 2018). For phylogenetic metrics like UniFrac (Lozupone and Knight, 2005), a rooted phylogenetic tree was generated with FastTree2 (Price et al., 2010) based on a masked MAFFT alignment. Data normalization was achieved by scaling with ranked subsampling and the q2-srs QIIME2 plugin (Heidrich et al., 2021) at Cmin 50, 100 and 1000 followed by calculating core metrics for alpha and beta diversity. Feature differentials were determined with the QIIME2 plugins aldex2 (Fernandes et al., 2013), ANCOM (Lin and Peddada, 2020), ANCOM-BC (Lin and Peddada, 2020), and the R packages ANCOM2 (https://github.com/FrederickHuangLin/ANCOM-Code-Archive) and MaAsLin2 (Mallick et al., 2021) including time as a fixed effect and different subjects as a random effect where applicable, while feature loadings were determined with the QIIME2 plugin DEICODE (Martino et al., 2019). Longitudinal analysis covering feature volatility over time as well as other numerical metadata categories like age, BMI or skin physiology measurements including i.e. pH, skin fat or skin wetness were realized with the q2-longitudinal plugin in QIIME2 (Bokulich et al., 2018). The same plugin was used to calculate linear-mixed-effect models of Shannon diversity and weighted UniFrac distances along principal coordinate axis 1. Amplicons of the *amoA* marker gene were processed in a similar way without read truncations for DADA2 and read classifications by a pre-trained Naïve-Bayes classifier based on the *amoA* reference database (Alves et al., 2018; Wang et al., 2021). Connection of 16S rRNA to *amoA* gene information was achieved by following the tutorial at https://github.com/alex-bagnoud/Linking-16S-to-amoA-taxonomy/. Prediction of metadata categories were realized with the q2-sample-classifier QIIME2 plugin (Bokulich et al., 2018).

#### Custom python code for regressing all metadata categories

To identify potential associations between taxa and recorded numerical and categorical metadata an ordinary least squares (OLS) regression fit method was used with python’s statsmodels smf formula (<u>statsmodels.regression.linear_model.OLS -statsmodels 0.15.0</u> <u>(+8)</u>). Microbial counts of the different investigated cohorts were first clr transformed and filtered for less than 25% missing values. These values were then used as the response variable in dependence of each individual, their sex, age, BMI, body site, the overall microbial diversity indicated by the Shannon index (H’), and where applicable the time of sampling. The model was then evaluated for each genus and recorded metadata according to their q-values (FDR-corrected p-values) and estimated regression coefficients (python code available at GitHub: https://github.com/CME-lab-research/SkinArchaeome).

#### Network analysis

Network analyses were conducted to identify mutually exclusive and co-occurring clusters of archaea and bacteria in our merged (archaea-specific and universal) 16S rRNA gene amplicon dataset. After SRS normalization to Cmin = 100 and TSS and log transformations the 40 most abundant taxa were represented as nodes within the network where node size reflects taxa abundance, and edges indicate only positive associations of Spearman correlations between pairs of taxa. Then, we used PCoA to ordinate the nodes in a two-dimensional plot, such that correlating nodes were close together and anti-correlating nodes were far apart. In addition, correlations of the network were further supported by the Sparse Co-Occurrence Network Investigation for Compositional data (SCNIC) package (https://github.com/lozuponelab/SCNIC) using the sparCC metric with default settings for filterings and the R value, and bootstrap adjusted p-values.

#### Phylogenomic tree reconstruction

A genome database was created comprising the two AOA genomes presented in this study together with 141 MAGs and completely sequenced genomes (133 AOA and 8 non-AOA genomes) from IMG, DDBJ, or NCBI databases followed by protein prediction using Prodigal v2.6.3 (Hyatt et al., 2010). The workflow proposed by Graham et al. (2018) (Graham et al., 2018) based on the archaeal single-copy gene collection (Rinke et al., 2013) was followed to identify phylogenetic markers (e-value 10^-10^) to reconstruct the phylogenomic tree. A total of 36 ribosomal protein families encoded in at least 100 out of the 143 genomes present in our genome database were selected. Each protein family was aligned independently using MAFFT 7.520 (Katoh and Standley, 2013) (L-INS-i algorithm) followed by the trimming of the alignments using BMGE (Criscuolo and Gribaldo, 2010) with default parameters. Trimmed protein families were concatenated using a custom python script and the concatenated alignment was used to reconstruct a maximum likelihood (ML) phylogenomic tree in IQ-TREE v2.2.2.7 (Minh et al., 2020) under the LG+C20+F+G model with 1000 ultrafast bootstrap replicates.

#### Phylogenetic analysis of 16S rRNA sequences

The identification of 16S rRNA sequences in our AOA genome database (see above) was performed using Barrnap v0.9 (https://github.com/tseemann/barrnap). The 16S rRNA sequence encoded in the MAG GCA_025930575.1 (NS-Epsilon) was used as a query in a BLASTN search against the NCBI database to improve the sequence representation of this sub lineage with environmental sequences. Sequences were aligned with the SINA v1.7.2 aligner (Pruesse et al., 2012) using the SILVA database SSU Ref NR 99 release 138.1 (Klindworth et al., 2013) with default parameters followed by the alignment of the skin ASVs using MAFFT 7.520 (Katoh and Standley, 2013) (--addfragments option, L-INS-i algorithm). The alignment was trimmed manually using AliView (Larsson, 2014). The maximum likelihood phylogenetic tree was reconstructed using IQ-TREE v2.2.2.7 (Minh et al., 2020) with the best-fit model TIM2e+I+R3 chosen by ModelFinder (“-m MFP”) (Kalyaanamoorthy et al., 2017) according to BIC (Bayesian Information criterion), SH-like approximate likelihood ratio test (“-alrt 1000”) (Guindon et al., 2010) and 1,000 ultrafast bootstrap replicates (“-bb 1000”).

#### qPCR specific methods

A SYBR green-based quantitative real-time PCR (qPCR) approach targeting the archaeal *amoA* gene (19F-629R) and the 16S rRNA gene of archaea (A806F-A958R) and bacteria (Bac_331F-Bac_797R) was used to determine the abundance of AOA, all archaea and all bacteria on selected body sites. Primers, reaction mixtures and thermocycler programs are listed in Supplementary Table S17. All qPCRs followed the MIQE Guidelines.

qPCRs were performed with the Bio-Rad SsoAdvanced™ Universal SYBR® Green Supermix. Each reaction contained 1µl of DNA template and 9µl of mastermix. The mastermix for the archaeal *amoA* approach consisted of 5µl SsoAdvanced™ Universal SYBR® Green Supermix (Bio-Rad), 1.25µl of each primer (1.25µM) and 1.5µl of PCR grade water. The mastermix for the archaeal and bacterial 16S rRNA gene approach consisted of 5µl SsoAdvanced™ Universal SYBR® Green Supermix (Bio-Rad), 0.3µl of each primer (10µM) and 3.4µl of PCR grade water. Amplifications were carried out in a Bio-Rad CFX96 Touch™ Real-Time PCR Detection System. The qPCR cycling conditions were as follows: initial denaturation at 95°C for 1.5min, followed by 45 cycles of denaturation for 15sec at 95°C, annealing for 30sec at 60°C, elongation for 40sec at 72°C, a final denaturation for 15sec at 95°C, 5sec at 60°C and a storage temperature of 8°C. A plasmid containing the *amoA* gene of *Nitrososphaera viennensis* with a starting concentration of 104.1ng/µl (pscA_Ampkan_AmoA, clone 3) was used as a quantification standard in a 1:10 dilution series from 10^-2^ to 10^-10^. Product specificity was confirmed by melting curve analysis. All qPCR experiments were performed in triplicates and the results were averaged for further calculations. Data from a previous experiment on the total archaeal community (based on qPCR of the 16S rRNA gene) was also included in the analyses. To account for the different area sizes of the sampled body sites, copy numbers were adjusted to 300 cm^2^ (*amoA* gene copies per 300 cm^2^; see Supplementary Table S8). The qPCR files were analyzed using the CFX Manager™ Software, version 3.1 and processed with the R software, version 3.6.1 and Microsoft Excel.

## Results

### Cultivation of AOA from human skin

Different types of inocula were used to initiate enrichment cultures under oxic conditions in the presence of 0.5 mM ammonia as sole energy source. Pooled samples were used after swabbing the face, neck, upper body, back, arms, and legs of different individuals. After 2 to 9 months, amplification and sequencing of archaeal 16S rRNA and *amoA* gene fragments identified the presence of single AOA strains in initial enrichments. The usage of pooled samples of three different individuals resulted in three enrichment cultures: *Ca.* Nitrosocosmicus “T1S”, “X2B”, and “Z3A”. A fourth enrichment culture was obtained from cut fingernail edges as initial inoculum (“R2S”) (details of enrichment source are provided in Supplementary Table S2).

The current study focuses on two of the enrichment cultures, “T1S” and “R2S”, isolated from skin swabs and fingernail clippings respectively. These enrichments exhibited high enrichment of 88-98% based on microscopy and molecular techniques (Supplementary Fig. 3) and stoichiometric conversion of ammonia to nitrite every 3-4 weeks (Fig. 1 A-B) after repeated transfers over the course of more than a year in medium supplemented with 0.6 mM ammonium, 2 mM of either sodium carbonate (R2S) or sodium bicarbonate (T1S) and an antibiotic mix. No bacterial contaminants were detected via amplification of the bacterial 16S rRNA gene (Supplementary Fig. 4), or by light and scanning electron microscopy (Supplementary Fig. 3). Based on the sequenced internal transcribed spacer (ITS) region the remaining contaminant in both enrichments was a single fungal species with 99% sequence identity to *Exophiala* sp. DAOM 216391, a genus of fungi frequently found in manmade environments and known to be an opportunistic pathogen of humans (Untereiner and Naveau, 1999; Usuda et al., 2021).

**Fig. 1.**
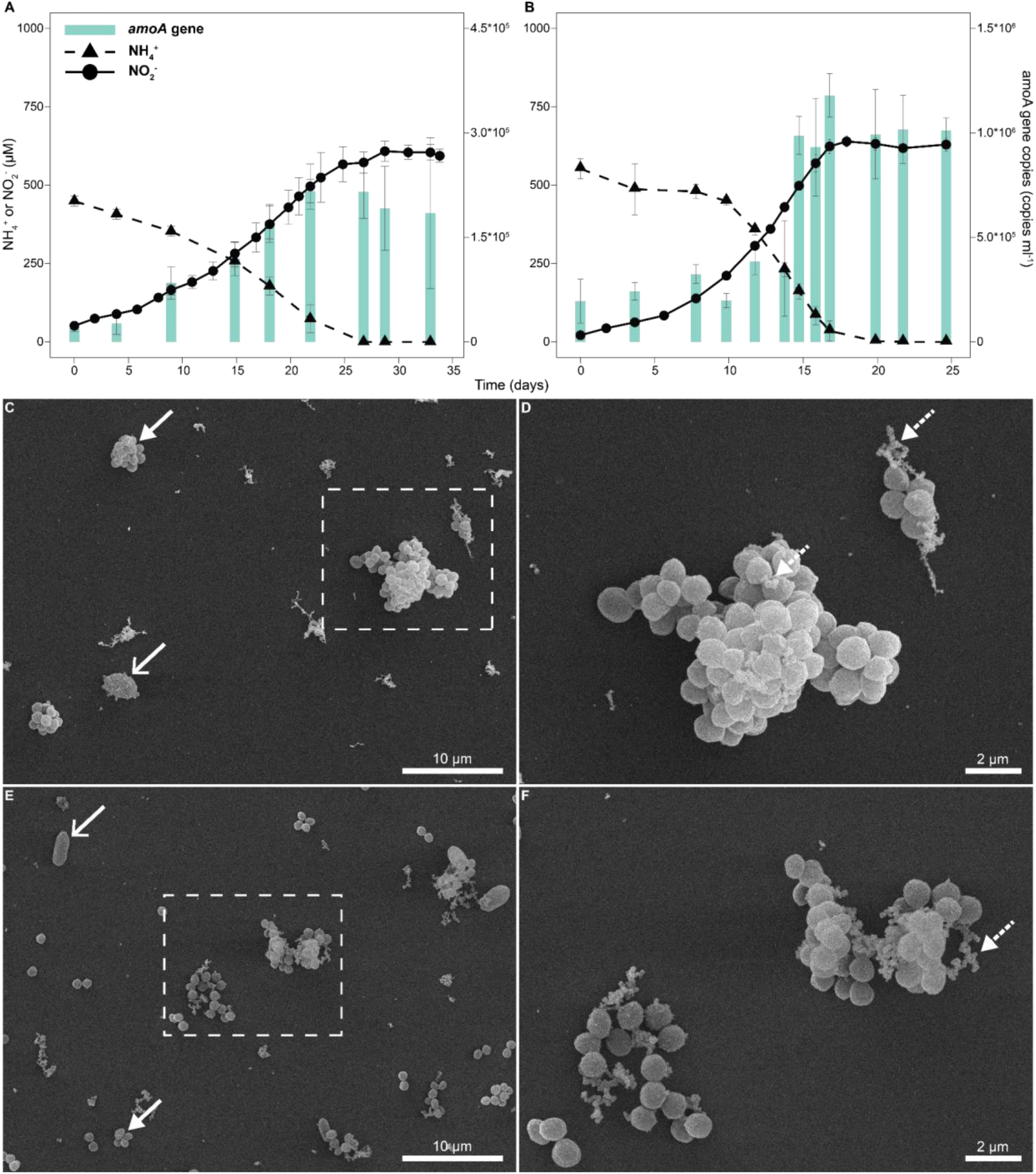
Growth and morphology of skin enrichments R2S (A) and T1S (B) displaying near stoichiometric conversion of ammonia (triangles, dashed line) to nitrite (circles, solid line) paralleled by cell growth, estimated by qPCR of archaeal *amoA* gene copies (orange bars). Ammonia and nitrite measurements are averages of biological triplicates. Gene copies of *amoA* are averages of technical triplicates of each biological triplicate. Error bars depict the standard deviation of the mean. Scanning electron micrographs of skin enrichments R2S (C, D) and T1S (E, F) with white arrows indicating AOA cells based on the morphologies seen in Supplementary Fig. 3 and already published members of the *Ca.* Nitrosocosmicus (Alves et al., 2019; Jung et al., 2016; Lehtovirta-Morley et al., 2016; Liu et al., 2021). Dashed white arrows indicate putative extracellular polymeric substances (D, F). Black arrows indicate the remaining fungal contaminant.

Cell growth of the two strains R2S and T1S correlated with nitrite production as determined by quantification of *amoA* genes using qPCR, as has previously been shown for other AOA (Fig. 1, A-B) (Abby et al., 2018; Alves et al., 2019; Tourna et al., 2011). The shortest generation times based on nitrite production were approximately five days at 28°C. After the oxidation of ∼500 µM ammonium enrichments R2S and T1S reached cell densities of 2.16*10^5^ (± 1.68*10^4^) and 1.18*10^6^ (± 1.05*10^5^) cells ml^-1^, respectively. This is similar to the cell densities of *Ca.* Nitrosocosmicus franklandianus C13 (3.8*10^6^ cells ml^-1^) (Lehtovirta-Morley et al., 2016) and *Ca.* Nitrosocosmicus oleophilus MY3 (1.6*10^6^ cells ml^-1^) (Jung et al., 2016) after oxidation of ∼500 µM ammonium.

For light microscopy, putative AOA were observed as cocci with diameters between 1-2 µm which frequently formed aggregates and were partly enclosed in putative extracellular polymeric substances (EPS) (Supplementary Fig. 3). These morphological features are comparable with those of previously published members of the genus *Ca.* Nitrosocosmicus (Alves et al., 2019; Jung et al., 2016; Lehtovirta-Morley et al., 2016; Liu et al., 2021) and were confirmed by scanning electron microscopy (Fig. 1, C-F).

### Taxonomic placement and genomic features of the AOA representatives from human skin

Sequencing and assembly of the genomes of the two skin enrichments resulted in one circular contiguous sequence affiliated with AOA for each strain. The genome sizes, G+C content and predicted protein-coding genes of the strains were well within the range of other representatives of the genus *Ca.* Nitrosocosmicus (Table 1).

Like other representatives of this genus, both strains encoded three identical copies of the 16S and 23S rRNA gene operons. The average nucleotide identity (ANI) and average amino acid identity (AAI) values between strains R2S and T1S (87.4% ANI, 86.2% AAI), and between them and other *Ca.* Nitrosocosmicus genomes and proteomes were well below the threshold of 95% ANI/AAI proposed for species delineation (Konstantinidis et al., 2017) (Supplementary Tables S9 and S10). We therefore propose that they represent two distinct novel species, preliminarily named *Ca.* Nitrosocosmicus epidermidis strain T1S (e.pi.der’mi.dis. Gr. fem. n. epidermis, the outer skin; N.L. gen. n. epidermidis, of the epidermis) and *Ca.* Nitrosocosmicus unguis R2S (un’gu.is. L. gen. n. unguis, of a fingernail).

A maximum likelihood analysis based on 36 concatenated ribosomal protein marker genes present in one copy in at least 100 of the 143 genomes of a dataset comprising representative AOA and non-AOA of *Nitrososphaeria* confirmed the affiliation of the two novel strains with the NS-zeta lineage as defined by Alves et al. 2018 (Alves et al., 2018). Specifically, the two strains formed a highly supported cluster with *Ca.* N. arcticus (obtained from Arctic soil), *Ca.* N. oleophilus (from coal-tar contaminated sediment), and other uncharacterized MAGs (Fig 2, and Supplementary Table S9-S11).

**Fig. 2.**
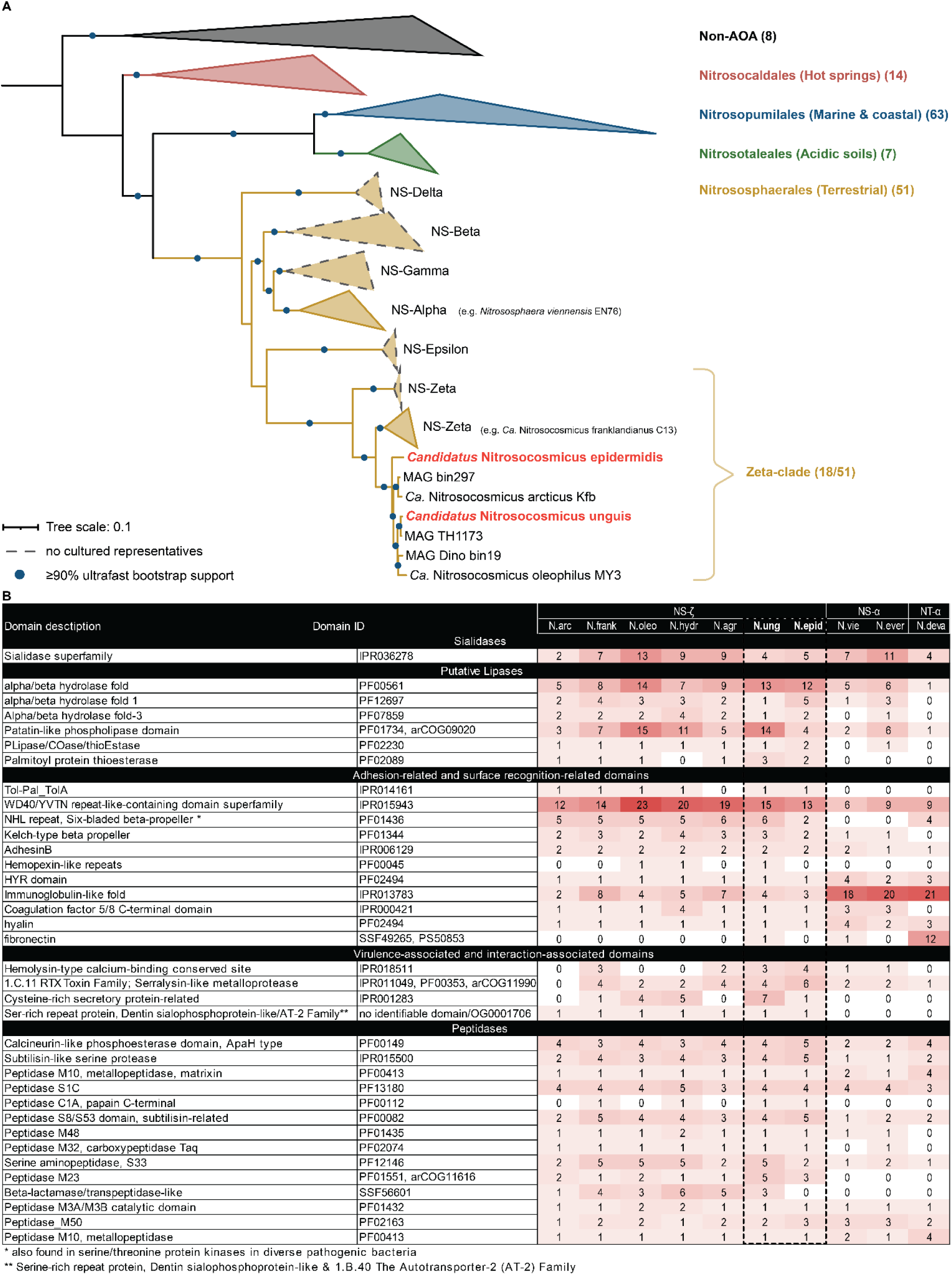
(A) Maximum-likelihood phylogenomic tree of AOA and non-AOA reconstructed based on the concatenated alignment of 36 ribosomal proteins comprising a total of 4,707 amino acid sites. Numbers in parentheses represent the number of complete genomes and MAGs in each lineage. NS = Nitrososphaerales. (B) Heatmap displaying the detection frequency of specific functional domains discussed in the text in all analyzed *Ca.* Nitrosocosmicus genomes. Numbers refer to the number of occurrences of the respective domain in the genome, but do not always correspond do distinct CDS (i.e. multiple domains on one CDS). Domain IDs refer to Interpro, Pfam. TIGRfam, Superfamily Database entries.

To investigate the specific genomic repertoire of *Ca.* N. unguis and *Ca.* N. epidermidis, we compiled a dataset consisting of the predicted proteomes of all available genomes from cultivated strains of the genus *Ca.* Nitrosocosmicus: *Ca*. N. franklandianus C13, *Ca.* N. oleophilus MY3, *Ca.* N. hydrocola G61, *Ca.* N. agrestis SS, and *Ca.* N. arcticus Kfb (we refrain from the “Ca.” prefix for better readability in the following). This protein dataset was clustered into 3,301 orthologous protein families using Orthofinder (see Methods, Supplementary Data: Orthologs, and Supplementary Table S12). From these orthologous protein families, 1,592 represented the core genome of the genus, comprising about 50% of the predicted CDS of the genomes (Supplementary Table S13). The NS-Zeta subclade containing N. epidermidis, N. arcticus, N. unguis and N. oleophilus shared 60 protein families N. unguis and N. epidermidis shared only 20 protein families to the exclusion of all others, while both of them shared more protein families with N. oleophilus (125 and 22 respectively) than any other *Ca.* Nitrosocosmicus genome. The shell (=strain-specific) proteome of each skin isolate consisted of 17 and 16 families for N. unguis and N. epidermidis respectively. The content of arCOG functional categories did not differ among the *Ca.* Nitrosocosmicus genomes, indicating that presence and survival on human skin did not require a particular expansion in a functional category (Supplementary Fig. 5).

### A genomic island and expanded gene families implicate factors involved in host interactions

The *Ca.* N. *epidermidis* genome contains genomic islands flanked by integrases, encoding proteins potentially involved in host interactions (Supplementary Data: Annotations). One such island (Nepid_2202–2261) includes a putative autotransporter-1 (AT-1) family protein (TCDB: (1.B.12), Nepid_2213) with extracellular domains of unknown function. These proteins typically use a C-terminal β-barrel to export N-terminal virulence factors (van Ulsen et al., 2018). In *Ca.* N. epidermidis, leucine-rich repeats on the N-terminal could facilitate protein– protein recognition(Kobe, 2001). The island also encodes a putative α-arrestin-like protein (Nepid_2245), containing arrestin N-and C-terminal domains (IPR011021 and IPR014752), which in eukaryotes bind to and regulate 7TM and kinase receptors (e.g., GPCRs) (Alvarez, 2008). Though uncharacterized in prokaryotes, some pathogens exploit host β-arrestin pathways to traverse epithelial barriers (Marullo and Coureuil, 2014). Additionally, the island includes eight membrane-bound serine/threonine kinases and four serine-rich adhesins, known in streptococci for mediating tissue and extracellular matrix binding (Chan et al., 2020). In *Ca.* N. *unguis*, expanded gene families also suggest roles in host interaction (Fig. 2B, Supplementary Data). Notably, it contains seven cysteine-rich secretory (CAP) family proteins (vs. one in *Ca.* N. *epidermidis* and only a few in other *Ca.* Nitrosocosmicus genomes), a group associated in eukaryotes with adhesion, signaling, and ion channel regulation, and potentially linked to pathogenesis or antimicrobial defense (Gibbs et al., 2008; Yeats et al., 2003). *Ca.* N. *unguis* is also enriched in beta-propeller fold proteins, which may mediate environmental interactions through ligand binding, signaling, or enzymatic activity in the periplasmic space, and are more abundant in *Ca.* Nitrosocosmicus than in other AOA lineages (Chen et al., 2011).

### Archaeal ammonia oxidizers form a rare, yet prevalent population on human skin

A complementary, but independent approach was used to understand the abundance, prevalence, distribution and stability of archaeal, and particularly *Ca.* Nitrosocosmicus representatives on human skin. To achieve this, 47 healthy individuals were independently recruited and eight defined body locations were sampled (cohorts A1 (age 20-40 years), A2 (60-80+ years), study overview: Supplementary Fig. 2 STORM chart, data types C). According to their microbial profiles (high AOA, low AOA, see methods), 12 of these individuals were then selected and longitudinally sampled on four body sites for an entire year (cohort B; study overview: Supplementary Fig. 2, data types L). Samples were quantitatively (qPCR) analyzed using taxonomic (16S rRNA) and functional (*amoA*) marker genes, targeted towards the detection of archaea and archaeal ammonia oxidizers (AOA). As metagenomic sequencing failed due to the low biomass and high background in human DNA, diversity assessments of the skin archaeome were performed via amplicon sequencing to profile the bacterial, archaeal and AOA-specific signatures. A total of 1,770 amplicons (PCR reactions (16S rRNA gene, *amoA*) per sample including controls) were processed (details see Materials and Methods, Supplementary Fig. 2, and Supplementary Table S14).

Throughout the study (cohorts A, B), archaeal signatures were found at least once in all study participants, but were overall low, according to qPCR measurements (approx. 0.15±0.07% relative abundance, average across all body sites; data types Cquant1, Cquant2, Cquant3). AOA-specific qPCR signals accounted for 1.54*10^3^ gene copies per 300 cm^2^ (Fig. 3A). The highest amount of *amoA* gene copies were found in samples from the forehead (Fig. 3B); there, AOA signatures were found in samples from every tested person at each time point (prevalence: 100%; data types Lquant1, Lquant2, and Lquant3).

**Fig. 3.**
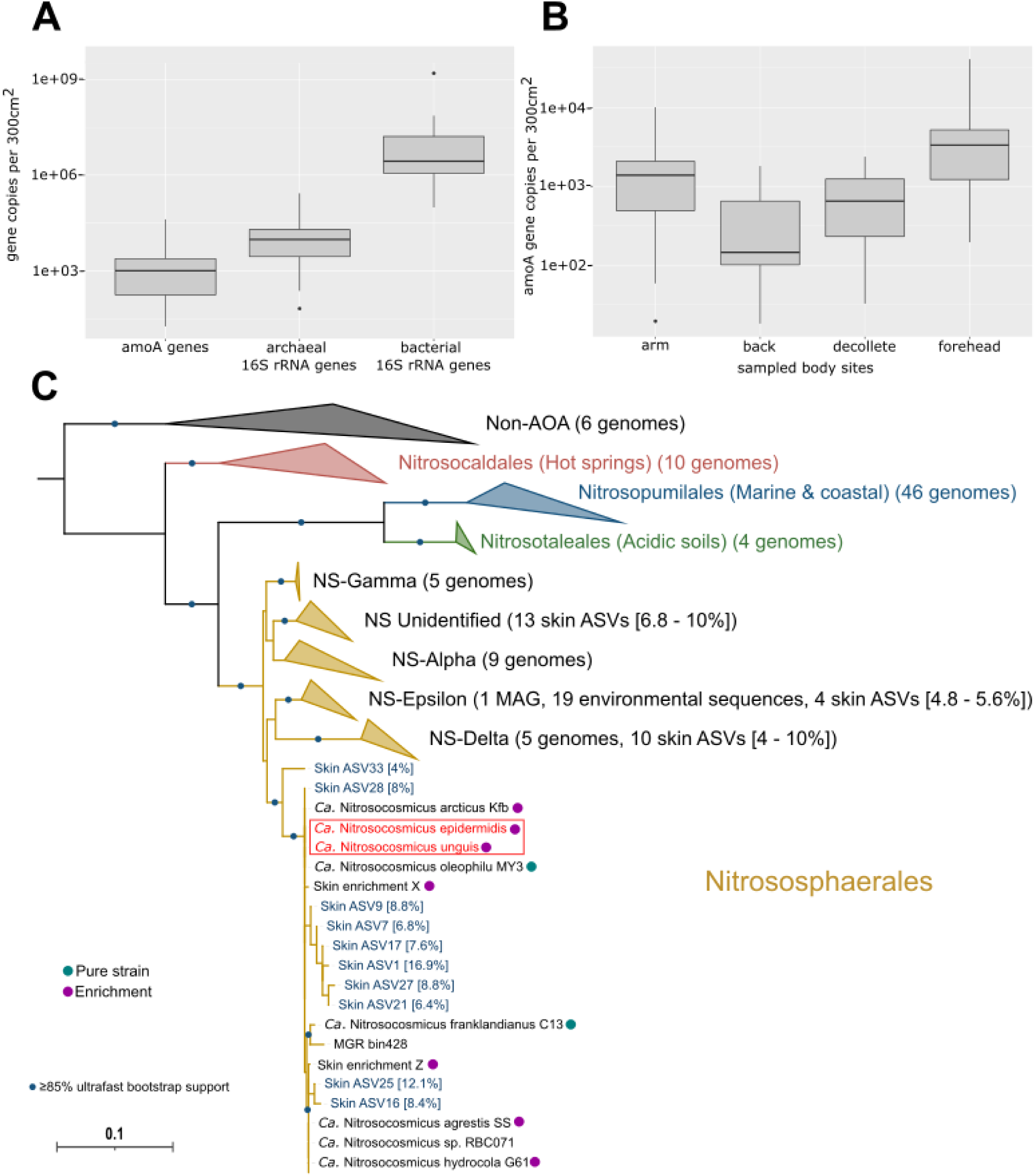
(A and B) Quantitative analysis (qPCR) of AOA, archaeal and bacterial 16S rRNA gene signatures on human skin (n=284; data types: Lquant1, Lquant2, and Lquant3; Supplementary Fig. 2). The median of 16S rRNA gene copies across all sampling sites is shown; data are corrected for different 16S rRNA gene copy numbers in archaea (median: 1) and bacteria (6); https://rrndb.umms.med.umich.edu/). Gene copies per 300 cm^2^ of AOA, Archaea and Bacteria, shown on a logarithmic scaled Y-axis. (B) arm vs. back (p = 5.9*10^-4^), arm vs. decollete (p = 7.2*10^-4^), arm vs. forehead (p = 2.1*10^-12^), back vs. forehead (p = 5.3*10^-13^), decollete vs. forehead (p = 6.1*10^-13^). *AmoA* gene copies per 300 cm^2^, across sampling sites. (C) Maximum-likelihood phylogenetic tree of 16S rRNA gene sequences of AOA and non-AOA, including most prevalent and abundant ASVs of cohort A and B. Blue sequences represent ASVs retrieved from amplicon sequencing of skin samples (cohorts A, B). Values in brackets show the percentage of prevalence of the ASV in our samples.

In amplicon sequencing, on average 6.88±4.67% of all samples contained archaea when the universal PCR primer combination was applied (Supplementary Table S15, data types: Cqual1 and Lqual1; Supplementary Fig. 2), whereas an average of 69.98±18.05% were positive with the archaea-targeted combination of 16S rRNA gene primers (Supplementary Table S8, data types: Cqual2 and Lqual2; Supplementary Fig. 2), emphasizing once more the necessity to use archaea-targeted methodology for optimized detection. Also, with this approach, all individuals were found to be positive for AOA signatures at least once throughout the study period (on average 86±25%; 1.1*10^5^±2.1*10^5^ reads of all samples).

Our independent approach with 16S rRNA gene sequencing detected most abundant AOA-associated ASVs within the *Ca.* Nitrosocosmicus genus which matched the identified genus from the cultivated enrichments (Fig. 3C). The most prevalent archaeal family within 326 samples (cohort A and B) was Nitrososphaeraceae (74% positive samples) with *Ca.* Nitrosocosmicus being the most prominent genus in this family (24%) (Supplementary Table S16; data types: Cqual2 and Lqual2; Supplementary Fig. 2). The joint classification of 16S rRNA to the *amoA* marker gene (see Methods for details) resulted in a prevalence of 38% for the NS-Zeta clade, across all samples (Supplementary Table S16; data types: Cqual2 and Lqual2; Supplementary Fig. 2).

### *Ca.* Nitrosocosmicus signatures show longitudinal coherence and unique biogeographic patterns across body sites

Across all eight body sites sampled (cohort A; data type: Cqual2; Supplementary Fig. 2), Nitrososphaeraceae and *Ca.* Nitrosocosmicus signatures were mostly detected on forehead, décolleté, back and arms, indicating a skin physiology-dependent pattern of ammonia-oxidizing archaea in general (Fig. 4, 16S rRNA gene amplicons). High proportions of AOA from the head region could be confirmed quantitatively (qPCR), as highest copy numbers of the *amoA* gene (up to 4.11*10^4^ per 300 cm^2^) were retrieved from the head samples (part and forehead; Fig. 3B; data type: Lquant3; Supplementary Fig. 2).

**Fig. 4.**
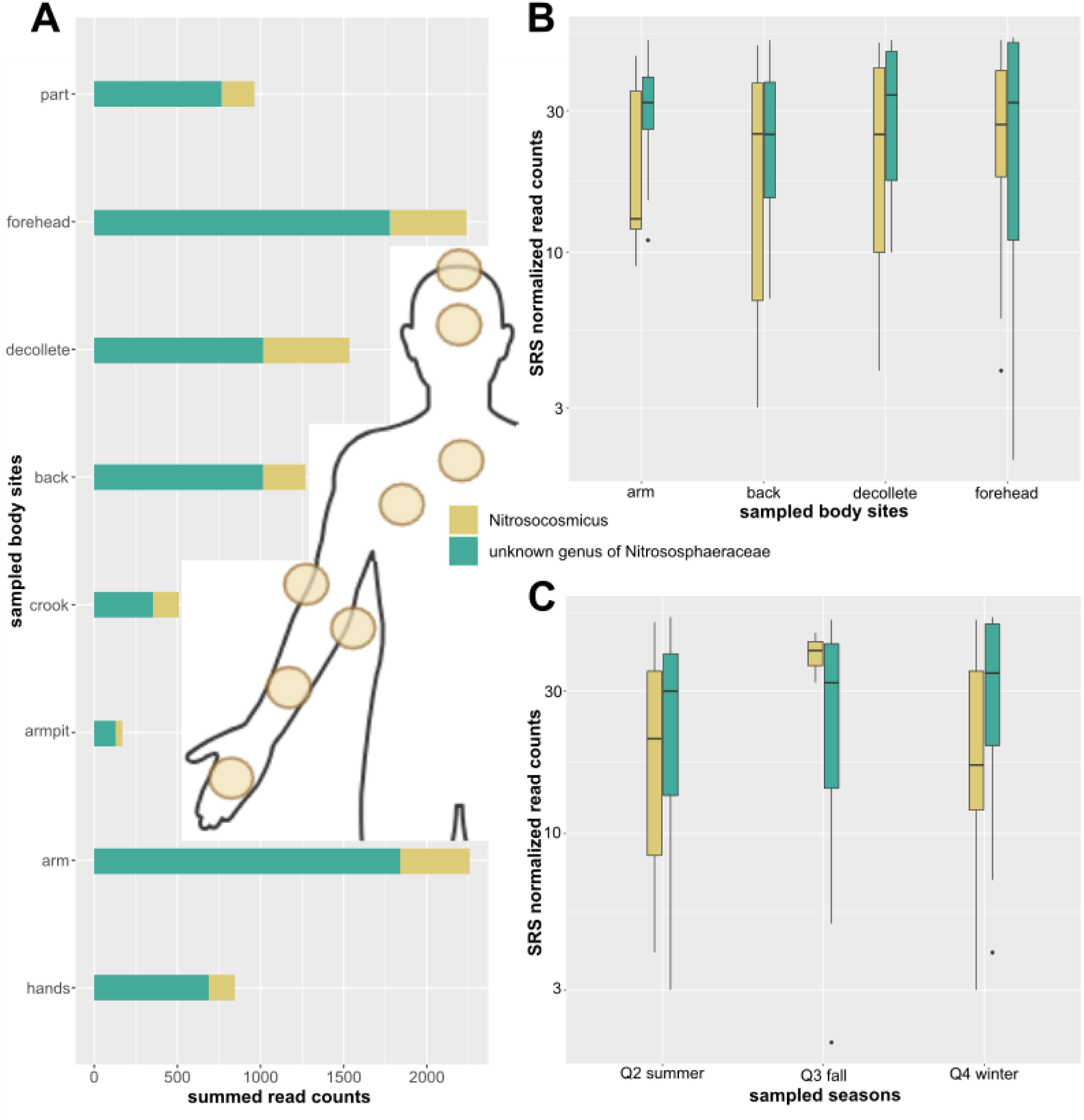
(A) AOA prevalence across all samples from different body sites based on 16S rRNA gene amplicon sequences, shown by summed read counts (n=326 samples; data types: Cqual2; Supplementary Fig. 2). The reads obtained were assigned to *Ca.* Nitrosocosmicus and other non-higher resolved genera belonging to the family Nitrososphaeraceae. Read counts and percentage of all sampled body sites for *Ca.* Nitrosocosmicus (part: 197, 9%; forehead: 461, 21%; décolleté: 519, 24%; back: 256, 12%; crook: 156, 7%; armpit: 41, 2%; arm: 418, 19%; hands: 157, 7%). Read counts and percentage of all sampled body sites for all AOA (part: 964, 10%; forehead: 2239, 23%; décolleté: 1535, 16%; back: 1271, 13%; crook: 509, 5%; armpit: 170, 2%; arm: 2258, 23%; hands: 846, 9%). (B) Boxplot showing the abundance of AOA (normalized read counts) per body site. (C) Boxplot showing AOA dynamics (normalized read counts) across the three covered seasons. Both B) and C) show read counts on a log-scaled Y-axis and refer to the data of the longitudinal cohort (1-year, 6 time points, 3 seasons, 4 body sites, 283 samples; data type: Lqual2; Supplementary Fig. 2).

*Ca.* Nitrosocosmicus signatures were stable longitudinally (1-year sampling period, Supplementary Information; data type: Lqual2; Supplementary Fig. 2), and even individuals that were not Archaea positive at the initial sampling time point became positive over the course of the study period. In addition, participants who carried archaea at the initial time point consistently showed archaeal signatures above the detection limit throughout all subsequent time points (on average 78±10%; 1.4*10^5^±9.4*10^4^ reads of all samples).

Seasons did not have a significant impact on the abundance of AOA (MaAsLin2, q-value = 0.95), indicating that contact with the outer environment, e.g. through contact to soil by gardening work (MaAsLin2, q-value = 0.49; data type: Lqual2; see Supplementary Fig. 2 STORM chart for more details), did not influence the skin archaeome. We then tried to identify AOAs on genus level that followed a longitudinal pattern by regression analysis. Signatures assigned to *Ca.* Nitrosocosmicus (global mean = 0.17; global variance = 0.09; importance = 0.17) and other genera of the family *Nitrososphaeraceae* (global mean = 0.33; global variance = 0.15; importance = 0.12) achieved high importance in linear regression analysis over time supporting our assumption that AOA represent a coherent longitudinal trait on human skin (Supplementary Table S17; data type: Lqual2; Supplementary Fig. 2). Further information on bacterial ammonia oxidizers can be found in the Supplementary Information.

### AOA are an integral component of the healthy human skin microbiome

Network analyses (data types: Lqual1 and Lqual2; Supplementary Fig. 2) were performed to understand the integration of AOA into the skin microbiome. These showed a significant positive correlation of *Ca.* Nitrosocosmicus with integral bacterial members of the human skin microbiome (in particular *Lawsonella* and *Finegoldia* SCNIC sparCC p-adjust 0.26; see also initial *in silico* metabolic modeling in Supplementary Fig. 6). In contrast to other microbiome components, this interaction was relatively stable across all sampling time points (Fig. 5A, as visualized by the dark color of the node, see legend). A specific integration of *Ca.* Nitrosocosmicus into the skin microbiome network was underscored by the lack of positive correlations of other archaeal signatures that are not common on human skin (e.g., *Methanobrevibacter*).

**Fig. 5.**
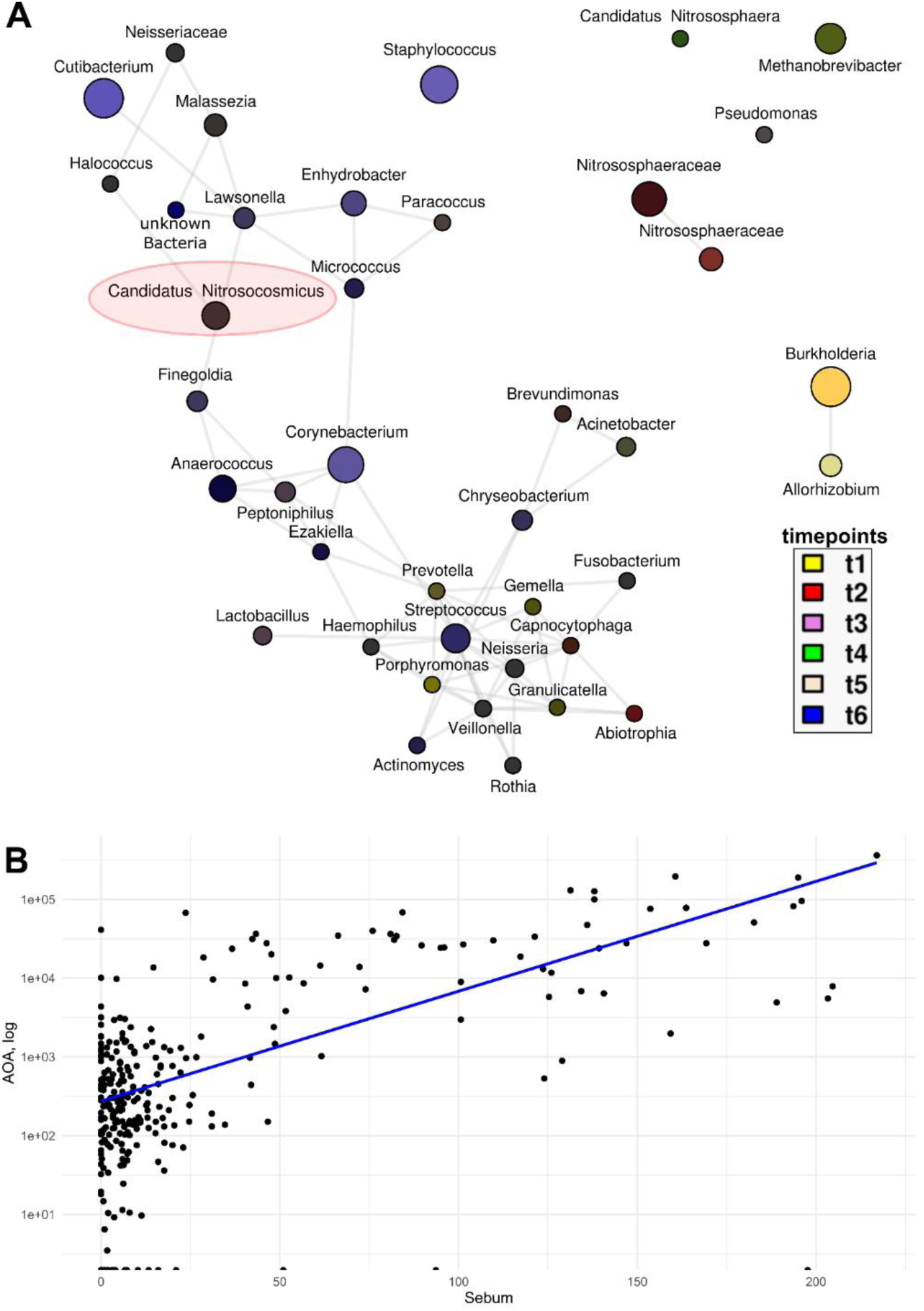
(A) Microbial co-occurrence network (n = 283; data types: Lqual1 and Lqual2; Supplementary Fig. 2) between the 40 most abundant archaeal and bacterial taxa (SRS normalized Cmin = 100, TSS and log transformed). Significant <u>positive</u> associations from Spearman correlations are highlighted by connecting edges. Taxa (nodes) were colored according to color mixtures of each sampling event (t1 yellow, t2 red, t3 pink, t4 green, t5 beige, t6 blue), resulting in dark gray colors for longitudinally consistent taxa in the dataset. The location of *Ca.* Nitrosocosmicus within the network is highlighted by a red ellipse. (B) Spearman correlation plot of AOA signatures and sebum content of the skin.

Ordinary least squares (OLS) regression models were used to find associations between archaeal taxa and acquired metadata (skin physiology, demographics of the cohort and their lifestyle as well as health parameters; data type: Cqual2; Supplementary Fig. 2). Considering the compositional nature of our amplicon datasets we used CLR (center log ratio) transformed values and regressed them with all numerical and categorical metadata categories while taking care of several potential confounders (which partly had significant impact on the microbial communities) like a subject’s sex, age, BMI, different sampled body sites and the overall microbial diversity (Shannon diversity). According to this analysis, signatures of Nitrososphaeraceae (n = 156) were significantly positively associated (q = 0.02) with qPCR-retrieved quantity for AOA (Supplementary Table S18). This finding supports the congruence of amplicon-based qualitative and qPCR-based quantitative molecular analyses.

The abundance of *amoA* genes as detected by qPCR (tp1, n=48) was not significantly influenced by sex (female vs. male) and age group (group A1 (20-30 years) and A2 (>60 years)). However, the abundance of *amoA* signatures negatively correlated with lower pH (rho= -0.081, p=0.174), and significantly negatively with increasing age (rho=-0.16, p=7.7*10^-3^), trans-epidermal water-loss (“Tewameter”, rho=0.22, p=4.8*10^-4^), and particularly sebum concentration (rho=0.47, p=1.0*10^-16^) according to Spearman rank correlations (data type: Cquant3; Supplementary Fig. 2).

To assess the possibility whether AOA are involved in psoriatic disease, as hypothesized earlier (Probst et al., 2013), 20 subjects were recruited and n=72 samples from lesioned and healthy skin samples were taken (Cohort C, data type: P; Supplementary Fig. 2). However, according to differential abundance analysis based on ANCOM-BC (p adjust = 0.07) and MaAsLin2 (q-value = 0.54) samples from psoriatic skin did not reveal significantly higher or lower AOA counts (data type: Pqual2; Supplementary Fig. 2).

## Discussion

In this study, we demonstrate the consistent presence of AOA of the genus *Ca. Nitrosocosmicus* on human skin, using two independent, but complementary, approaches: cultivation and molecular profiling. Collectively, our results suggest that *Ca.* Nitrosocosmicus representatives are an integral component of the skin microbiome. This conclusion is supported by:

i. the repeated enrichment of *Ca.* Nitrosocosmicus strains from independent human skin samples
ii. unique genomic features indicative of habitat-specific adaptation
iii. their high prevalence and distinct biogeographic patterns in molecular analyses, and
iv. specific co-occurrence with established skin-associated bacterial taxa

These findings not only highlight the ecological relevance of *Ca. Nitrosocosmicus* on human skin but also raise important questions regarding their functional role. Both highly enriched AOA cultures exhibited a stoichiometric conversion of ammonia to nitrite that is typical of other isolated AOA species, and their genomes encode urease, enabling urea utilization. Given that ammonia and urea are the primary nitrogenous species excreted on human skin through sweat (Chen et al., 2020), these substrates likely represent a stable energy source for these organisms.

Although the archaeal ammonia oxidation pathway has yet to be resolved, the intermediate nitric oxide (NO) has been observed to play a critical role for these organisms (Jessica A Kozlowski et al., 2016). Since NO is one of the most important signaling molecules in mammalian physiology, and is involved in a range of physiological as well as pathological processes on the human skin (reviewed in (Cals-Grierson and Ormerod, 2004; Suschek et al., 2010)), AOA could contribute to the maintenance of NOx homeostasis on the human skin and thereby play a role in skin health or physiology. It is also worth noting that nitrite, the final product of nitrification, also serves as a signaling molecule in a NO-independent manner (Bryan et al., 2005; Gladwin et al., 2006).

Both enriched strains were placed phylogenetically within the genus *Ca.* Nitrosocosmicus, a genus that was also consistently detected in cohorts A and B of an independent cross-sectional as well as longitudinal study. Although the relative abundance appears low in these human samples, it is not outside the range observed for AOA in other environments (Alves et al., 2019; Leininger et al., 2006).

To date, *Ca.* Nitrosocosmicus species have predominantly been isolated from soil environments, and are known for their aggregate structures, propensity to grow in biofilms, and superior adhesion capacities (Dreer et al., 2024; Jung et al., 2016; Liu et al., 2021). These traits are supported by the specific encoding or expansion of adhesion proteins, acetamido sugar biosynthesis and export proteins, as well as protein families whose products can directly interact with and modulate other organisms, such as peptidases, phospholipases and triacylglycerol esterases/lipases ((Jung et al., 2016; Liu et al., 2021) and this study), with known roles in tissue invasion and colonization, virulence and mediation of the host immune response (Garcia et al., 2018).

All of these features support the interaction of this genus with a variety of species in their respective environments and is further supported by recent evidence of *Ca.* Nitrosocosmicus associations and interactions with the plant rhizosphere (Lee et al., 2024) . The strains *Ca.* N. epidermidis and *Ca.* N. unguis share these genomics features while also displaying unique attributes suggestive of skin-specific functions.

Host interaction capabilities and attachment strategies are well established among skin-associated microorganisms (Brandwein et al., 2016), and our data indicate that *Ca. N. epidermidis* exhibits similar potential. Its genome harbors a genomic island that is enriched in proteins that facilitate surface adhesion and interactions with various tissue types and proteins that could putatively interact with eukaryotic proteins including an autotransporter (Mondal et al., 2022) and arrestin-like protein (Alvarez, 2008; Marullo and Coureuil, 2014). Similarly, *Ca.* N. unguis has an unusually high amount of cysteine-rich secretory protein (CAP) family proteins and beta propeller fold containing proteins that are both known for mediating interactions with the surrounding environment including signaling and regulation of host proteins (Chen et al., 2011; Gibbs et al., 2008; Yeats et al., 2003). The distinct repertoire of each strain may reflect micro-habitat preferences on the heterogeneous skin landscape.

Currently, it remains difficult to unequivocally identify proteins that are specific to the skin environment, but this is not a *Ca.* Nitrosocosmicus-specific difficulty; many skin-associated microorganisms display ecological flexibility occupying both, mammalian host-associated and environmental niches. Such dual lifestyles are well documented across microbial taxa.

This holds true for archaea in the case of the order Methanomassiliicoccales, which are able to thrive in both the gastrointestinal tract of mammals and diverse natural habitats (Cozannet et al., 2020; De La Cuesta-Zuluaga et al., 2021), as well as for many opportunistic pathogens of bacteria including *Vibrio sp.* (aquatic environments(Soto and Nishiguchi, 2021)), *Stenotrophomonas maltophilia* (soil, plant environments (Brooke, 2012)), or *Bukholderia pseudomallei* (Ghazali et al., 2023). The constant interaction between host and environment can further confound the identification of host-related proteins without further in-depth analysis of individual strains.

Regardless, the consistent presence of AOA on human skin within the longitudinal study supports the proposal of *Ca.* Nitrosocosmicus sp. being an emerging commensal skin microorganism. Their presence was found to be independent of age, sex, or time of year suggesting that they are a stable and integral component of the microbial epidermal skin microbiome. The detected presence additionally showed correlations with skin parameters, in particular a positive correlation with sebum content. Sebum is an oily substance that contains many lipids with hydrophobic properties and this correlation suggests that this AOA lineage can thrive in this type of environment. This fits not only with the several lipid interaction genes found within *Nitrososcosmicus,* but also with the placement of *Ca.* N. epidermidis, *Ca.* N. unguis, and several skin AOA ASVs next to the AOA species *Ca.* N. *oleophilus* which was isolated from coal-tar contaminated soil and exhibited a strong affinity for hydrophobic surfaces (Jung et al., 2016). The positive correlations with both the genera *Lawsonella* and *Finegoldia* are also revealing as both have been associated with high sebum areas such as the forehead ((Kim et al., 2021; Li et al., 2024).These co-occurrence patterns may reflect ecological interdependencies or shared niche preferences.

The metabolic activity of AOA, particularly their role in nitrogen cycling, may be crucial in shaping the skin microbiome. Given the scarcity of AOB on human skin, AOA likely fulfill this ecological function. Additionally, the capacity to produce crucial vitamins, such as B12 (Bayer et al., 2019), via an autotrophic metabolism may also play an important role in the community structure of the skin microbiome as it does in other environments.

Their interactions with integral bacterial members of the human microbiome may also have a positive impact on the host that is so far unknown, as no association with psoriasis was found. Nevertheless, many aspects of skin health remain unexplored in the context of archaeal involvement. Based on our findings, we can conclude, that *Ca.* Nitrosocosmicus is an integral and stable component of the microbial epidermal skin microbiome that is associated with relevant bacteria potentially via the exchange of ammonia and is characteristic of healthy skin with higher sebum content, less moisture and lower pH.

## Conclusions

In conclusion, this study marks the first AOA enrichments obtained from human skin and confirms the presence of the obtained genera, *Ca.* Nitrosocosmicus, on a wide range of individuals of varying sex and age. Initial genomic and physiological observations suggest that these AOA are adapted for growth on hydrophobic tissue surfaces as both enriched strains contain unique genes and protein families compared to other *Ca.* Nitrosocosmicus species involved in either adhesion or extracellular protein interaction. Further studies of skin archaea will be needed to reveal their evolutionary paths towards commensalism and to elucidate their roles in the skin microbiome and human health. The archaeal cultures obtained in this study are crucial prerequisites for advancing our knowledge in this area.

## Declarations

### Ethics approval and consent to participate

Subjects for microbiome analyses were recruited as part of the project “Detection of Archaea on Healthy and Diseased Skin” with the ethical vote EK: 27-289 ex 14/15 (Medical University of Graz) and all ethical requirements were followed and fulfilled including an informed consent to participate in the presented study. The following exclusion criteria were taken into account: Antibiotics therapy within the last three months, pregnancy, ongoing breast-feeding and non-compliance. The participants were asked to not shower or apply skin cream the day before sampling.

### Consent for publication

All authors approved the manuscript and give their consent for publication.

### Availability of data and material

All raw sequencing data were submitted to NCBI’s Sequence Read Archive (SRA) https://www.ncbi.nlm.nih.gov/sra and are accessible under the following projects: PRJNA1105887 (archaea specific 16S rRNA gene amplicons of cohort A; data type: Cqual2), PRJNA1107153 (universal 16S rRNA gene amplicons of cohort A; data type: Cqual1), PRJNA1107462 (AOA specific *amoA* gene amplicons of cohort A; data type: Cqual4), PRJNA1107502 (archaea specific 16S rRNA gene amplicons of cohort B; data type: Lqual2), PRJNA1107512 (universal 16S rRNA gene amplicons of cohort B; data type: Lqual1), PRJNA1105887 (archaea specific 16S rRNA gene amplicons of cohort C; data type: Pqual2), PRJNA1108073 (universal 16S rRNA gene amplicons of cohort C; data type: Pqual1), PRJNA1108908 (pooled shotgun metagenomics data of cohort A), PRJNA1108960 (Illumina sequencing data of enrichments), PRJNA1109334 (Nanopore sequencing data of enrichments). Assemblies are publicly available under NCBI BioProjects PRJNA1103140 (*Ca.* Nitrosocosmicus epidermidis T1S) and PRJNA1103141 (*Ca.* Nitrosocosmicus unguis). Annotated assemblies are also available at the MicroScope Microbial Genome Annotation and Analysis Platform, developed by the Laboratory of Bioinformatics Analyses for Genomics and Metabolism, Genoscope, Evry, France.

### Code availability

The present study did not generate new code, and mentioned tools used for the data analysis were applied with default parameters unless specified otherwise in Methods and our Github repo. https://github.com/CME-lab-research/SkinArchaeome

## Authors’ contributions

AM supervised and performed microbiome sampling and analysis. MD supervised and performed cultivation and microscopy. MM and LH supervised and supported cultivation work. UP performed cultivation. SD, AL, TG, DB performed sampling, sequencing, and data analysis of qPCR and amplicon data. RP performed phylogenetic analyses and genome assembly. TG and PW recruited patients and performed sampling. MK performed genome annotations and contributed to study concept. CME and CS developed the study concept, wrote initial proposal, and wrote, together with MK and AM the manuscript. All authors contributed to the finalization of the manuscript.

## Supporting information

Supplementary Data Annotation

Supplementary Data Orthologs

## Acknowledgements

We acknowledge the technical support by Ignaz Herzeg, Lisa Wink, Polona Mertelj, Sarah Harrer and all collaborators at the Department of Dermatology and Venereology, Medical University of Graz (Andrea Alejandra Catalan Griffiths, Patra Vijaykumar, Isabella Perchthaler, Alexandra Gruber-Wackernagel, Wolfgang Weger, and univ. Prof. Dr. Wolfgang Salmhofer). We thank Daniela Gruber from the Core Facility Cell Imaging & Ultrastructure Research (LeWi).

We greatly value the computational resources of the MedBioNode provided by the Medical University of Graz and the Life Science Compute Cluster (LiSC) operated by the Computational Systems Biology group at the University of Vienna as well as the Medical University of Graz ZMF Galaxy Team: Core Facility Computational Bioanalytics, Medical University of Graz, funded by the Austrian Federal Ministry of Education, Science and Research, Hochschulraum-Strukturmittel 2016 grant as part of BioTechMed Graz. SD was supported by the local PhD program MolMed.

## List of abbreviations

(AOA): Ammonia-oxidizing archaea
(amoA): ammonia monooxygenase subunit A gene
(NO): nitric oxide
(DNA): Deoxyribonucleic Acid
(RNA): Ribonucleic Acid
(PCR): Polymerase chain reaction
(qPCR): Quantitative polymerase chain reaction
(FWM): freshwater medium
(MOPS): 3-(N-morpholino) propanesulfonic acid
(HCl): Hydrogen chloride
(PBS): Phosphate buffered saline
(HMW): High molecular weight
(MCE): mixed cellulose ester
(ONT): Oxford Nanopore Technologies
(LiSC): Life Science Compute Cluster
(arCOG): archaeal Clusters of Orthologous Genes
(CAZymes): Carbohydrate-active enzymes assignments
(QIIME): Quantitative Insights Into Microbial Ecology
(DADA): Divisive Amplicon Denoising Algorithm
(MAFFT): Multiple Alignment using Fast Fourier Transform
(ANCOM-BC): Analysis of Compositions of Microbiomes with Bias Correction
(MaAsLin): Microbiome Multivariable Associations with Linear Models
(Aldex): ANOVA-Like Differential Expression tool
(DEICODE): Differential Eigenvalue Interpretation for Compositional Omics Data Exploration
(RESCRIPt): Reference Sequence Curation Pipeline Tool
(OLS): ordinary least squares
(CLR): center log ratio
(FDR): false discovery rate
(MAG): metagenome assembled genome
(BMI): Body Mass Index
(PCoA): Principal Coordinates Analysis
(PCA): Principal Component Analysis
(SCNIC): Sparse Cooccurrence Network Investigation for Compositional data
(ML): maximum likelihood
(SINA): SILVA Incremental Aligner
(SSU): Small Subunit
(BLASTN): Basic Local Alignment Search Tool for Nucleotides
(ANI): Average Nucleotide Identity
(AAI): Amino Acid Identity
(NS): Nitrososphaerales
(ASV): Amplicon Sequence Variant

## References

1. Abby, S.S., Melcher, M., Kerou, M., Krupovic, M., Stieglmeier, M., Rossel, C., Pfeifer, K., Schleper, C., 2018. Candidatus Nitrosocaldus cavascurensis, an ammonia oxidizing, extremely thermophilic archaeon with a highly mobile genome. Front Microbiol 9, 346006. 10.3389/FMICB.2018.00028/BIBTEX

2. Alvarez, C.E., 2008. On the origins of arrestin and rhodopsin. BMC Evol Biol 8, 222. 10.1186/1471-2148-8-222

3. Alves, R.J.E., Kerou, M., Zappe, A., Bittner, R., Abby, S.S., Schmidt, H.A., Pfeifer, K., Schleper, C., 2019. Ammonia oxidation by the arctic terrestrial thaumarchaeote candidatus nitrosocosmicus arcticus is stimulated by increasing temperatures. Front Microbiol 10, 1571. 10.3389/FMICB.2019.01571/FULL

4. Alves, R.J.E., Minh, B.Q., Urich, T., Haeseler, A., Schleper, C., 2018. Unifying the global phylogeny and environmental distribution of ammonia-oxidising archaea based on amoA genes. Nat Commun 9, 1517.

5. Arce, M.I., von Schiller, D., Bengtsson, M.M., Hinze, C., Jung, H., Alves, R.J.E., Urich, T., Singer, G., 2018. Drying and Rainfall Shape the Structure and Functioning of Nitrifying Microbial Communities in Riverbed Sediments. Front Microbiol 9. 10.3389/fmicb.2018.02794

6. Bang, C., Schmitz, R.A., 2015. Archaea associated with human surfaces: not to be underestimated. FEMS Microbiol Rev 39, 631–648. 10.1093/femsre/fuv010

7. Bayer, B., Hansman, R.L., Bittner, M.J., Noriega-Ortega, B.E., Niggemann, J., Dittmar, T., Herndl, G.J., 2019. Ammonia-oxidizing archaea release a suite of organic compounds potentially fueling prokaryotic heterotrophy in the ocean. Environ Microbiol 21, 4062– 4075. 10.1111/1462-2920.14755

8. Bengtsson-Palme, J., Ryberg, M., Hartmann, M., Branco, S., Wang, Z., Godhe, A., 2013. Improved software detection and extraction of ITS1 and ITS2 from ribosomal ITS sequences of fungi and other eukaryotes for analysis of environmental sequencing data. Meth Ecol Evol 4.

9. Berg, G., Rybakova, D., Fischer, D., Cernava, T., Vergès, M.-C.C., Charles, T., Chen, X., Cocolin, L., Eversole, K., Corral, G.H., 2020. Microbiome definition re-visited: old concepts and new challenges. Microbiome 8, 1–22.

10. Bokulich, N., Dillon, M., Bolyen, E., Kaehler, B., Huttley, G., Caporaso, J., 2018. q2-sample-classifier: machine-learning tools for microbiome classification and regression. J Open Source Softw 3, 934. 10.21105/joss.00934

11. Bolger, A.M., Lohse, M., Usadel, B., 2014. Trimmomatic: a flexible trimmer for Illumina sequence data. Bioinformatics 30, 2114–2120. 10.1093/bioinformatics/btu170

12. Bolyen, E., Rideout, J.R., Dillon, M.R., Bokulich, N.A., Abnet, C., Ghalith, G.A. Al, Alexander, H., Alm, E.J., Arumugam, M., Bai, Y., Bisanz, J.E., Bittinger, K., Brejnrod, A., Colin, J., Brown, C.T., Callahan, B.J., Mauricio, A., Rodríguez, C., Chase, J., Cope, E., Silva, R. Da, Dorrestein, P.C., Douglas, G.M., Duvallet, C., Edwardson, C.F., Ernst, M., Fouquier, J., Gauglitz, J.M., Gibson, D.L., Gonzalez, A., Huttley, G.A., Janssen, S., Jarmusch, A.K., Kaehler, B.D., Kang, K. Bin, Keefe, C.R., Keim, P., Kelley, S.T., Ley, R., Loftfield, E., Marotz, C., Martin, B., Mcdonald, D., Mciver, L.J., Alexey, V., Metcalf, J.L., Morgan, S.C., Morton, J.T., Naimey, A.T., 2018. QIIME 2 : Reproducible , interactive , scalable , and extensible microbiome data science. 10.7287/peerj.preprints.27295

13. Borrel, G., Brugère, J.F., Gribaldo, S., Schmitz, R.A., Moissl-Eichinger, C., 2020. The host-associated archaeome. Nat Rev Microbiol. 10.1038/s41579-020-0407-y

14. Brandwein, M., Steinberg, D., Meshner, S., 2016. Microbial biofilms and the human skin microbiome. npj Biofilms and Microbiomes 2016 2:1 2, 1–6. 10.1038/s41522-016-0004-z

15. Brooke, J.S., 2012. Stenotrophomonas maltophilia: an Emerging Global Opportunistic Pathogen. Clin Microbiol Rev 25, 2–41. 10.1128/CMR.00019-11

16. Bryan, N.S., Fernandez, B.O., Bauer, S.M., Garcia-Saura, M.F., Milsom, A.B., Rassaf, T., Maloney, R.E., Bharti, A., Rodriguez, J., Feelisch, M., 2005. Nitrite is a signaling molecule and regulator of gene expression in mammalian tissues. Nat Chem Biol 1, 290–297. 10.1038/nchembio734

17. Buchfink, B., Xie, C., Huson, D.H., 2015. Fast and sensitive protein alignment using DIAMOND. Nat Methods 12, 59. 10.1038/nmeth.3176

18. Byrd, A.L., Belkaid, Y., Segre, J.A., 2018. The human skin microbiome. Nat Rev Microbiol 16, 143–155. 10.1038/nrmicro.2017.157

19. Callahan, B.J., McMurdie, P.J., Rosen, M.J., Han, A.W., Johnson, A.J.A., Holmes, S.P., 2016. DADA2: High-resolution sample inference from Illumina amplicon data. Nat Methods 13, 581–583. doi:10.1038/nmeth.3869

20. Cals-Grierson, M.-M., Ormerod, A.D., 2004. Nitric oxide function in the skin. Nitric Oxide 10, 179–193. 10.1016/j.niox.2004.04.005

21. Cantalapiedra, C.P., Hernández-Plaza, A., Letunic, I., Bork, P., Huerta-Cepas, J., 2021. eggNOG-mapper v2: Functional Annotation, Orthology Assignments, and Domain Prediction at the Metagenomic Scale. Mol Biol Evol 38, 5825–5829. 10.1093/molbev/msab293

22. Chan, J.M., Gori, A., Nobbs, A.H., Heyderman, R.S., 2020. Streptococcal Serine-Rich Repeat Proteins in Colonization and Disease. Front Microbiol 11. 10.3389/fmicb.2020.593356

23. Chaumeil, P.A., Mussig, A.J., Hugenholtz, P., Parks, D.H., 2020. GTDB-Tk: A toolkit to classify genomes with the genome taxonomy database. Bioinformatics 36, 1925–1927. 10.1093/bioinformatics/btz848

24. Chen, C.K.-M., Chan, N.-L., Wang, A.H.-J., 2011. The many blades of the β-propeller proteins: conserved but versatile. Trends Biochem Sci 36, 553–561. 10.1016/j.tibs.2011.07.004

25. Chen, Y.-L., Kuan, W.-H., Liu, C.-L., 2020. Comparative Study of the Composition of Sweat from Eccrine and Apocrine Sweat Glands during Exercise and in Heat. Int J Environ Res Public Health 17, 3377. 10.3390/ijerph17103377

26. Chibani, C.M., Mahnert, A., Borrel, G., Almeida, A., Werner, A., Brugère, J.-F., Gribaldo, S., Finn, R.D., Schmitz, R.A., Moissl-Eichinger, C., 2022. A catalogue of 1,167 genomes from the human gut archaeome. Nat Microbiol 7, 48–61. 10.1038/s41564-021-01020-9

27. Chklovski, A., Parks, D.H., Woodcroft, B.J., Tyson, G.W., 2022. CheckM2: a rapid, scalable and accurate tool for assessing microbial genome quality using machine learning. bioRxiv 2022.07.11.499243. 10.1101/2022.07.11.499243

28. Cozannet, M., Borrel, G., Roussel, E., Moalic, Y., Allioux, M., Sanvoisin, A., Toffin, L., Alain, K., 2020. New Insights into the Ecology and Physiology of Methanomassiliicoccales from Terrestrial and Aquatic Environments. Microorganisms 9, 30. 10.3390/microorganisms9010030

29. Criscuolo, A., Gribaldo, S., 2010. BMGE (Block Mapping and Gathering with Entropy): a new software for selection of phylogenetic informative regions from multiple sequence alignments. BMC Evol Biol 10, 210. 10.1186/1471-2148-10-210

30. Davis, N.M., Proctor, D.M., Holmes, S.P., Relman, D.A. B.J. C., 2018. Simple statistical identification and removal of contaminant sequences in marker-gene and metagenomics data. Microbiome 6, 226. 10.1186/s40168-018-0605-2

31. De Coster, W., D’Hert, S., Schultz, D.T., Cruts, M., Van Broeckhoven, C., 2018. NanoPack: visualizing and processing long-read sequencing data. Bioinformatics 34, 2666–2669. 10.1093/bioinformatics/bty149

32. De La Cuesta-Zuluaga, J., Spector, T.D., Youngblut, N.D., Ley, R.E., 2021. Genomic Insights into Adaptations of Trimethylamine-Utilizing Methanogens to Diverse Habitats, Including the Human Gut. mSystems 6, e00939–20.

33. De La Torre, J.R., Walker, C.B., Ingalls, A.E., Könneke, M., Stahl, D.A., 2008. Cultivation of a thermophilic ammonia oxidizing archaeon synthesizing crenarchaeol. Environ Microbiol 10, 810–818. 10.1111/j.1462-2920.2007.01506.x

34. Dombrowski, N., Williams, T.A., Sun, J., Woodcroft, B.J., Lee, J.-H., Minh, B.Q., Rinke, C., Spang, A., 2020. Undinarchaeota illuminate DPANN phylogeny and the impact of gene transfer on archaeal evolution. Nat Commun 11, 3939. 10.1038/s41467-020-17408-w

35. Dreer, M., Pribasnig, T., Hodgskiss, L.H., Luo, Z.-H., Pozaric, F., Schleper, C., 2024. Biofilm lifestyle as a common trait of ammonia-oxidizing archaea. Biorxiv. 10.1101/2024.11.18.624116

36. Eddy, S.R., 2011. Accelerated Profile HMM Searches. PLoS Comput Biol 7, e1002195. 10.1371/journal.pcbi.1002195

37. Emms, D.M., Kelly, S., 2019. OrthoFinder: phylogenetic orthology inference for comparative genomics. Genome Biol 20, 238. 10.1186/s13059-019-1832-y

38. Fan, Y., Pedersen, O., 2020. Gut microbiota in human metabolic health and disease. Nature Reviews Microbiology 2020 19:1 19, 55–71. 10.1038/s41579-020-0433-9

39. Fan, Y., Wu, L., Zhai, B., 2023. The mycobiome: interactions with host and implications in diseases. Curr Opin Microbiol 75, 102361. 10.1016/J.MIB.2023.102361

40. Fernandes, A.D., Macklaim, J.M., Linn, T.G., Reid, G., Gloor, G.B., 2013. ANOVA-Like Differential Expression (ALDEx) Analysis for Mixed Population RNA-Seq. PLoS One 8, e67019. 10.1371/journal.pone.0067019

41. Flowers, L., Grice, E.A., 2020. The Skin Microbiota: Balancing Risk and Reward. 10.1016/j.chom.2020.06.017

42. Gaci, N., Borrel, G., Tottey, W., O’Toole, P.W., Brugère, J.-F., 2014. Archaea and the human gut: New beginning of an old story. World J Gastroenterol 20, 16062. 10.3748/wjg.v20.i43.16062

43. Garcia, C.J., Pericleous, A., Elsayed, M., Tran, M., Gupta, S., Callaghan, J.D., Stella, N.A., Franks, J.M., Thibodeau, P.H., Shanks, R.M.Q., Kadouri, D.E., 2018. Serralysin family metalloproteases protects Serratia marcescens from predation by the predatory bacteria Micavibrio aeruginosavorus. Sci Rep 8, 14025. 10.1038/s41598-018-32330-4

44. Ghazali, A.-K., Firdaus-Raih, M., Uthaya Kumar, A., Lee, W.-K., Hoh, C.-C., Nathan, S., 2023. Transitioning from Soil to Host: Comparative Transcriptome Analysis Reveals the Burkholderia pseudomallei Response to Different Niches. Microbiol Spectr 11. 10.1128/spectrum.03835-22

45. Gibbs, G.M., Roelants, K., O’Bryan, M.K., 2008. The CAP Superfamily: Cysteine-Rich Secretory Proteins, Antigen 5, and Pathogenesis-Related 1 Proteins—Roles in Reproduction, Cancer, and Immune Defense. Endocr Rev 29, 865–897. 10.1210/er.2008-0032

46. Gladwin, M.T., Raat, N.J.H., Shiva, S., Dezfulian, C., Hogg, N., Kim-Shapiro, D.B., Patel, R.P., 2006. Nitrite as a vascular endocrine nitric oxide reservoir that contributes to hypoxic signaling, cytoprotection, and vasodilation. American Journal of Physiology-Heart and Circulatory Physiology 291, H2026–H2035. 10.1152/ajpheart.00407.2006

47. Gomaa, E.Z., 2020. Human gut microbiota/microbiome in health and diseases: a review. Antonie Van Leeuwenhoek 113, 2019–2040. 10.1007/s10482-020-01474-7

48. Graham, E.D., Heidelberg, J.F., Tully, B.J., 2018. Potential for primary productivity in a globally-distributed bacterial phototroph. ISME J 12, 1861–1866. 10.1038/s41396-018-0091-3

49. Gremme, G., Steinbiss, S., Kurtz, S., 2013. GenomeTools: A Comprehensive Software Library for Efficient Processing of Structured Genome Annotations. IEEE/ACM Trans Comput Biol Bioinform 10, 645–656. 10.1109/TCBB.2013.68

50. Grice, E.A., Segre, J.A., 2011. The skin microbiome. Nat Rev Microbiol 9, 244–253.

51. Großkopf, R., Janssen, P.H., Liesack, W., 1998. Diversity and structure of the methanogenic community in anoxic rice paddy soil microcosms as examined by cultivation and direct 16S rRNA gene sequence retrieval. Appl Environ Microbiol 64, 960–969.

52. Guindon, S., Dufayard, J.-F., Lefort, V., Anisimova, M., Hordijk, W., Gascuel, O., 2010. New Algorithms and Methods to Estimate Maximum-Likelihood Phylogenies: Assessing the Performance of PhyML 3.0. Syst Biol 59, 307–321. 10.1093/sysbio/syq010

53. Heidrich, V., Karlovsky, P., Beule, L., 2021. ‘SRS’ R Package and ‘q2-srs’ QIIME 2 Plugin: Normalization of Microbiome Data Using Scaling with Ranked Subsampling (SRS). Applied Sciences 11, 11473. 10.3390/app112311473

54. Huerta-Cepas, J., Szklarczyk, D., Heller, D., Hernández-Plaza, A., Forslund, S.K., Cook, H., Mende, D.R., Letunic, I., Rattei, T., Jensen, L.J., 2019. eggNOG 5.0: a hierarchical, functionally and phylogenetically annotated orthology resource based on 5090 organisms and 2502 viruses. Nucleic Acids Res 47, D309–D314.

55. Hyatt, D., Chen, G.-L., LoCascio, P.F., Land, M.L., Larimer, F.W., Hauser, L.J., 2010. Prodigal: prokaryotic gene recognition and translation initiation site identification. BMC Bioinformatics 11, 119. 10.1186/1471-2105-11-119

56. Jung, M.Y., Kim, J.G., Sinninghe Damsté, J.S., Rijpstra, W.I.C., Madsen, E.L., Kim, S.J., Hong, H., Si, O.J., Kerou, M., Schleper, C., Rhee, S.K., 2016. A hydrophobic ammonia-oxidizing archaeon of the Nitrosocosmicus clade isolated from coal tar-contaminated sediment. Environ Microbiol Rep 8, 983–992. 10.1111/1758-2229.12477

57. Kalyaanamoorthy, S., Minh, B.Q., Wong, T.K.F., von Haeseler, A., Jermiin, L.S., 2017. ModelFinder: fast model selection for accurate phylogenetic estimates. Nat Methods 14, 587–589. 10.1038/nmeth.4285

58. Katoh, K., Standley, D.M., 2013. MAFFT Multiple Sequence Alignment Software Version 7: Improvements in Performance and Usability. Mol Biol Evol 30, 772–780. 10.1093/molbev/mst010

59. Kim, J.-H., Son, S.-M., Park, H., Kim, B.K., Choi, I.S., Kim, H., Huh, C.S., 2021. Taxonomic profiling of skin microbiome and correlation with clinical skin parameters in healthy Koreans. Sci Rep 11, 16269. 10.1038/s41598-021-95734-9

60. Klein, T., Poghosyan, L., Barclay, J.E., Murrell, J.C., Hutchings, M.I., Lehtovirta-Morley, L.E., 2022. Cultivation of ammonia-oxidising archaea on solid medium. FEMS Microbiol Lett 369. 10.1093/femsle/fnac029

61. Klindworth, A., Pruesse, E., Schweer, T., Peplies, J., Quast, C., Horn, M., Glockner, F.O., 2013. Evaluation of general 16S ribosomal RNA gene PCR primers for classical and next-generation sequencing-based diversity studies. Nucleic Acids Res 41, e1–e1. 10.1093/nar/gks808

62. Kobe, B., 2001. The leucine-rich repeat as a protein recognition motif. Curr Opin Struct Biol 11, 725–732. 10.1016/S0959-440X(01)00266-4

63. Kolmogorov, M., Bickhart, D.M., Behsaz, B., Gurevich, A., Rayko, M., Shin, S.B., Kuhn, K., Yuan, J., Polevikov, E., Smith, T.P.L., Pevzner, P.A., 2020. metaFlye: scalable long-read metagenome assembly using repeat graphs. Nat Methods 17, 1103–1110. 10.1038/s41592-020-00971-x

64. Kong, H.H., 2011. Skin microbiome: genomics-based insights into the diversity and role of skin microbes. Trends Mol Med 17, 320. 10.1016/J.MOLMED.2011.01.013

65. Konstantinidis, K.T., Rosselló-Móra, R., Amann, R., 2017. Uncultivated microbes in need of their own taxonomy. ISME J 11, 2399–2406. 10.1038/ismej.2017.113

66. Kozlowski, Jessica A., Kits, K.D., Stein, L.Y., 2016. Comparison of Nitrogen Oxide Metabolism among Diverse Ammonia-Oxidizing Bacteria. Front Microbiol 7. 10.3389/fmicb.2016.01090

67. Kozlowski, Jessica A, Stieglmeier, M., Schleper, C., Klotz, M.G., Stein, L.Y., 2016. Pathways and key intermediates required for obligate aerobic ammonia-dependent chemolithotrophy in bacteria and Thaumarchaeota. ISME J 10, 1836–1845. 10.1038/ismej.2016.2

68. Kumpitsch, C., Fischmeister, F.P.S., Mahnert, A., Lackner, S., Wilding, M., Sturm, C., Springer, A., Madl, T., Holasek, S., Högenauer, C., 2021. Reduced B12 uptake and increased gastrointestinal formate are associated with archaeome-mediated breath methane emission in humans. Microbiome 9, 1–18. 10.1186/s40168-021-01130-w

69. Lane, D., 1991. 16S/23S rRNA sequencing., in: Stackebrandt, E., Goodfellow, M. (Eds.), Nucleic Acid Techniques in Bacterial Systematics. John Wiley & Sons, Chichester, pp. 115–175.

70. Larsson, A., 2014. AliView: a fast and lightweight alignment viewer and editor for large datasets. Bioinformatics 30, 3276–3278. 10.1093/bioinformatics/btu531

71. Lee, U.-J., Gwak, J.-H., Choi, S., Jung, M.-Y., Lee, T.K., Ryu, H., Imisi Awala, S., Wanek, W., Wagner, M., Quan, Z.-X., Rhee, S.-K., 2024. “ Ca. Nitrosocosmicus” members are the dominant archaea associated with plant rhizospheres. mSphere 9. 10.1128/msphere.00821-24

72. Lehtovirta-Morley, L.E., Ross, J., Hink, L., Weber, E.B., Gubry-Rangin, C., Thion, C., Prosser, J.I., Nicol, G.W., 2016. Isolation of ‘Candidatus Nitrosocosmicus franklandus’, a novel ureolytic soil archaeal ammonia oxidiser with tolerance to high ammonia concentration. FEMS Microbiol Ecol 92, 57. 10.1093/FEMSEC/FIW057

73. Leininger, S., Urich, T., Schloter, M., Schwark, L., Qi, J., Nicol, G.W., Prosser, J.I., Schuster, S.C., Schleper, C., 2006. Archaea predominate among ammonia-oxidizing prokaryotes in soils. Nature 442, 806–809.

74. Li, H., 2021. New strategies to improve minimap2 alignment accuracy. Bioinformatics 37, 4572–4574. 10.1093/bioinformatics/btab705

75. Li, M., Kopylova, E., Mao, J., Namkoong, J., Sanders, J., Wu, J., 2024. Microbiome and lipidomic analysis reveal the interplay between skin bacteria and lipids in a cohort study. Front Microbiol 15. 10.3389/fmicb.2024.1383656

76. Lin, H., Peddada, S. Das, 2020. Analysis of compositions of microbiomes with bias correction. Nat Commun 11, 3514. 10.1038/s41467-020-17041-7

77. Liu, L., Liu, M., Jiang, Y., Lin, W., Luo, J., 2021. Production and Excretion of Polyamines To Tolerate High Ammonia, a Case Study on Soil Ammonia-Oxidizing Archaeon “ Candidatus Nitrosocosmicus agrestis.” mSystems 6. 10.1128/MSYSTEMS.01003-20

78. Lozupone, C., Knight, R., 2005. UniFrac: a New Phylogenetic Method for Comparing Microbial Communities. Appl Environ Microbiol 71, 8228–8235. 10.1128/AEM.71.12.8228-8235.2005

79. Lu, J., Salzberg, S.L., 2020. Ultrafast and accurate 16S rRNA microbial community analysis using Kraken 2. Microbiome 8, 124. 10.1186/s40168-020-00900-2

80. Mahnert, A., Blohs, M., Pausan, M.R., Moissl-Eichinger, C., 2018. The human archaeome: methodological pitfalls and knowledge gaps. Emerg Top Life Sci 2.4, 469–482. 10.1042/ETLS20180037

81. Makarova, K.S., Galperin, M.Y., Koonin, E. V., 2015. Comparative genomic analysis of evolutionarily conserved but functionally uncharacterized membrane proteins in archaea: Prediction of novel components of secretion, membrane remodeling and glycosylation systems. Biochimie 118, 302–312. 10.1016/j.biochi.2015.01.004

82. Mallick, H., Rahnavard, A., McIver, L.J., Ma, S., Zhang, Y., Nguyen, L.H., Tickle, T.L., Weingart, G., Ren, B., Schwager, E.H., Chatterjee, S., Thompson, K.N., Wilkinson, J.E., Subramanian, A., Lu, Y., Waldron, L., Paulson, J.N., Franzosa, E.A., Bravo, H.C., Huttenhower, C., 2021. Multivariable association discovery in population-scale meta-omics studies. PLoS Comput Biol 17, e1009442. 10.1371/journal.pcbi.1009442

83. Martino, C., Morton, J.T., Marotz, C.A., Thompson, L.R., Tripathi, A., Knight, R., Zengler, K., 2019. A Novel Sparse Compositional Technique Reveals Microbial Perturbations. mSystems 4. 10.1128/mSystems.00016-19

84. Marullo, S., Coureuil, M., 2014. Arrestins in host-pathogen interactions. Handb Exp Pharmacol 219, 361–74. 10.1007/978-3-642-41199-1_18

85. Meulenbroek, E.M., Peron Cane, C., Jala, I., Iwai, S., Moolenaar, G.F., Goosen, N., Pannu, N.S., 2013. UV damage endonuclease employs a novel dual-dinucleotide flipping mechanism to recognize different DNA lesions. Nucleic Acids Res 41, 1363–1371. 10.1093/nar/gks1127

86. Minh, B.Q., Schmidt, H.A., Chernomor, O., Schrempf, D., Woodhams, M.D., von Haeseler, A., Lanfear, R., 2020. IQ-TREE 2: New Models and Efficient Methods for Phylogenetic Inference in the Genomic Era. Mol Biol Evol 37, 1530–1534. 10.1093/molbev/msaa015

87. Moissl-Eichinger, C., 2011. Archaea in artificial environments: their presence in global spacecraft clean rooms and impact on planetary protection. ISME J 5, 209–19. 10.1038/ismej.2010.124

88. Moissl-Eichinger, C., Probst, A.J., Birarda, G., Auerbach, A., Koskinen, K., Wolf, P., Holman, H.-Y.N., 2017. Human age and skin physiology shape diversity and abundance of Archaea on skin. Sci Rep 7. 10.1038/s41598-017-04197-4

89. Mondal, A.K., Lata, K., Singh, M., Chatterjee, S., Chauhan, A., Puravankara, S., Chattopadhyay, K., 2022. Cryo-EM elucidates mechanism of action of bacterial pore-forming toxins. Biochimica et Biophysica Acta (BBA) - Biomembranes 1864, 184013. 10.1016/j.bbamem.2022.184013

90. Nayfach, S., Páez-Espino, D., Call, L., Low, S.J., Sberro, H., Ivanova, N.N., Proal, A.D., Fischbach, M.A., Bhatt, A.S., Hugenholtz, P., Kyrpides, N.C., 2021. Metagenomic compendium of 189,680 DNA viruses from the human gut microbiome. Nature Microbiology 2021 6:7 6, 960–970. 10.1038/s41564-021-00928-6

91. Nicol, G.W., Leininger, S., Schleper, C., Prosser, J.I., 2008. The influence of soil pH on the diversity, abundance and transcriptional activity of ammonia oxidizing archaea and bacteria. Environ Microbiol 10, 2966–2978. 10.1111/j.1462-2920.2008.01701.x

92. Ochsenreiter, T., Selezi, D., Quaiser, A., Bonch-Osmolovskaya, L., Schleper, C., 2003. Diversity and abundance of Crenarchaeota in terrestrial habitats studied by 16S RNA surveys and real time PCR. Environ Microbiol 5, 787–797. 10.1046/j.1462-2920.2003.00476.x

93. Oh, J., Byrd, A.L., Park, M., Kong, H.H., Segre, J.A., 2016. Temporal Stability of the Human Skin Microbiome. Cell 165, 854–866. 10.1016/j.cell.2016.04.008

94. Olm, M.R., Brown, C.T., Brooks, B., Banfield, J.F., 2017. DRep: A tool for fast and accurate genomic comparisons that enables improved genome recovery from metagenomes through de-replication. ISME Journal 11, 2864–2868. 10.1038/ismej.2017.126

95. Parks, D.H., Chuvochina, M., Chaumeil, P.-A., Rinke, C., Mussig, A.J., Hugenholtz, P., 2020. A complete domain-to-species taxonomy for Bacteria and Archaea. Nat Biotechnol 38, 1079–1086. 10.1038/s41587-020-0501-8

96. Parks, D.H., Chuvochina, M., Rinke, C., Mussig, A.J., Chaumeil, P.A., Hugenholtz, P., 2022. GTDB: an ongoing census of bacterial and archaeal diversity through a phylogenetically consistent, rank normalized and complete genome-based taxonomy. Nucleic Acids Res 50, D785–D794. 10.1093/NAR/GKAB776

97. Pausan, M.R., Csorba, C., Singer, G., Till, H., Schöpf, V., Santigli, E., Klug, B., Högenauer, C., Blohs, M., Moissl-Eichinger, C., 2019. Exploring the Archaeome: Detection of Archaeal Signatures in the Human Body. Front Microbiol 10. 10.3389/fmicb.2019.02796

98. Pester, M., Rattei, T., Flechl, S., Gröngröft, A., Richter, A., Overmann, J., Reinhold-Hurek, B., Loy, A., Wagner, M., 2012. amoA-based consensus phylogeny of ammonia-oxidizing archaea and deep sequencing of amoA genes from soils of four different geographic regions. Environ Microbiol 14, 525–539. 10.1111/j.1462-2920.2011.02666.x

99. Price, M.N., Dehal, P.S., Arkin, A.P., 2010. FastTree 2 - Approximately maximum-likelihood trees for large alignments. PLoS One 5. 10.1371/journal.pone.0009490

100. Probst, A.J., Auerbach, A.K., Moissl-Eichinger, C., 2013. Archaea on human skin. PLoS One 8, e65388. 10.1371/journal.pone.0065388

101. Pruesse, E., Peplies, J., Glöckner, F.O., 2012. SINA: accurate high-throughput multiple sequence alignment of ribosomal RNA genes. Bioinformatics 28, 1823–1829.

102. Rayan, G.M., Flournoy, D.J., 1987. Microbiologic flora of human fingernails. J Hand Surg Am 12, 605–607. 10.1016/S0363-5023(87)80217-4

103. Reyes, C., Hodgskiss, L.H., Baars, O., Kerou, M., Bayer, B., Schleper, C., Kraemer, S.M., 2020. Copper limiting threshold in the terrestrial ammonia oxidizing archaeon Nitrososphaera viennensis. Res Microbiol 171, 134–142. 10.1016/J.RESMIC.2020.01.003

104. Rinke, C., Chuvochina, M., Mussig, A.J., Chaumeil, P.-A., Davín, A.A., Waite, D.W., Whitman, W.B., Parks, D.H., Hugenholtz, P., 2021. A standardized archaeal taxonomy for the Genome Taxonomy Database. Nat Microbiol 1–14.

105. Rinke, C., Schwientek, P., Sczyrba, A., Ivanova, N.N., Anderson, I.J., Cheng, J.-F., Darling, A., Malfatti, S., Swan, B.K., Gies, E.A., Dodsworth, J.A., Hedlund, B.P., Tsiamis, G., Sievert, S.M., Liu, W.-T., Eisen, J.A., Hallam, S.J., Kyrpides, N.C., Stepanauskas, R., Rubin, E.M., Hugenholtz, P., Woyke, T., 2013. Insights into the phylogeny and coding potential of microbial dark matter. Nature 499, 431–437. 10.1038/nature12352

106. Robeson, M.S., O’Rourke, D.R., Kaehler, B.D., Ziemski, M., Dillon, M.R., Foster, J.T., Bokulich, N.A., 2021. RESCRIPt: Reproducible sequence taxonomy reference database management. PLoS Comput Biol 17, e1009581. 10.1371/journal.pcbi.1009581

107. Ross, A.A., Doxey, A.C., Neufeld, J.D., 2017. The Skin Microbiome of Cohabiting Couples. mSystems 2. 10.1128/mSystems.00043-17

108. Saier, M.H., Reddy, V.S., Tsu, B. V., Ahmed, M.S., Li, C., Moreno-Hagelsieb, G., 2016. The Transporter Classification Database (TCDB): recent advances. Nucleic Acids Res 44, D372–D379. 10.1093/nar/gkv1103

109. Soto, W., Nishiguchi, M.K., 2021. Environmental Stress Selects for Innovations That Drive Vibrio Symbiont Diversity. Front Ecol Evol 9. 10.3389/fevo.2021.616973

110. Steinegger, M., Söding, J., 2017. MMseqs2 enables sensitive protein sequence searching for the analysis of massive data sets. Nat Biotechnol 35, 1026–1028. 10.1038/nbt.3988

111. Sun, Y., Liu, Y., Pan, J., Wang, F., Li, M., 2019. Perspectives on Cultivation Strategies of Archaea. Microb Ecol. 10.1007/s00248-019-01422-7

112. Suschek, C. V., Opländer, C., van Faassen, E.E., 2010. Non-enzymatic NO production in human skin: Effect of UVA on cutaneous NO stores. Nitric Oxide 22, 120–135. 10.1016/j.niox.2009.10.006

113. Taylor, D.L., Bruns, T.D., 1999. Community structure of ectomycorrhizal fungi in a *Pinus muricata* forest: minimal overlap between the mature forest and resistant propagule communities. Mol Ecol 8, 1837–1850. 10.1046/j.1365-294x.1999.00773.x

114. Tourna, M., Freitag, T.E., Nicol, G.W., Prosser, J.I., 2008. Growth, activity and temperature responses of ammonia-oxidizing archaea and bacteria in soil microcosms. Environ Microbiol 10, 1357–1364. 10.1111/j.1462-2920.2007.01563.x

115. Tourna, M., Stieglmeier, M., Spang, A., Könneke, M., Schintlmeister, A., Urich, T., Engel, M., Schloter, M., Wagner, M., Richter, A., Schleper, C., 2011. Nitrososphaera viennensis, an ammonia oxidizing archaeon from soil. Proc Natl Acad Sci U S A 108, 8420–8425. 10.1073/PNAS.1013488108/SUPPL_FILE/PNAS.201013488SI.PDF

116. Umbach, A.K., Stegelmeier, A.A., Neufeld, J.D., 2021. Archaea Are Rare and Uncommon Members of the Mammalian Skin Microbiome. mSystems 6. 10.1128/MSYSTEMS.00642-21/ASSET/1CFB65AC-B6A0-4252-8E75-629C6EA38AC1/ASSETS/IMAGES/LARGE/MSYSTEMS.00642-21-F004.JPG

117. Untereiner, W.A., Naveau, F.A., 1999. Molecular systematics of the Herpotrichiellaceae with an assessment of the phylogenetic positions of Exophiala dermatitidis and Phialophora americana. Mycologia 91, 67–83. 10.1080/00275514.1999.12060994

118. Usuda, D., Higashikawa, T., Hotchi, Y., Usami, K., Shimozawa, S., Tokunaga, S., Osugi, I., Katou, R., Ito, S., Yoshizawa, T., Asako, S., Mishima, K., Kondo, A., Mizuno, K., Takami, H., Komatsu, T., Oba, J., Nomura, T., Sugita, M., 2021. Exophiala dermatitidis. World J Clin Cases 9, 7963–7972. 10.12998/wjcc.v9.i27.7963

119. Vallenet, D., Calteau, A., Dubois, M., Amours, P., Bazin, A., Beuvin, M., Burlot, L., Bussell, X., Fouteau, S., Gautreau, G., Lajus, A., Langlois, J., Planel, R., Roche, D., Rollin, J., Rouy, Z., Sabatet, V., Médigue, C., 2020. MicroScope: an integrated platform for the annotation and exploration of microbial gene functions through genomic, pangenomic and metabolic comparative analysis. Nucleic Acids Res 48, D579–D589. 10.1093/NAR/GKZ926

120. van Ulsen, P., Zinner, K.M., Jong, W.S.P., Luirink, J., 2018. On display: autotransporter secretion and application. FEMS Microbiol Lett 365. 10.1093/femsle/fny165

121. Walters, W.A., Caporaso, J.G., Lauber, C.L., Berg-Lyons, D., Fierer, N., Knight, R., 2011. PrimerProspector: de novo design and taxonomic analysis of barcoded polymerase chain reaction primers. Bioinformatics 27, 1159–1161.

122. Wang, H., Bagnoud, A., Ponce-Toledo, R.I., Kerou, M., Weil, M., Schleper, C., Urich, T., 2021. Linking 16S rRNA Gene Classification to *amoA* Gene Taxonomy Reveals Environmental Distribution of Ammonia-Oxidizing Archaeal Clades in Peatland Soils. mSystems 6. 10.1128/mSystems.00546-21

123. Wood, D.E., Lu, J., Langmead, B., 2019. Improved metagenomic analysis with Kraken 2. Genome Biol 20, 257. 10.1186/s13059-019-1891-0

124. Yeats, C., Bentley, S., Bateman, A., 2003. New Knowledge from Old: In silico discovery of novel protein domains in Streptomyces coelicolor. BMC Microbiol 3, 3. 10.1186/1471-2180-3-3

125. Zhang, H., Yohe, T., Huang, L., Entwistle, S., Wu, P., Yang, Z., Busk, P.K., Xu, Y., Yin, Y., 2018. dbCAN2: a meta server for automated carbohydrate-active enzyme annotation. Nucleic Acids Res 46, W95–W101. 10.1093/nar/gky418

