## Supplementary Information for "Molecular Tracking and Cultivation Reveal Ammonia-Oxidizing Archaea as Emerging Commensals of the Human Skin Microbiome"

Alexander Mahnert*[,1], Maximilian Dreer*[,2], Ülkü Perier[2], Michael Melcher[2], Stefanie Duller[1], Adina Lehnen[1], Theodora Goessler[1], Daniela Brunner[1], Thomas Graier[3], Peter Wolf[3], Rafael I. Ponce-Toledo[2,#], Logan H. Hodgskiss[2], Melina Kerou[+,2], Christine Moissl-Eichinger[+,1,4], and Christa Schleper[+,2]

* Shared authorship

[+] Shared corresponding authorship

[1] Medical University of Graz, D&R Institute for Hygiene, Microbiology, and Environmental Medicine, Graz, Austria

[2] University of Vienna, Archaea Biology and Ecogenomics Unit, Vienna, Austria

[3] Medical University of Graz, Department of Dermatology and Venereology, Graz, Austria

[4] BioTechMed Graz, Graz, Austria

[#] Present address: Institut de Systématique, Evolution et Biodiversité (ISYEB), Sorbonne Université, CNRS, Muséum National d'Histoire Naturelle, Paris, France

### SUPPLEMENTARY METHODS

#### Enrichment process of AOA from skin inocula

Multiple approaches to selectively enrich or directly isolate skin AOA from initial enrichment cultures were tested and are listed below:

**Antibiotics**: The antibiotics used to selectively enrich for AOA in human skin samples were selected based on available information regarding their inertness against AOA. The following antibiotics from the respective antibiotic groups were used individually and in different combinations: aminoglycosides (kanamycin: 25 µg/ml), streptomycin: 10 µg/ml), carboxypenicillins (carbenicillin: 25 µg/ml), aminopenicillins (ampicillin: 25 µg/ml) (Tourna et al., 2011), aminocoumarins (novobiocin: 10 µg/ml) (Abby et al., 2018) and amphenicols (chloramphenicol dissolved in water: 10 µg/ml) (Bayer et al., 2016). While some of the above listed antibiotics led to growth retardation, none completely inhibited nitrite production of skin AOA enrichment cultures and were thus used to selectively enrich for AOA.

**Filtration**: Members of the *Ca.* Nitrosocosmicus clade are known to form aggregates of differing sizes (Jung et al., 2016; Lehtovirta-Morley et al., 2016; Liu et al., 2021, 2019). Using filters with multiple pore sizes, which are larger than the biggest observed contaminants would therefore theoretically retain voluminous AOA aggregates on the filters while removing most or all contaminants. After filtration, the filters were washed or immediately rinsed using growth medium and the resulting cell suspension was used for inoculation.

Unsuccessful approaches to reduce contamination levels included elevated $NO_2$- levels, freeze-thawing (-70°C/room temperature), dilution-to-extinction, and plating with the recently published “Liquid-Solid” method (Klein et al., 2022). Enrichment cultures using these methods resulted in very limited or no reduction of bacterial and/or fungal contaminants.

**Culture maintenance**

Enrichments are routinely cultivated in sterile 30 ml polystyrene screw cap tubes containing 18 ml FWM supplemented with 0.6 mM $NH_4Cl$, 2 mM $Na_2CO_3$ (R) or $NaHCO_3$ (T, X, Z), 20 µl non-chelated trace element solution without molybdenum ($Na_2MoO_4 \cdot 2H_2O$), 7.5 µM FeNaEDTA, 20 µl vitamin solution as well as 25 µg/ml carbenicillin and kanamycin and are transferred every two to four weeks upon reaching nitrite concentrations of approximately 450 µM. The inoculation volumes were decreased to 10% (v/v).

The medium of enrichment R additionally contains 10 mM MOPS ((3-(N-morpholino) propanesulfonic acid), titrated to pH 7 with HCl) and has a final pH of 7. Ammonium and nitrite concentrations were measured colorimetrically by the indophenol method and Griess reagent respectively, as described previously (Reyes et al., 2020; Tourna et al., 2011). Aliquots used for ammonium measurements were frozen at -20°C and measured jointly at a later time point. Cultures were incubated aerobically at 28°C without shaking in the dark. Solutions were autoclaved (FWM, $NH_4Cl$) or filter sterilized with a 0.25 µm filter (trace element solution, vitamin solution, FeNaEDTA, antibiotics) prior to use.

AOA are known to produce large amounts of cofactor $F_{420}$ (Spang et al., 2012). Therefore, autofluorescence of $F_{420}$ was used to visualize putative AOA in the enrichments (Supplementary Fig. 3, false-colored in cyan).

### SUPPLEMENTARY RESULTS AND DISCUSSION

#### Genome characterization of *Ca.* N. unguis and *Ca.* N. epidermidis

Analyses of the genome content of the novel strains reveal that they encode all 6 subunits of the archaeal ammonia monooxygenase (amoABCXYZ) (Hodgskiss et al., 2023), as well as urease and two urea transporters (UT and SSS families). Genes characteristic of the *Ca.* Nitrosocosmicus genus (we refrain from the “Ca.” prefix for better readability in the following sections) are also found, including a V-type ATPase related to tolerance in acidic environments and only the low affinity ammonia permease. The glutamate synthase (GOGAT) is encoded by *Ca.* N. unguis but not by *Ca.* N. epidermidis, indicating a possible acquisition from the common ancestor of *Ca.* N. oleophilus, *Ca.* N. arcticus, and *Ca.* N. unguis (Supplementary Data: Orthologs and Annotations).

The MAGs harbor the canonical genomic repertoire for carbon fixation, glyconeogenesis, and storage compound synthesis. Both isolates, as well as *Ca. N.* oleophilus, encode an additional acetyl-CoA/propionyl-CoA carboxylase, the key carbon fixation enzyme of the HPHB cycle, which could potentially alleviate a bottleneck in carbon fixation or assist with the carboxylation of accumulated propionyl-CoA within the skin tissue, which could otherwise have toxic effects. As all *Ca.* Nitrosocosmicus genomes, they encode arrays of membrane-bound PQQ dependent Glucose/sorbosone dehydrogenase family proteins, which could supplement electrons through the oxidation of sugars in the pseudoperiplasm.

An inspection of the CAZY database of carbohydrate-active enzymes and auxiliary proteins in *Ca.* Nitrosocosmicus reveals the extensive genomic repertoire of this genus enabling the utilization of various complex carbohydrates. Examples include multiple copies of PQQ-dependent oxidoreductases, putative cellulases, putative (oligo)xyloglycanases, and various glycosyl-transferase families. *Ca.* N. unguis encodes additionally an alpha-amylase (GH119), which would confer the ability to degrade starch, as well as a protein of the GMC (glucose-methanol-choline oxidase) oxidoreductase family (AA3) which comprises FAD-dependent enzymes participating in lignocellulose degradation. Sequence and phylogenetic analyses indicate that this enzyme belongs to the pyranose2-oxidase (POx) subfamily, present mostly in Actinobacteria, which can oxidize various monosaccharides with either $O_2$ (generating $H_2O_2$), (substituted) quinones or complexed metal ions as electron acceptors. While the proposed metabolic role of this enzyme is to supply lignolytic peroxidases with $H_2O_2$, it was also proposed that it could detoxify semiquinones, phenolic radicals, or reactive metal complexes and thus prevent cellular damage (Herzog et al., 2019; Sützl et al., 2018). In either case, the generation and modulation of ROS on the skin can have important implications for skin homeostasis (Ni et al., 2022). GMC oxidoreductases have been implicated in establishing successful plant-bacterial and plant-insect symbioses, mostly by virtue of their roles in ROS detoxification and iron homeostasis (Zou et al., 2020).

Both skin isolates encode the basic repertoire for detoxifying oxidative stress, such as superoxide dismutase, Mn-catalase, alkyl hydroperoxide reductases (AhpC), and thioredoxin systems. All characterized *Ca.* Nitrosocosmicus isolates except *Ca. N.*

agrestris and *Ca.* N. epidermidis encode additionally a DyP-type peroxidase, a heme peroxidase with optimal activity at low pH (4-5), which intriguingly can be packaged in encapsulin nanocompartments essential for oxidative stress resistance and virulence in *Mycobacterium tuberculosis* (Lien et al., 2021). Although no encapsulin homologs were found in AOA, the role of this enzyme in AOA physiology remains to be elucidated. Curiously, *Ca.* N. unguis does not encode a disulfide bond oxidoreductase D family protein (DsbD), raising the question of repairing extracellular protein damage from ROS. In addition, though, all *Ca.* Nitrosocosmicus, different from all other AOA, encode a membrane-associated putative Alkyl hydroperoxide reductase F homolog, which would enable the faster regeneration of AhpC homologs.

Both skin isolates encode large and small conductance mechanosensitive ion channels (MscL, MscS), showing a patchy distribution in other genomes of the lineage. Among AOA, only NS-zeta genomes encode homologs of the K+-selective channel in endosomes and lysosomes (KEL) family (1.A.78), with *Ca.* N. unguis and *Ca.* N. oleophilu*s* encoding 5 homologs. Both skin strains encode two families of low affinity/high yield Trk-type K+ transporters in addition to a Trk-type transporter family shared by all NS-zeta strains, with a total of 4 genes per strain, indicating their capacity to deal with changes in the osmotic environment.

The skin strains share only 20 protein families, among which we can identify two families of low affinity/high yield Trk-type K+ transporters, one $F_{420}$ $H_2$-dependent quinone reductase and a collagen triple-repeat containing protein (such domains are also encoded by other AOA). Among the shell proteome of each strain, we find extracellular peptidases, glycosyl transferases, and cell surface-related proteins hypothesized to enable the adaptation to a skin environment presenting challenges such as desiccation and increased sodium concentrations. In *Ca.* N. unguis, we find a specific transporter of the bile acid/$Na^+$ symporter (BASS) family (2.A.28), with hits to characterized pyruvate: $Na^+$ transporters, an extracellular alpha-amylase family protein, and a putative muconolactone isomerase. The latter participates in the catabolism of catechol to succinate- and acetyl-CoA in the beta-ketoadipate pathway, while a putative catechol 2,3-dioxygenase is also encoded by the genomes of *Ca.* Nitrosocosmicus. Catechol is a phenolic compound occurring naturally in fruits and vegetables, while also used in the production of insecticides, perfumes, drugs, and other chemicals such as antiseptics. It can be degraded by soil bacteria and fungi, raising the interesting possibility of AOA participating in this pathway and supplementing carbon compounds for anabolic purposes.

*Ca.* N. unguis encodes two copies of the DNA repair enzyme alpha-ketoglutarate-dependent dioxygenase alkB, present only in *Ca.* N. oleophilus and *Ca.* N. agrestis among AOA. This enzyme can remove methyl adducts and larger alkylation lesions from DNA bases. This common type of DNA damage can be caused by either environmental agents such as alkylating anticancer drugs (typically nitrosamines) or endogenous compounds. Both skin strains encode in addition the full repertoire of DNA repair systems found in AOA (BER, NER, Uvr, PolY) including the non-homologous end-joining system found only in *Ca.* Nitrosocosmicus (Abby et al., 2020).

### *Ca.* Nitrosocosmicus representatives are uniquely equipped for the colonization of host surfaces

Rather than encoding specific adaptations, genomic investigations of this clade indicate its suitability for colonization of animal and plant hosts. *Ca.* N. franklandianus, *Ca.* N. agrestis, and the two skin strains encode three families of ~200aa proteins with significant hits to the Pore-forming RTX Toxin (RTX-toxin) Family (1.C.11), containing serralysin-like metalloprotease C-terminal domains and RTX calcium-binding nonapeptide repeats. These are absent from other AOA, which encode at most two longer proteins with the same domains. These extracellular putative Zn-dependent endopeptidases are encoded in clusters of 3 genes in these genomes (2 clusters in *Ca.* N. epidermidis), and in Gram-negative bacteria are frequently implicated in virulence, as they are able to cause extensive tissue damage, or affect surface attachment (Garcia et al., 2018; Mondal et al., 2022). In general, *Ca.* Nitrosocosmicus genomes are enriched in various families of peptidases compared to other AOA lineages, with the skin isolates complement consistently at the higher end of the distribution for each family (e.g. peptidase family M23, Subtilisin-like serine protease). Secreted proteases by the skin microbiota can have diverse roles both in tissue invasion and establishment but also in modulation of other virulence factors (Chua et al., 2022).

One of the most widespread mechanisms during bacterial colonization/infection of host epithelia is the expression of various kinds of lipases or esterases which enable either the catabolism of host fatty acids in order to alter host immune response or utilize them for anabolic purposes (Chen and Alonzo, 2019; Kumar et al., 2021). *Ca.* Nitrosocosmicus

AOA encode specifically a family of predicted extracellular/membrane-attached putative triacylglycerol esterases/lipases (arCOG06923, PF02089), of which the skin strains encode 2-3 orthologs. Moreover, *Ca.* N. unguis (together with *Ca.* N. oleophilus) is enriched in patatin-like phospholipase domain proteins (arCOG09020). While it is unclear whether these genes participate in similar functions during colonization of the skin environment by AOA, the similarity to known epithelial colonizing strains is intriguing (Chen and Alonzo, 2019; Kumar et al., 2021).

*Ca.* Nitrosocosmicus isolates have been shown to produce rich biofilms and exhibit superior adhesion capacities compared to other AOA lineages, a trait observed for almost all microbial species residing on the human skin (Brandwein et al., 2016). This is mirrored in their proteomes which are enriched in acetamidosugar biosynthesis and adhesion-related proteins, as well as being the only AOA clade encoding LPSE family lipopolysaccharide exporters (Supplementary Data: Annotations) (Jung et al., 2016; Liu et al., 2021). For example, *Ca.* Nitrosocosmicus genomes specifically encode one family of extracellular serine-rich repeat proteins with roles including adhesion and biofilm formation, with 35% sequence identity to dentin sialoprotein-like proteins from eukaryotes (Cinar et al., 2024) (Supplementary Data: Annotations).

**Alpha and beta diversity analyses of cross-sectional cohort A**

47 subjects were recruited for cohort A, which comprised 21 healthy individuals in the age of 20-40 years (cohort A1) as well as 26 healthy individuals aged 60-80+ years (cohort A2). From each subject eight defined body sites (decollete, forehead, crook, arm, back, armpit, hands, and part) were sampled and skin physiological parameters (skin health,

pH, skin fat, wetness, and temperature) were recorded (Supplementary Fig. 7 and Supplementary Tables S3 to S6). Each subject provided a comprehensive questionnaire, which led to a rich collection of categorical and numerical metadata covering demographic, health, and lifestyle details. All samples were subjected to bacterial qPCR and ammonia-oxidizing archaea (AOA) specific amoA-qPCR, as well as 16S rRNA gene profiling with three different primer combinations to cover all three domains of life and more specifically archaea and AOA (details see materials & methods). This endeavor resulted in a total of 1,770 amplicons (PCR reactions (16S rRNA gene, *amoA*) per sample including controls). (Supplementary Fig. 2 STORM chart)

Our results support observations described in (Moissl-Eichinger et al., 2017). Reclassification of representative 16S rRNA gene sequences from (Moissl-Eichinger et al., 2017; Probst et al., 2013) with the Silva database (v.138) revealed that most archaeal sequences from human skin belong to *Ca.* Nitrosocosmicus and unknown genera of the Nitrososphaeraceae family (Supplementary Fig. 8).

Hence, older subjects showed a higher diversity of archaea on their skin (highest archaeal diversity was determined for samples from the arm and reached a maximum of microbial diversity (Shannon index H' ~ 3). According to weighted UniFrac metrics, the largest significant dissimilarities were visible for samples of the arm, part, and back. The composition of archaeal communities was also significantly different between younger and older subjects (weighted UniFrac, PERMANOVA q-value = 0.004). An unidentified archaeon of the family Nitrososphaeraceae was significantly differentially abundant (ANCOM W = 18 and 29) and showed highest proportions on the arms of elderly subjects. Extending this analysis to bacterial and fungal diversity, significant differences (Kruskal-

Wallis tests) were evident between subjects (P = 0.01) and different body sites (P = 7.3E-18). Significant positive Spearman rank correlations could be drawn for the skin sebum content (P = 0.01), meat consumption (P = 0.0003), as well as a subject's body size (P = 0.004) and BMI (P = 0.04). Hence, while significant differences per age group (A1 and A2, see Supplementary Table S3 and S4) were only evident for body size (q < 0.001), body weight (q = 0.004), skin pH group (q = 0.004) and skin moisture (Corneometer: q < 0.001), sampled body sites showed significant differences for all recorded skin physiological parameters (p < 0.001 for skin health, skin pH, skin fat, skin wetness and skin temperature (see Supplementary Table S5). On the contrary microbial alpha diversity was not significantly different between age groups (A1 and A2, see Supplementary Table S19), but showed significant differences between sampled body sites for each metric (q < 0.001 for Shannon diversity, Faith's phylogenetic diversity, richness and Pielou's evenness (see Supplementary Table S20). Also on beta diversity level, significant weighted UniFrac dissimilarities were visible between samples from the armpit and back compared to other body sites as well as between younger and older subjects (PERMANOVA q-value = 0.015). Finally, sequences assigned to *Micrococcus*, *Cutibacterium* (particularly on the back), *Lawsonella* (most of all from the part), *Enhydrobacter* (especially on subject's arms), *Methanobrevibacter*, *Ralstonia* were significantly differentially abundant on younger subjects except *Paracoccus* (according to ANCOM W = 325 to 297).

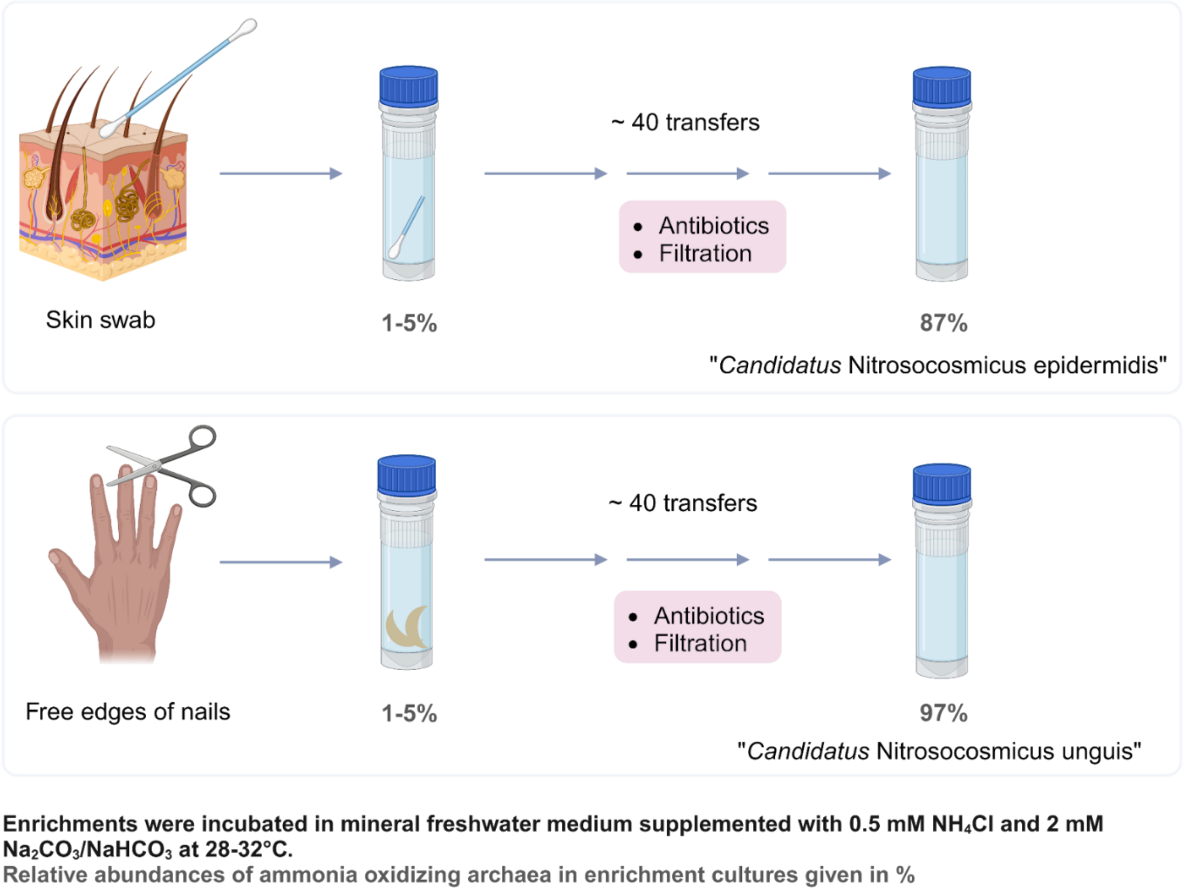


**Supplementary Fig. 1:** Enrichment scheme for skin-residing AOA. Created with BioRender.com.

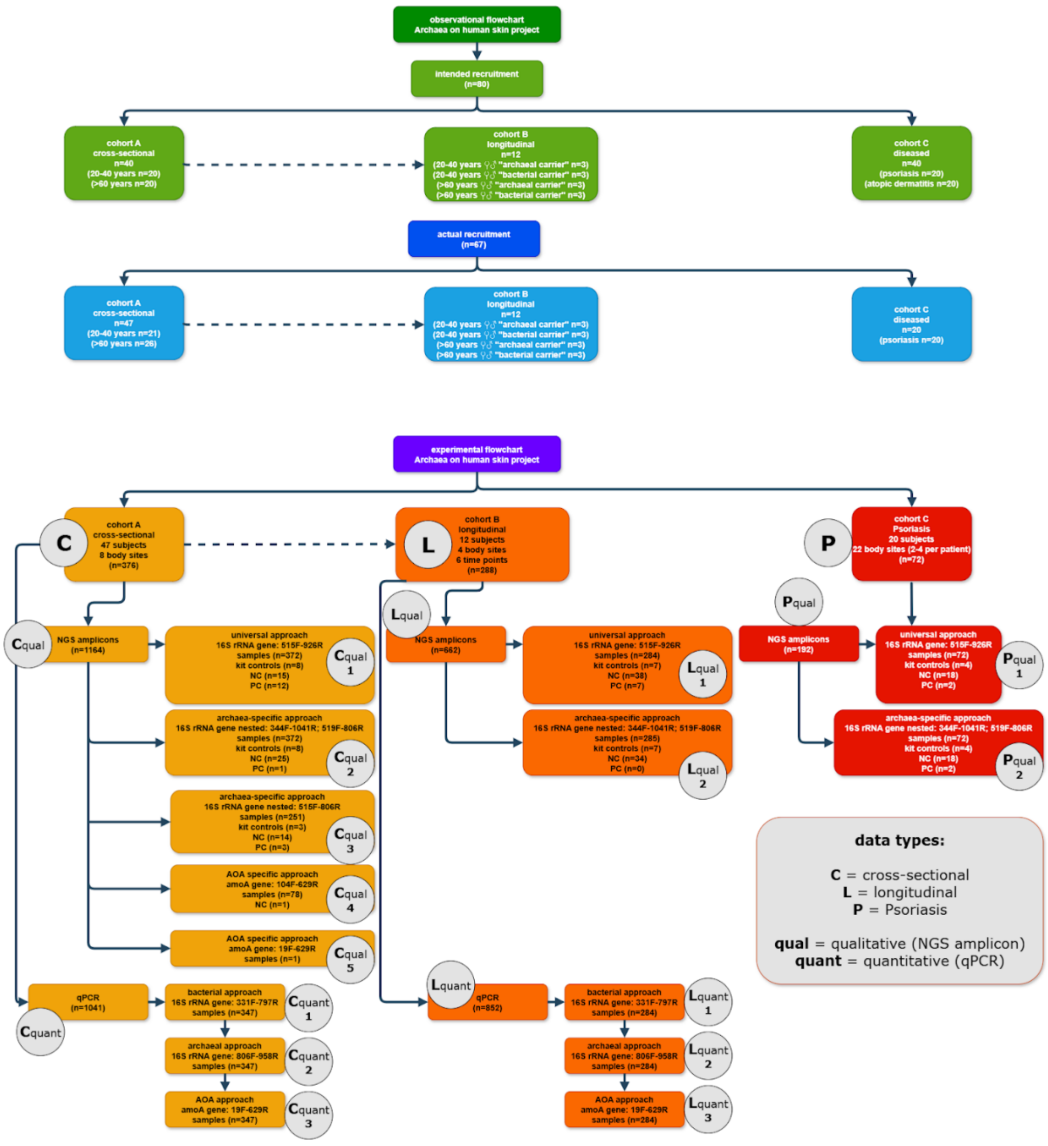


**Supplementary Figure 2** STORM chart: STORM flowchart of the project with identifiers for each different data type.

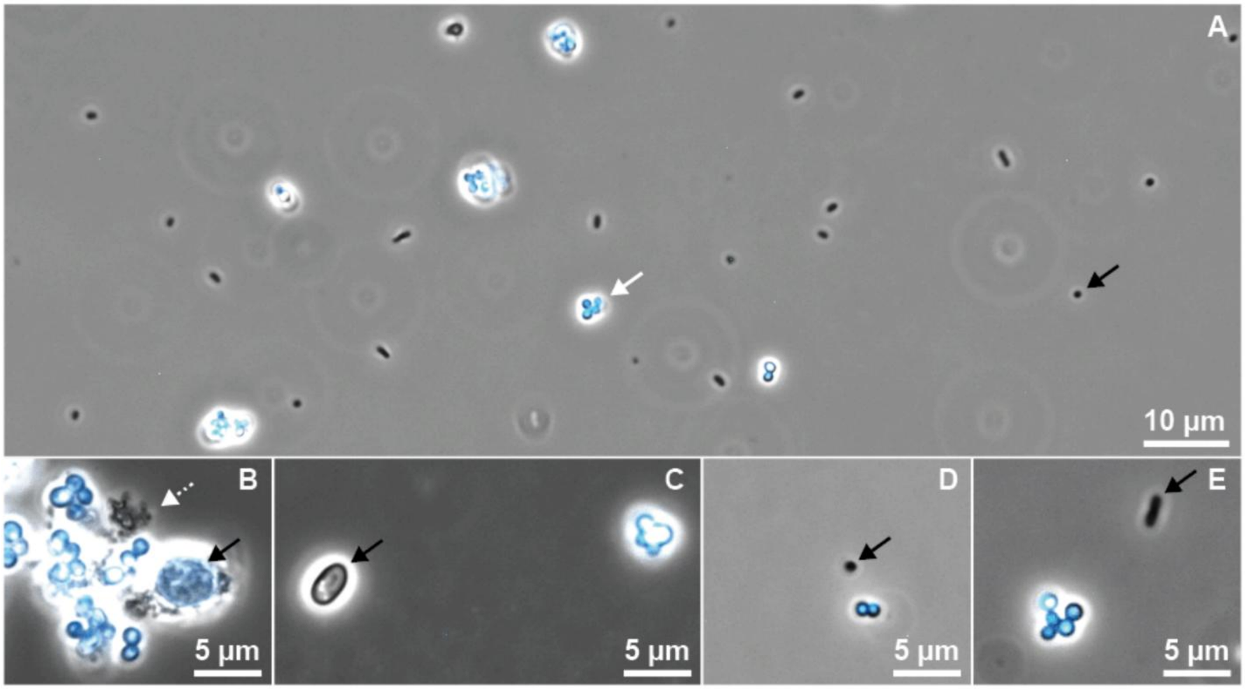


**Supplementary Fig. 3:** Identification of AOA based on F420 autofluorescence by light microscopy.

(A, E) Enrichment Z3A. (B) Enrichment R2S. (C) Enrichment T1S. (D) Enrichment X2B. The autofluorescence of AOA can be used to distinguish between AOA and other contaminants in complex enrichment cultures (A). Putative ammonia-oxidizing archaea were visualized by F420 autofluorescence false-colored in cyan (white arrow (A), cyan (B-E)). Contaminants were not autofluorescent (black arrows), except for minimal autofluorescence of the contaminants associated with aggregates of AOA of enrichment R2S (B, black arrow). The dashed white arrow in (B) indicates putative extracellular polymeric substances (EPS). Images are overlays of phase contrast light microscopy and false-colored F420 fluorescence microscopy using a standard DAPI filter cube. Enrichment Z3A was concentrated 10X times via centrifugation before imaging.

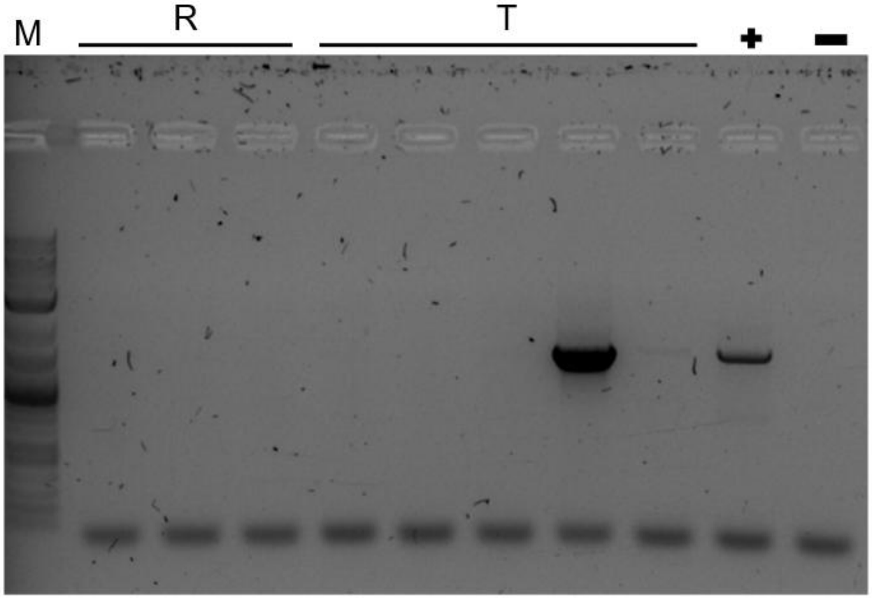


**Supplementary Fig. 4.** Agarose gel of bacterial 16S rRNA gene PCR of enrichments R2S and T1S. Multiple cultures of enrichments R and T were screened for the presence of bacterial contaminants via amplification of bacterial 16S rRNA genes (PCR, 33 cycles). Cultures free of bacterial contaminants were used as inoculum for future experiments. M: Marker, GeneRuler 1kb plus, +: positive control, -: negative control

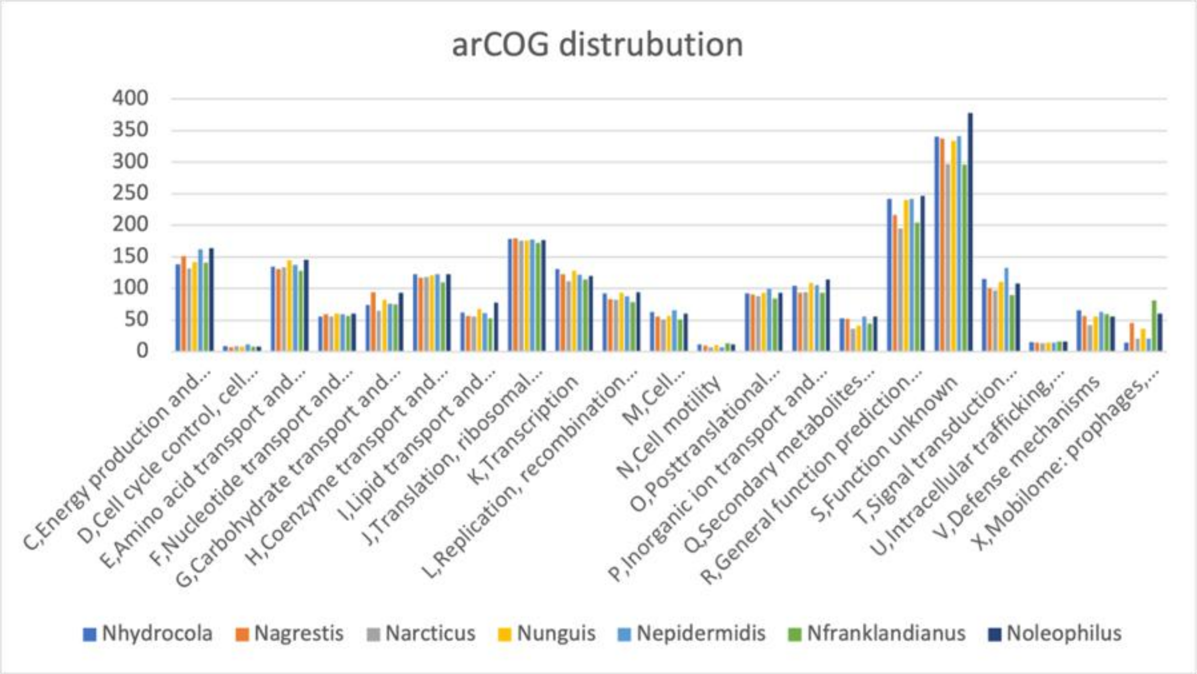

**Supplementary Fig. 5**: Distribution of arCOG functional categories among seven representative genomes of *Ca.* Nitrosocosmicus

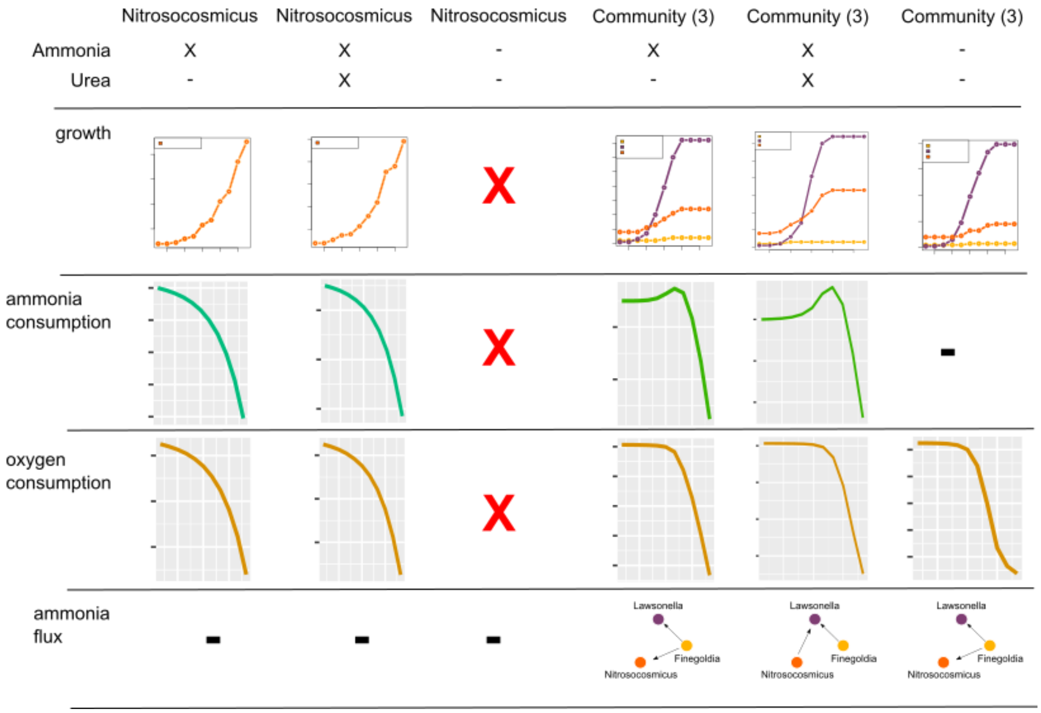


**Supplementary Fig. 6**: In silico metabolic modeling with BacArena showing modeled growth, ammonia consumption, oxygen consumption, and ammonia flux for *Ca.* Nitrosocosmicus alone and the co-occurring community of *Ca.* Nitrosocosmicus with Lawsonella and Finegoldia (see Fig. 5) with and without ammonia and urea as supplements in the provided media.

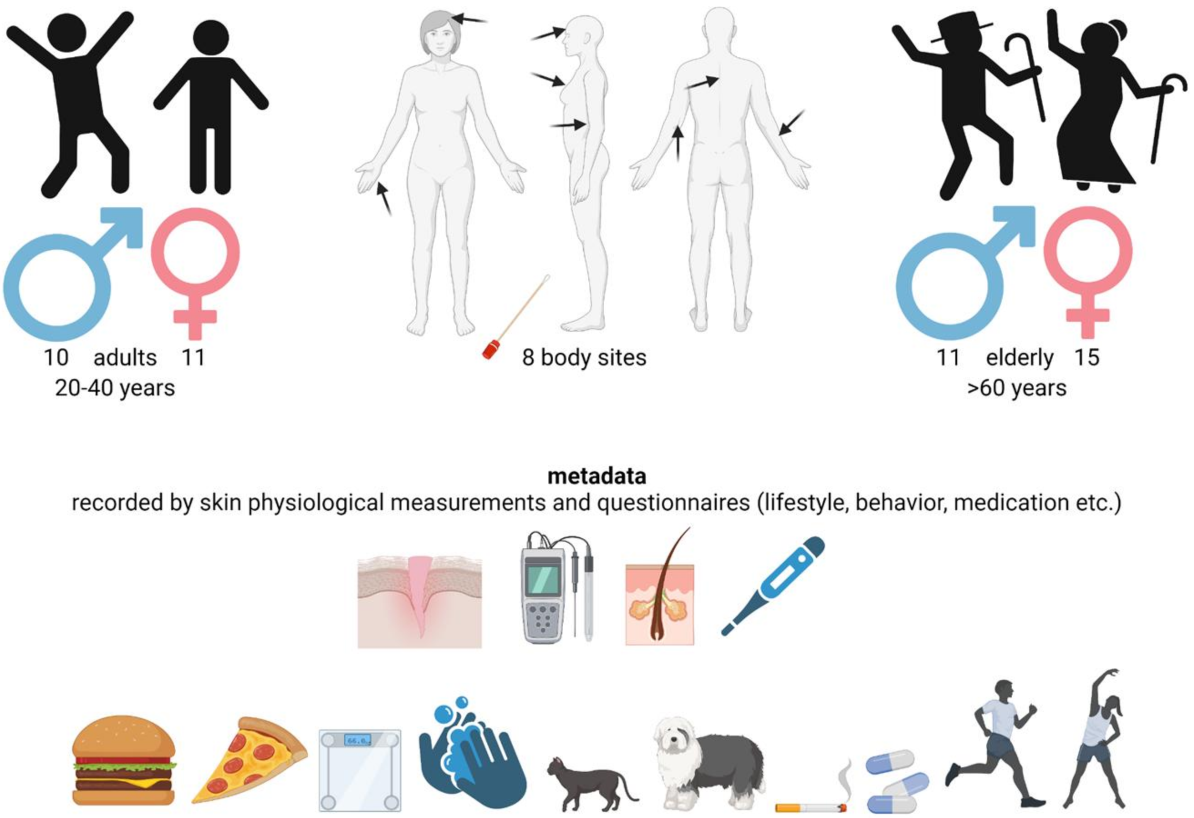


**Supplementary Fig. 7:** Sketch summarizing details about the recruited cohort A covering data types Cqual1 - 5, Cquant1 - 3 (including from top to bottom information about demographics, sampled body sites, and measured as well as recorded metadata.

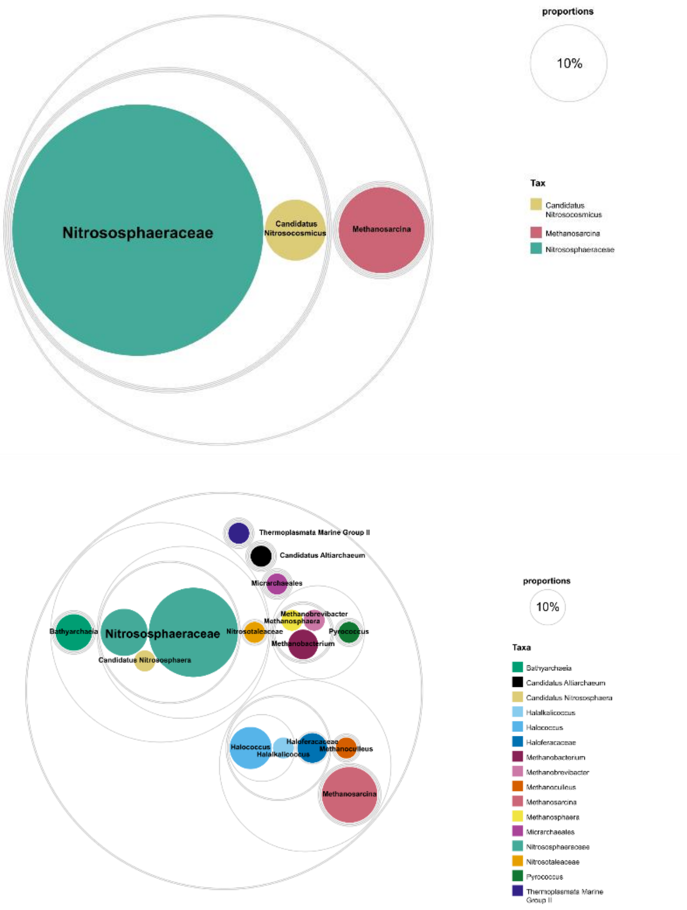


**Supplementary Fig. 8**: Circle packing plot according to the re-classification of archaeal representative sequences of [26] (n = 20, top) and [25] (n = 51, bottom).

### The skin archaeome is longitudinally persistent, while bacterial communities follow a seasonal pattern

From our initial cohort, we selected human archaea and bacteria carriers (characterized by high and low loads of archaea), balanced for sex and age (27-77 years old) and followed the skin microbiome of each individual on four selected body sites (decollete, forehead, crook, and back; due to insights obtained from cohort A) for an entire year (Fig.

4, Supplementary Fig. 9; total number of processed samples 1,230). Besides one subject, all participants were positive for archaea. Hence, also most “non-archaeal carriers” became archaea positive, and none of the ‘archaeal carriers' completely lost their archaeome during our year of frequent microbial monitoring. Signatures assigned to *Methanobrevibacter smithii* (global mean = 0.27; global variance = 0.15; importance = 0.44), *Ca.* Nitrosocosmicus (global mean = 0.17; global variance = 0.09; importance = 0.17) and other members of the *Nitrososphaera* family (global mean = 0.33; global variance = 0.15; importance = 0.12) achieved highest importance in linear regressions over time and also showed highest relative proportions of all archaea (Supplementary Fig. 10, Supplementary Fig. 11 and Supplementary Table S17). Signatures assigned to *Ca.* Nitrosocosmicus were consistently present over the entire sampling period, and only one subject was never positive for *Ca.* Nitrosocosmicus, but all subjects were positive for *M. smithii.*

In contrast to the skin archaeome (Supplementary Fig. 12) the bacterial communities followed a seasonal pattern, where higher proportions of common skin bacteria (*Cutibacterium, Corynebacterium, Staphylococcus*) could be retrieved during the colder seasons of the year, while environmental bacterial species (*Burkholderia* and *Allorhizobium*) were more prominent in the warmer seasons of the year. This observation was supported by feature volatility analysis (Supplementary Table S21), where *Burkholderia* (global mean = 0.28; global variance = 0.11; importance = 0.36) and *Allorhizobium* (global mean = 0.008; global variance = 0.00013; importance = 0.14) showed the highest seasonal dynamics in contrast to consistent patterns of typical skin bacteria like *Cutibacterium* (global mean = 0.21; global variance = 0.045; importance =

0.07), *Corynebacterium* (global mean = 0.1; global variance = 0.02; importance = 0.02), and *Staphylococcus* (global mean = 0.12; global variance = 0.024; importance = 0.04). While *Cutibacterium* was mainly recovered from back and forehead samples of younger male subjects with normal skin, *Burkholderia* was more present on samples from the arms and dry skin regions. In addition, higher proportions of *Ca.* Nitrosocosmicus were detected on male subjects. *Ca. Nitrosocosmicus* was strongly associated with normal and alkaline skin, while *M. smithii* could be also retrieved from acidic sites (Supplementary Fig. 13). Similar opposed patterns were observed for the most abundant bacterial taxa. Here, *Burkholderia* preferred acidic sites and *Cutibacterium* thrived on alkaline body sites.

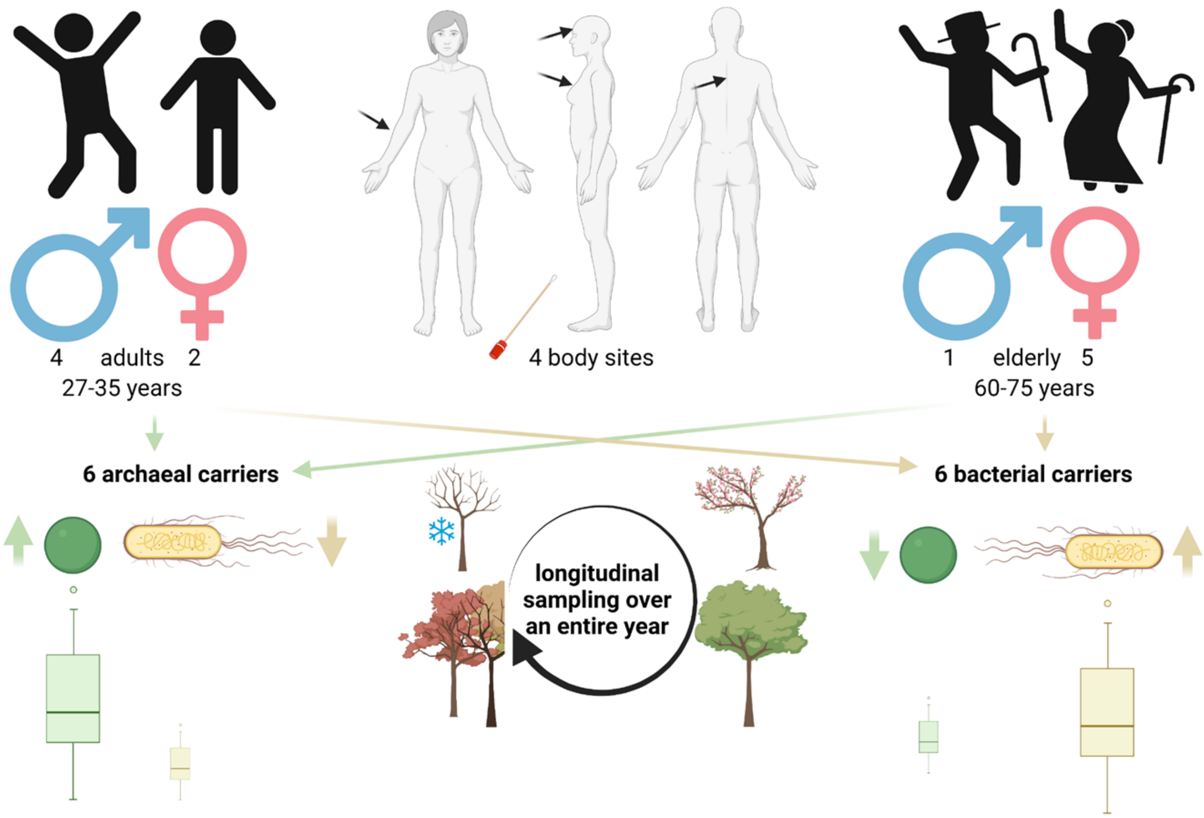

**Supplementary Fig. 9:** Sketch summarizing details about the recruited longitudinal cohort B covering data types Lqual1 - 2, and Lquant 1 - 3 (including from top to bottom information about demographics, sampled body sites, and our concept of archaeal (green) vs. bacterial (yellow) carriers.

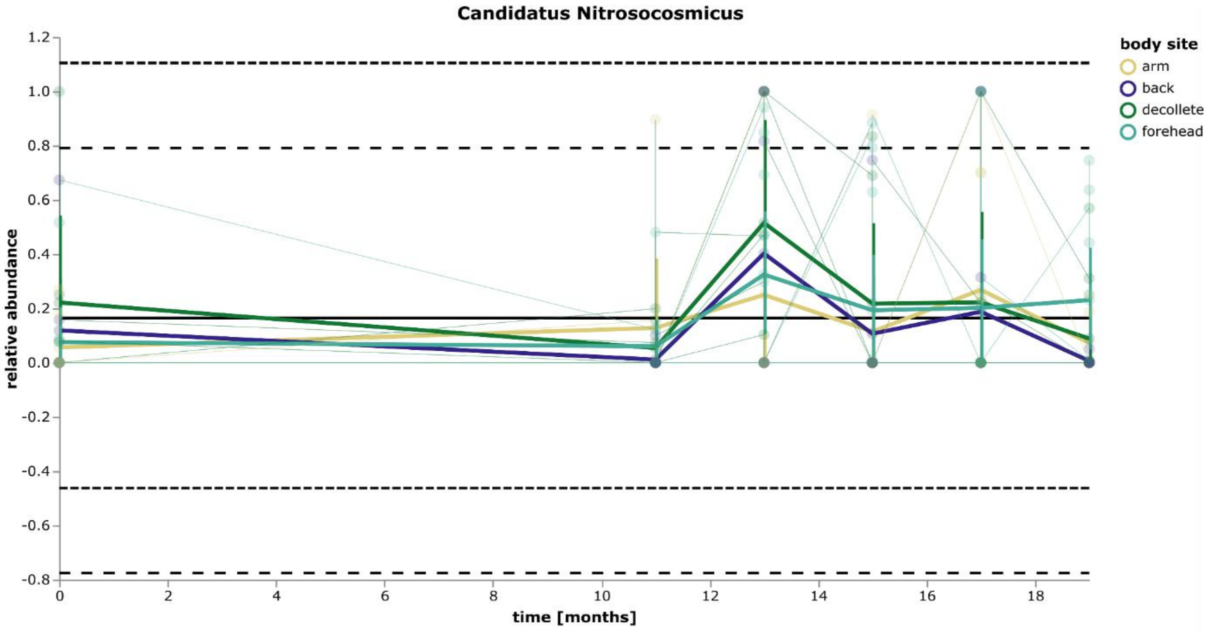


**Supplementary Fig. 10:** Volatility (spaghetti) plot of the most prevalent AOA - *Ca.* Nitrosocosmicus - on different body sites during the sampled timeframe. (n = 285; data type: Lqual2). Shown are relative abundances on genus level from archaea-specific 16S rRNA gene amplicons. The global mean is represented by a solid black horizontal line. The global control limits (+/- 2x and 3x standard deviations from the global mean) are indicated by dashed black horizontal lines. All data points are connected by colored horizontal lines grouped per body site. A thicker horizontal line shows the mean value per body site. Vertical colored lines give the standard deviation of the mean.

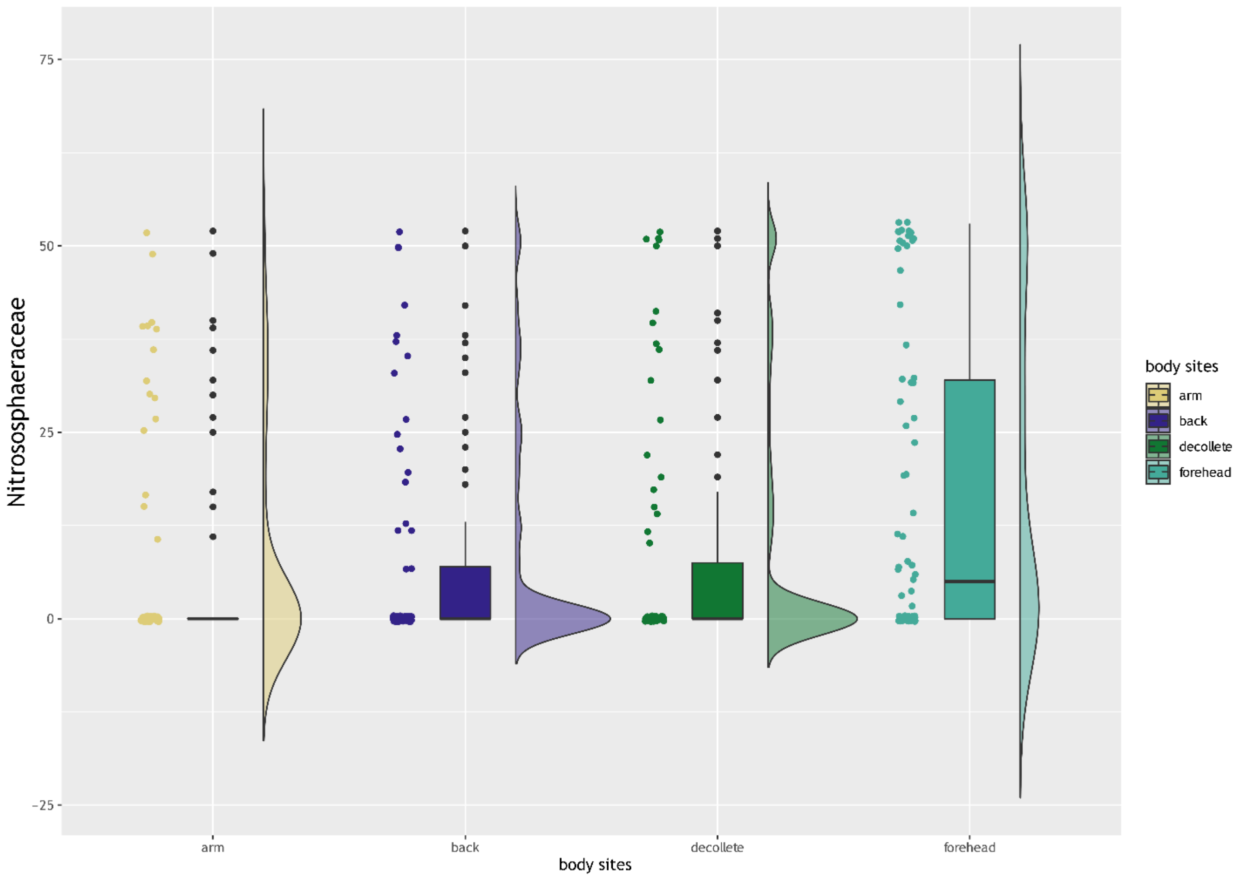


**Supplementary Fig. 11:** Raincloud plot showing significantly higher abundance of signatures from Nitrososphaeraceae on the forehead of sampled subject's (according to MaAsLin2: FDR: 7.108E−04, coefficient: −4.31E−01, value: arm; FDR: 3.353E−03, coefficient: −3.81E−01, value: back; FDR: 4.682E−03, coefficient: −3.68E−01, value: decollete; n = 285; data type: Lqual2).

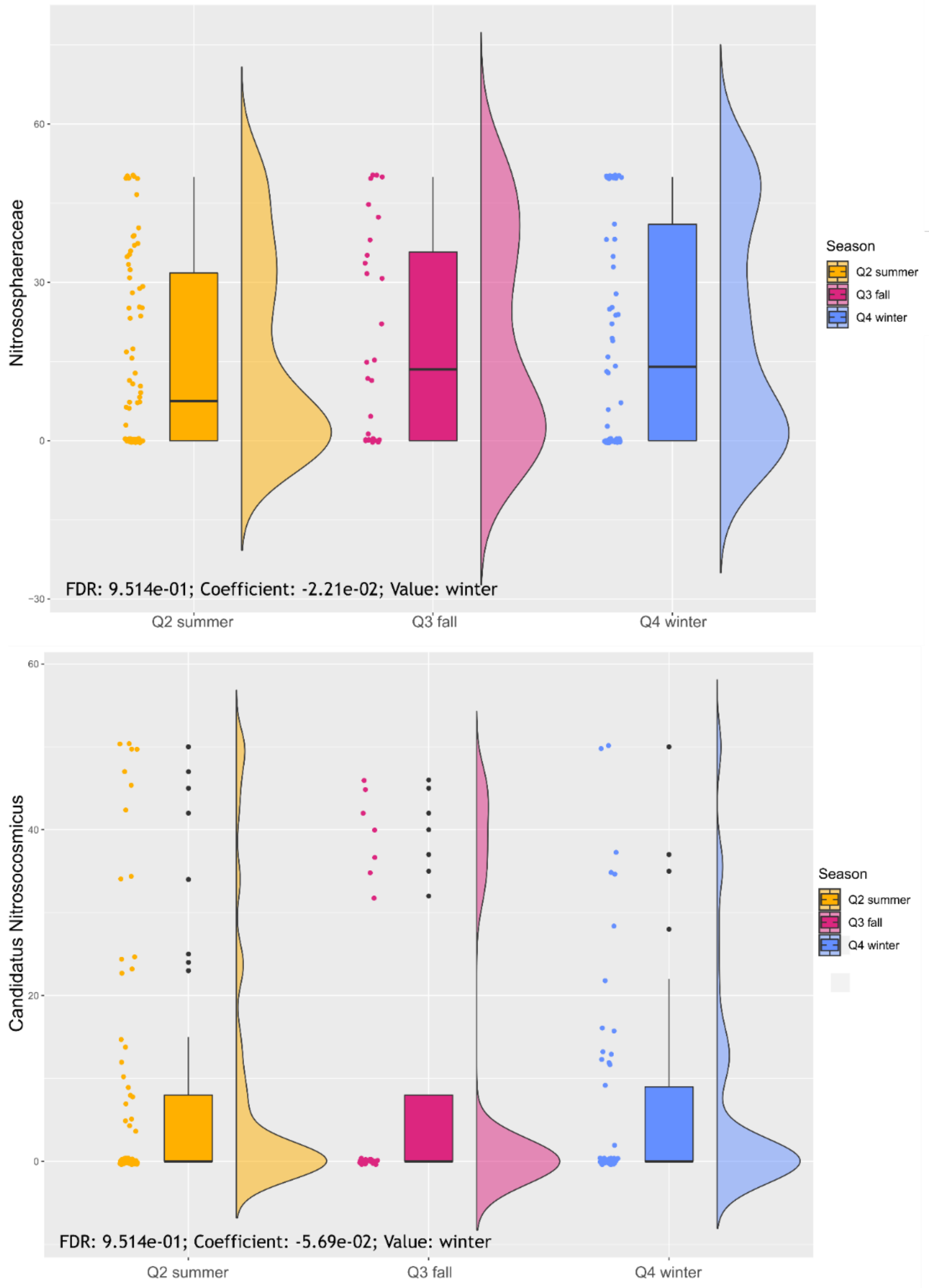


**Supplementary Fig. 12:** Raincloud plot showing dynamics of most prevalent AOA taxa throughout different sampling seasons (FDR corrected q-values from MaAsLin2; n = 285; data type: Lqual2).

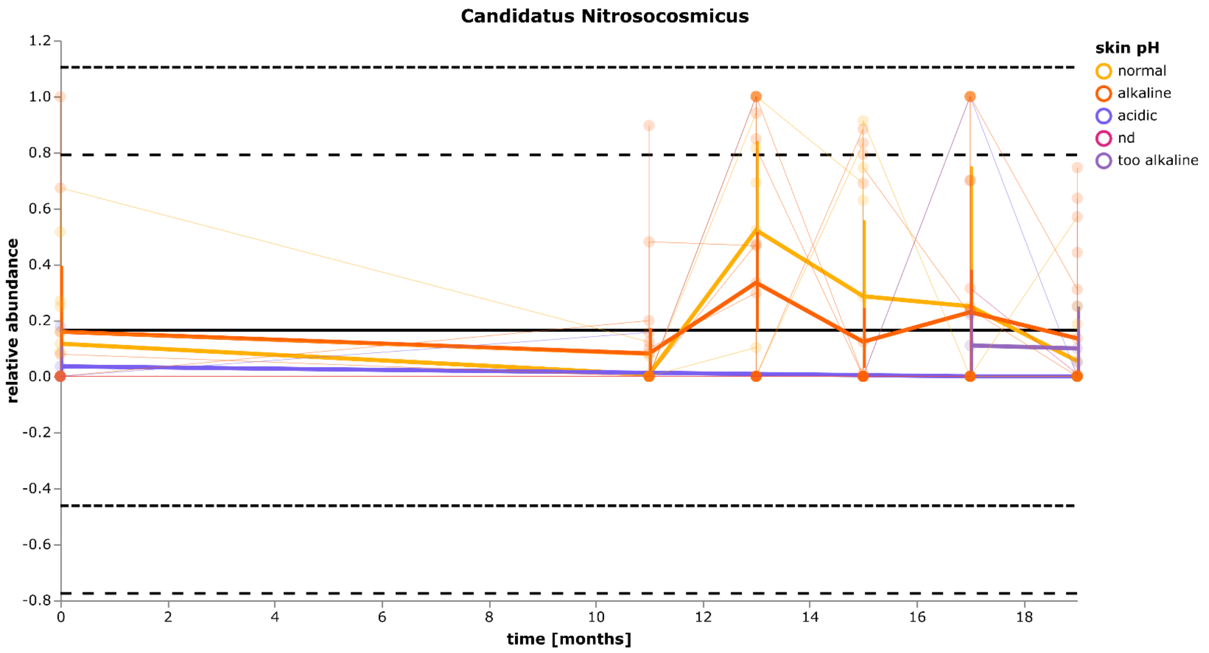


**Supplementary Fig. 13:** Volatility (spaghetti) plot showing the diverse pattern according to skin pH of *Ca.* Nitrosocosmicus over the sampled timeframe (n = 285; data type: Lqual2). Shown are relative abundances on genus level from archaea-specific 16S rRNA gene amplicons. The global mean is represented by a solid black horizontal line. The global control limits (+/- 2x and 3x standard deviations from global mean) are indicated by dashed black horizontal lines. All data points are connected by colored horizontal lines grouped per skin pH category. A thicker horizontal line shows the mean value per skin pH category. Vertical colored lines give the standard deviation of the mean.

### Proportions and abundances of AOA in comparison to bacteria were longitudinally consistent, but varying with a subject's age

Opposed age-dependent patterns were visible in linear mixed-effect models using MaAsLin2 and the QIIME2 plugin "linear-mixed-effects modeling" (Seabold & Perktold, 2010). Hence, while Nitrososphaeraceae (q-value = 0.056) and *Brevibacterium* (q-value

= 2.66*10E-11) showed positive correlations with increasing age, *Cutibacterium* (q-value = 3.78*10E-11) was significantly negatively correlated (Supplementary Figure 14). Our observation supports the common concept that not only the skin microbiome but also the skin archaeome, is shaped by physiological characteristics of diverse microhabitats on the body surface as well as to a minor extent by specific traits of the individual - for instance, a subject's age.

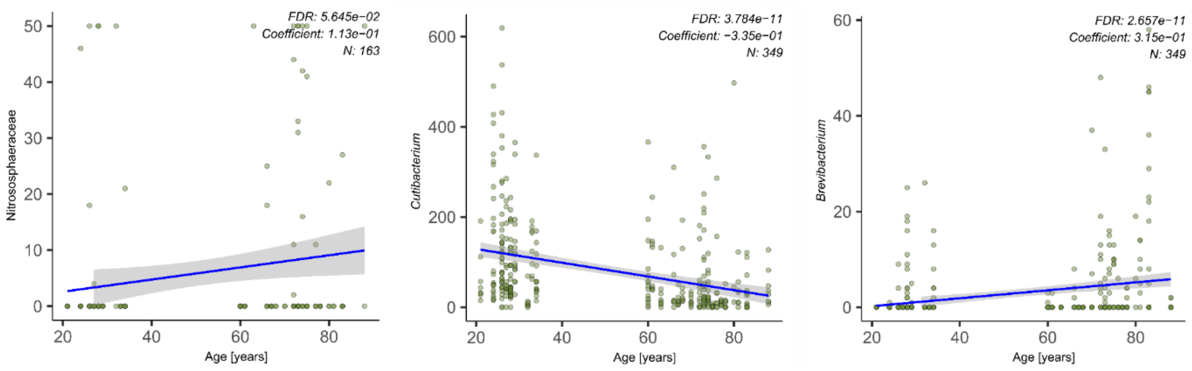


**Supplementary Fig. 14:** Correlation of selected microbial taxa with a subject's age and associations with sampled body sites (according to MaAsLin2; n = 284, data types: left Lqual2; center and right Lqual1).

We conducted quantitative PCR for bacteria (16S rRNA gene) and ammonia-oxidizing archaea (AOA, amoA gene) to integrate observations from microbial and AOA diversity also on a quantitative level. When ln transformed gene copy numbers per 300cm$^2$ of skin surface were plotted along the age of our participants in a volatility analysis, highest proportions of bacteria (21.65 on average) could be retrieved from the armpit (higher pH, higher Tewameter counts, lower sebum content), while highest proportions of AOA (10.38

on average) were located on samples from the part and forehead (lower pH, lower Tewameter counts, higher sebum content) (see Supplementary Fig. 15).

Furthermore, from our longitudinal analysis, consistently higher loads of AOA and bacteria were retrieved from skin samples of the forehead (up to 10.44 on average and 4 logs higher than samples from other body sites for AOA and up to 21.34 on average and 3 logs higher than samples from other body sites for bacterial abundances) (see Supplementary Fig. 15). This observation coincided longitudinally with a normal skin health (Tewameter measurements) and fat content (Sebumeter measurements) on acidic skin sites. However, while AOA showed higher abundances on young male subjects, bacterial abundances were comparable between different sexes and age groups.

In contrast, to the 16S rRNA gene profile, the *amoA* gene profile showed no significant differences between the two age groups (20 - 40 and 60 - 80+ years), but significant differences of amoA gene copy numbers between female and male subjects (P = 2.9*10E-3). Highest copy numbers of the amoA gene (up to 4.11*10E4 per 300 cm2) were retrieved from the head (part and forehead).

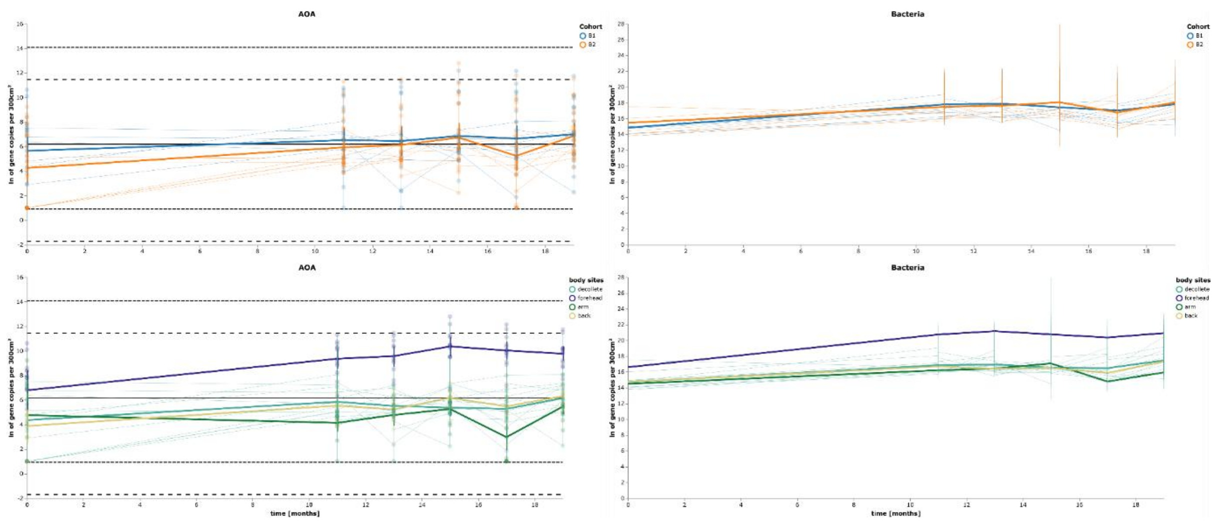

**Supplementary Fig. 15:** Volatility (spaghetti) plots of AOA (amoA gene) qPCR and bacterial (16S rRNA gene) qPCR shown per age group and per body site over one year. qPCR counts were normalized for each run and per 300cm$^2$ sampled skin surface and ln transformed for better comparability (n = 284, data type: Lquant1 and Lquant3).

### Prevalence of skin AOAs does not correlate with AOB or NOB.

We compared the prevalence of ammonia-oxidizing archaea (AOA) with the prevalence of ammonia-oxidizing bacteria (AOB) and nitrite-oxidizing bacteria (NOB) in all our samples (cohort A, B, and C; n=681). We could not detect any significant correlations between AOA, AOB or NOB. The overall prevalence for AOA was 3%, AOB 3%, and NOB 1%. The group of AOA comprised 7 genera (*Ca.* Nitrosocosmicus within Nitrososphaeraceae and *Nitrosotenuis* within Nitropumilaceae among other not higher resolved AOAs). AOB included 8 genera, which were all assigned to the family Nitrosomonadaceae and NOB covered 4 genera (*Leptospirillum*, *Nitrospira*, *Nitrotoga,* and *Omnitrophus*).

AOA, AOB, and NOB seem to prefer different niches on human skin since they only co-occurred in 1 out of 681 samples respectively (0.15% prevalence). With the archaea-specific 16S rRNA gene amplicon approach the co-occurrence of AOA and AOB or NOB increased only slightly (AOA with AOB, 1% prevalence; AOA with NOB, 0.6% prevalence) (Supplementary Tables S22 to S24).

Based on these observations, we conclude that AOA are stable inhabitants of skin samples. Their appearance is niche-specific and does not follow the pattern of AOB (Supplementary Fig. 16).

Significant positive associations (see Supplementary Tables S25 and S26) could be observed between the genus Nitrosomonadaceae_966-1 and the usage of skin creams (q = 1.88E-04, beta +0.27), consumption of meat (q = 7.9E-05, beta +0.02), physical activity (q = 9.12E-04, beta +0.01), outdoor activities (q = 3.43E-02, beta +0.004) or the food intake per kcal (q = 8.39E-03, beta 0.0003), while significant negative associations of this genus were observed for a subject's body weight (q = 3.33E-02, beta -0.01), consumption of beer (q = 2.4E-03, beta -0.06), contact to animals (q = 1.54E-04, beta -0.19), antimicrobial therapy (q = 9.39E-05, beta -0.26), or the number of meals (q = 1.9E-04, beta -0.34). In addition, a genus assigned as uncultured Nitrosomonadaceae bacterium showed a significant negative association with Faith's phylogenetic diversity (q = 5.48E-05, beta -0.005).

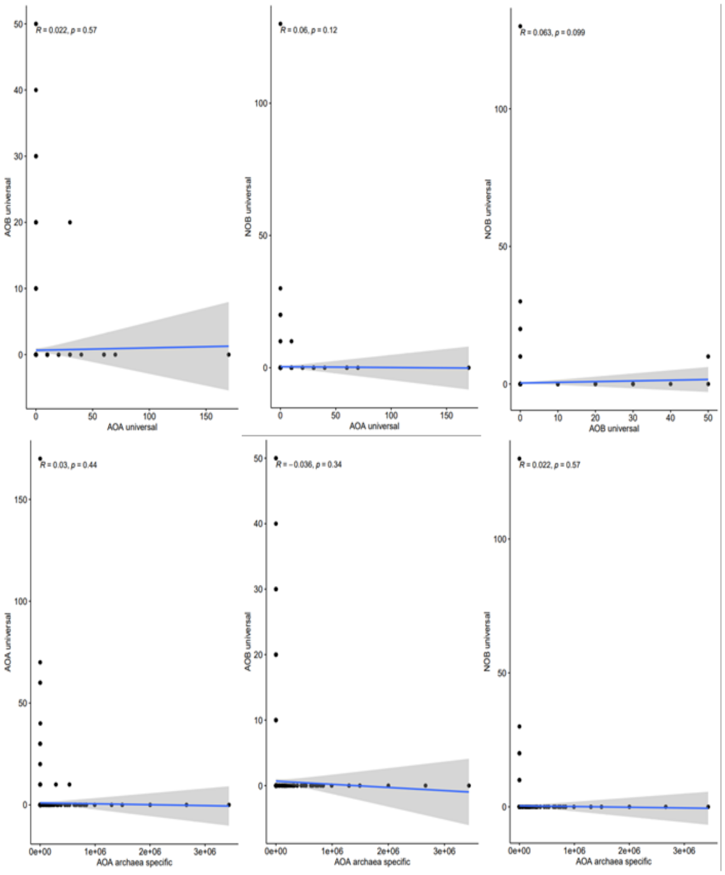


**Supplementary Fig. 16:** Scatterplots of AOA archaea-specific and non-specific vs. AOB and NOB. Potential correlations based on Spearman-rank correlations are highlighted at the top of each plot (n = 681, data types: Cqual1, Cqual2, Lqual1, Lqual2, Pqual1, and Pqual2).

**Very deep shotgun metagenomics of skin samples allowed the reconstruction of MAGs not yet included in the SMGC (skin microbial genome catalog).**

Very deeply sequenced shotgun metagenomes (>250 billion reads per sample) of selected subjects from cohort A allowed us to reconstruct 164 high-quality MAGs (>90% completeness, <10% contamination) from a pool of 1430 genome bins. While most MAGs were classified as common skin microbes (e.g. *Corynebacterium, Cutibacterium, Staphylococcus, Kocuria,* or *Anaerococcus*), our approach could also retrieve taxa, which were not included in the representative collection of the SMGC yet (Saheb Kashaf et al., 2022). Hence, with our customized genome-centric workflow we could recover MAGs representing 11 new genera (*Capnocytophaga, Gemella, Eubacterium_M, Neofamilia, Leptotrichia_A, Caulobacter, Burkholderia, Serratia, Haemophilus_A, Photobacterium* and *Thermomonas*) and 15 new species, which were not described to be present on human skin yet.

Furthermore, we could show that *Corynebacterium* and Micrococcus had the highest growth rates according to PTR calculations.

In addition, our intensive deep sequencing approach suggests that AOA on human skin are not missed due to their low abundance and/or shallow sequencing. In contrast, these taxa seem to be common but not ubiquitous residents of the human skin microbiome.

**The archaeome on human skin seems to be independent of common disease traits of psoriatic patients.**

20 subjects were recruited and n=72 samples from lesioned and healthy skin samples were taken. However, according to differential abundance analysis based on ANCOM-BC (p adjust = 0.07) and MaAsLin2 (q-value = 0.54) samples from psoriatic skin did not reveal significantly higher or lower AOA counts (Supplementary Fig. 17 and Supplementary Fig. 18, Supplementary Table S27).

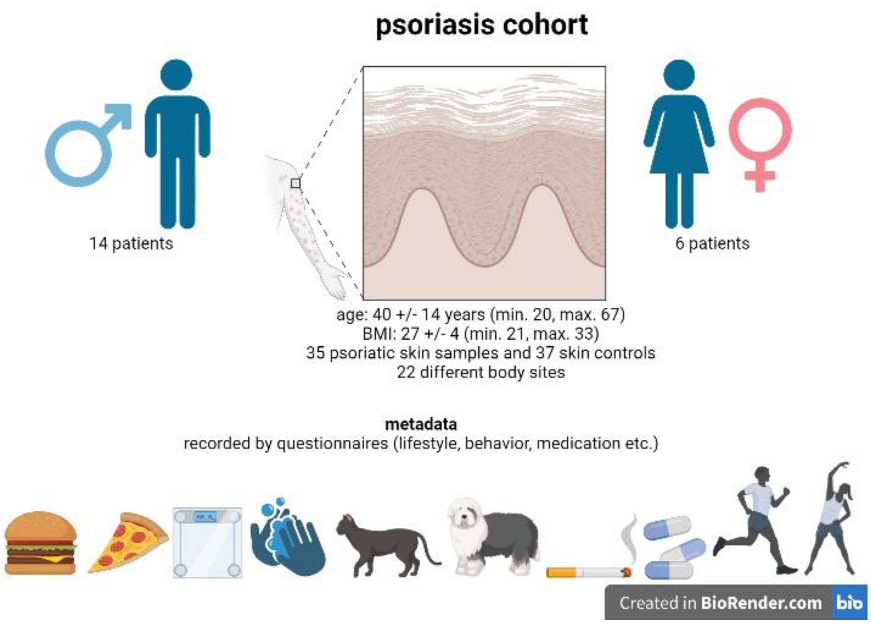


**Supplementary Fig. 17:** Sketch summarizing details about the recruited cohort C covering data types Pqual1 and Pqual 2 (including from top to bottom information about demographics, sampled body sites, and recorded metadata

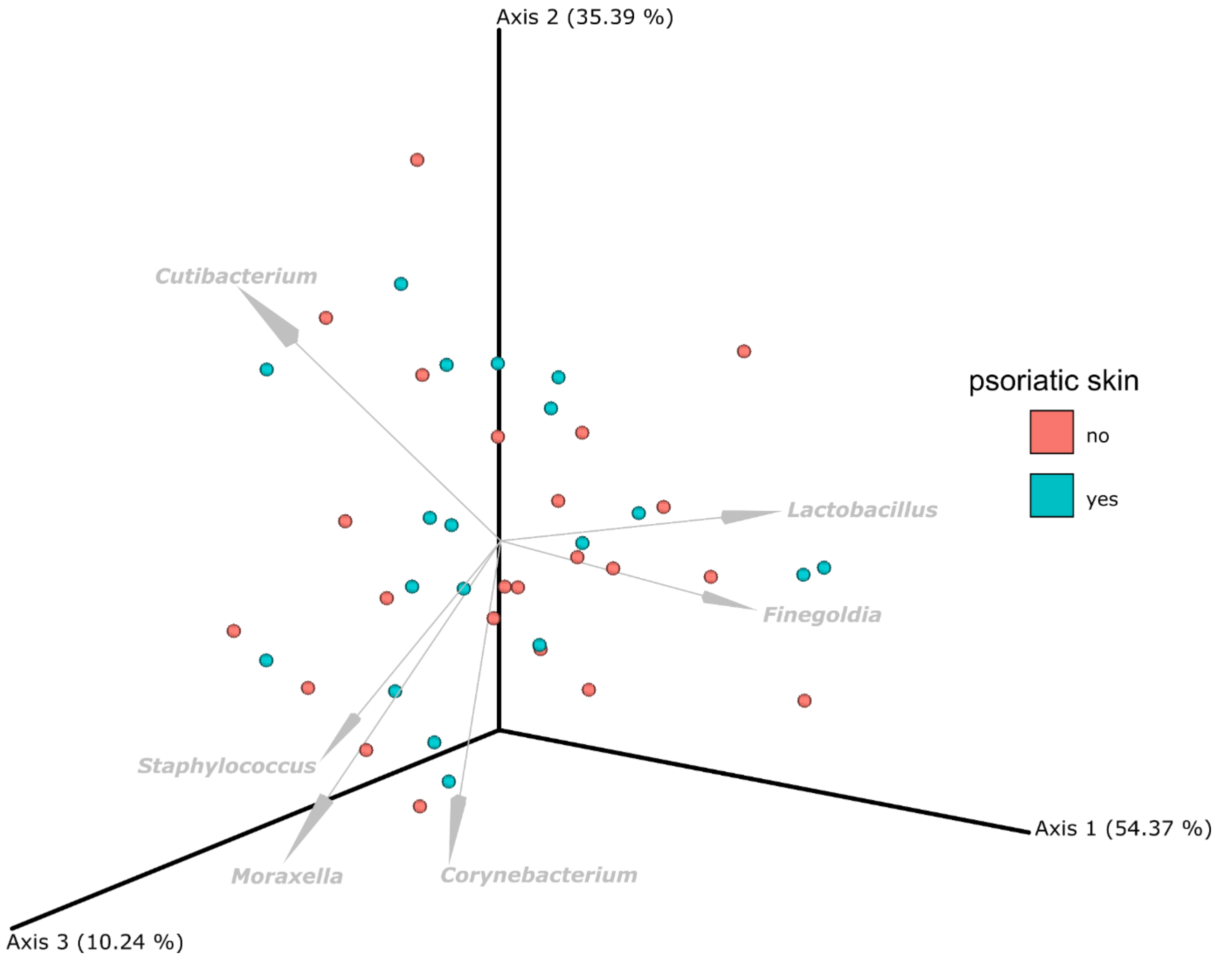


**Supplementary Fig. 18:** Compositional biplots based on robust Aitchison PCA to link specific taxa to beta-diversity. Microbial beta-diversity on affected and non-affected skin sites. The top six bacterial features are shown as superimposed vectors in the biplot (n = 72, data type Pqual1).

### REFERENCES


Abby, S.S., Kerou, M., Schleper, C., 2020. Ancestral reconstructions decipher major adaptations of ammonia-oxidizing archaea upon radiation into moderate terrestrial and marine environments. mBio 11, 1–20. https://doi.org/10.1128/MBIO.02371-20/SUPPL_FILE/MBIO.02371-20-SF004.PDF

Abby, S.S., Melcher, M., Kerou, M., Krupovic, M., Stieglmeier, M., Rossel, C., Pfeifer, K., Schleper, C., 2018. Candidatus Nitrosocaldus cavascurensis, an ammonia oxidizing, extremely thermophilic archaeon with a highly mobile genome. Front Microbiol 9, 346006. https://doi.org/10.3389/FMICB.2018.00028/BIBTEX

Bayer, B., Vojvoda, J., Offre, P., Alves, R.J.E., Elisabeth, N.H., Garcia, J.A.L., Volland, J.-M., Srivastava, A., Schleper, C., Herndl, G.J., 2016. Physiological and genomic characterization of two novel marine thaumarchaeal strains indicates niche differentiation. ISME J 10, 1051–1063. https://doi.org/10.1038/ismej.2015.200

Brandwein, M., Steinberg, D., Meshner, S., 2016. Microbial biofilms and the human skin microbiome. NPJ Biofilms Microbiomes 2, 3. https://doi.org/10.1038/s41522-016-0004-z

Chen, X., Alonzo, F., 2019. Bacterial lipolysis of immune-activating ligands promotes evasion of innate defenses. Proceedings of the National Academy of Sciences 116, 3764–3773. https://doi.org/10.1073/pnas.1817248116

Chua, W., Poh, S.E., Li, H., 2022. Secretory Proteases of the Human Skin Microbiome. Infect Immun 90. https://doi.org/10.1128/IAI.00397-21

Cinar, M.S., Niyas, A., Avci, F.Y., 2024. Serine-rich repeat proteins: well-known yet little-understood bacterial adhesins. J Bacteriol 206. https://doi.org/10.1128/jb.00241-23

Garcia, C.J., Pericleous, A., Elsayed, M., Tran, M., Gupta, S., Callaghan, J.D., Stella, N.A., Franks, J.M., Thibodeau, P.H., Shanks, R.M.Q., Kadouri, D.E., 2018. Serralysin family metalloproteases protects Serratia marcescens from predation by the predatory bacteria Micavibrio aeruginosavorus. Sci Rep 8, 14025. https://doi.org/10.1038/s41598-018-32330-4

Herzog, P.L., Sützl, L., Eisenhut, B., Maresch, D., Haltrich, D., Obinger, C., Peterbauer, C.K., 2019. Versatile Oxidase and Dehydrogenase Activities of Bacterial Pyranose 2-Oxidase Facilitate Redox Cycling with Manganese Peroxidase *In Vitro*. Appl Environ Microbiol 85. https://doi.org/10.1128/AEM.00390-19

Hodgskiss, L.H., Melcher, M., Kerou, M., Chen, W., Ponce-Toledo, R.I., Savvides, S.N., Wienkoop, S., Hartl, M., Schleper, C., 2023. Correction to: Unexpected complexity of the ammonia monooxygenase in archaea. ISME J 17, 947–947. https://doi.org/10.1038/s41396-023-01403-2

Jung, M.Y., Kim, J.G., Sinninghe Damsté, J.S., Rijpstra, W.I.C., Madsen, E.L., Kim, S.J., Hong, H., Si, O.J., Kerou, M., Schleper, C., Rhee, S.K., 2016. A hydrophobic ammonia-oxidizing archaeon of the Nitrosocosmicus clade isolated from coal tar-contaminated sediment. Environ Microbiol Rep 8, 983–992. https://doi.org/10.1111/1758-2229.12477

Klein, T., Poghosyan, L., Barclay, J.E., Murrell, J.C., Hutchings, M.I., Lehtovirta-Morley, L.E., 2022. Cultivation of ammonia-oxidising archaea on solid medium. FEMS Microbiol Lett 369. https://doi.org/10.1093/femsle/fnac029
Kumar, N.G., Contaifer, D., Wijesinghe, D.S., Jefferson, K.K., 2021. Staphylococcus aureus Lipase 3 (SAL3) is a surface-associated lipase that hydrolyzes short chain fatty acids. PLoS One 16, e0258106. https://doi.org/10.1371/journal.pone.0258106
Lehtovirta-Morley, L.E., Ross, J., Hink, L., Weber, E.B., Gubry-Rangin, C., Thion, C., Prosser, J.I., Nicol, G.W., 2016. Isolation of ‘Candidatus Nitrosocosmicus franklandus’, a novel ureolytic soil archaeal ammonia oxidiser with tolerance to high ammonia concentration. FEMS Microbiol Ecol 92, 57. https://doi.org/10.1093/FEMSEC/FIW057
Lien, K.A., Dinshaw, K., Nichols, R.J., Cassidy-Amstutz, C., Knight, M., Singh, R., Eltis, L.D., Savage, D.F., Stanley, S.A., 2021. A nanocompartment system contributes to defense against oxidative stress in Mycobacterium tuberculosis. Elife 10. https://doi.org/10.7554/eLife.74358
Liu, L., Li, S., Han, J., Lin, W., Luo, J., 2019. A Two-Step Strategy for the Rapid Enrichment of Nitrosocosmicus-Like Ammonia-Oxidizing Thaumarchaea. Front Microbiol 10. https://doi.org/10.3389/fmicb.2019.00875
Liu, L., Liu, M., Jiang, Y., Lin, W., Luo, J., 2021. Production and Excretion of Polyamines To Tolerate High Ammonia, a Case Study on Soil Ammonia-Oxidizing Archaeon “ Candidatus Nitrosocosmicus agrestis.” mSystems 6. https://doi.org/10.1128/MSYSTEMS.01003-20
Moissl-Eichinger, C., Probst, A.J., Birarda, G., Auerbach, A., Koskinen, K., Wolf, P., Holman, H.-Y.N., 2017. Human age and skin physiology shape diversity and abundance of Archaea on skin. Sci Rep 7. https://doi.org/10.1038/s41598-017-04197-4
Mondal, A.K., Lata, K., Singh, M., Chatterjee, S., Chauhan, A., Puravankara, S., Chattopadhyay, K., 2022. Cryo-EM elucidates mechanism of action of bacterial pore-forming toxins. Biochimica et Biophysica Acta (BBA) - Biomembranes 1864, 184013. https://doi.org/10.1016/j.bbamem.2022.184013
Ni, Q., Zhang, P., Li, Q., Han, Z., 2022. Oxidative Stress and Gut Microbiome in Inflammatory Skin Diseases. Front Cell Dev Biol 10, 849985. https://doi.org/10.3389/fcell.2022.849985
Probst, A.J., Auerbach, A.K., Moissl-Eichinger, C., 2013. Archaea on Human Skin. PLoS One 8, e65388. https://doi.org/10.1371/journal.pone.0065388
Reyes, C., Hodgskiss, L.H., Baars, O., Kerou, M., Bayer, B., Schleper, C., Kraemer, S.M., 2020. Copper limiting threshold in the terrestrial ammonia oxidizing archaeon Nitrososphaera viennensis. Res Microbiol 171, 134–142. https://doi.org/10.1016/J.RESMIC.2020.01.003
Spang, A., Poehlein, A., Offre, P., Zumbrägel, S., Haider, S., Rychlik, N., Nowka, B., Schmeisser, C., Lebedeva, E. V., Rattei, T., Böhm, C., Schmid, M., Galushko, A., Hatzenpichler, R., Weinmaier, T., Daniel, R., Schleper, C., Spieck, E., Streit, W., Wagner, M., 2012. The genome of the ammonia-oxidizing Candidatus Nitrososphaera gargensis: insights into metabolic versatility and environmental adaptations. Environ Microbiol 14, 3122–3145. https://doi.org/10.1111/j.1462-2920.2012.02893.x
Sützl, L., Laurent, C.V.F.P., Abrera, A.T., Schütz, G., Ludwig, R., Haltrich, D., 2018. Multiplicity of enzymatic functions in the CAZy AA3 family. Appl Microbiol Biotechnol 102, 2477–2492. https://doi.org/10.1007/s00253-018-8784-0

Tourna, M., Stieglmeier, M., Spang, A., Könneke, M., Schintlmeister, A., Urich, T., Engel, M., Schloter, M., Wagner, M., Richter, A., Schleper, C., 2011. Nitrososphaera viennensis, an ammonia oxidizing archaeon from soil. Proc Natl Acad Sci U S A 108, 8420–8425. https://doi.org/10.1073/PNAS.1013488108/SUPPL_FILE/PNAS.201013488SI.PDF

Zou, Q., Luo, S., Wu, H., He, D., Li, X., Cheng, G., 2020. A GMC Oxidoreductase GmcA Is Required for Symbiotic Nitrogen Fixation in Rhizobium leguminosarum bv. viciae. Front Microbiol 11. https://doi.org/10.3389/fmicb.2020.00394
