## Supplementary Figures for "Molecular Tracking and Cultivation Reveal Ammonia-Oxidizing Archaea as Emerging Commensals of the Human Skin Microbiome"

### Extended Data Figures

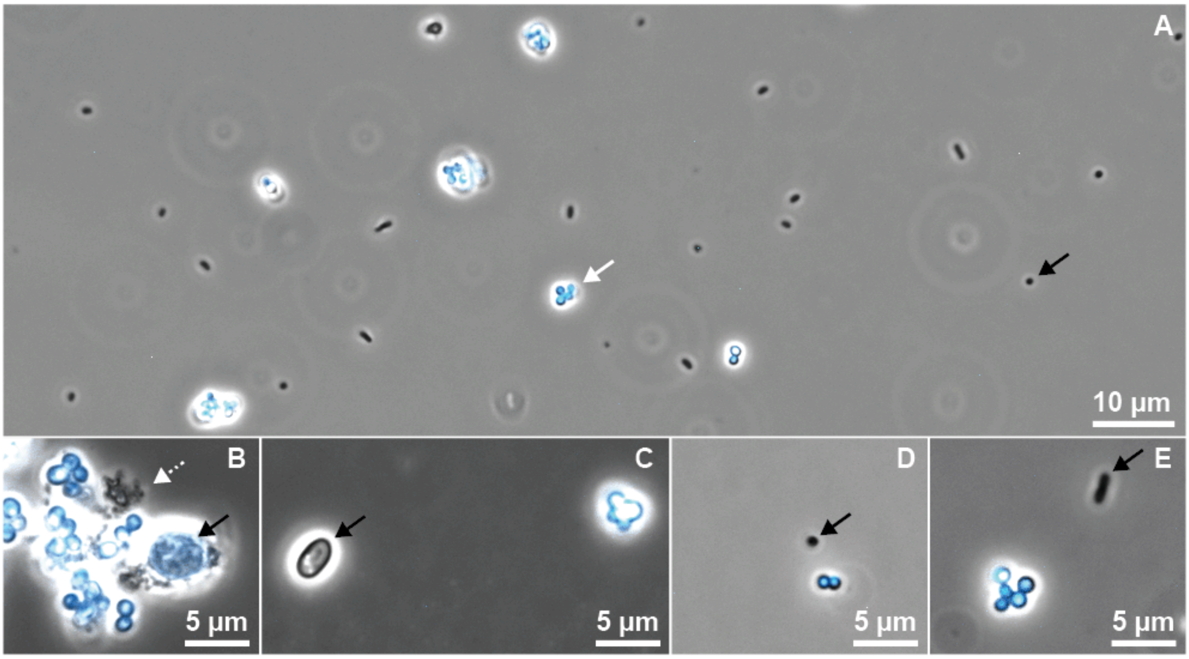


**Extended Data Fig. 1:**
Identification of AOA based on F420 autofluorescence by light microscopy.
(A, E) Enrichment Z3A. (B) Enrichment R2S. (C) Enrichment T1S. (D) Enrichment X2B. The autofluorescence of AOA can be used to distinguish between AOA and other contaminants in complex enrichment cultures (A). Putative ammonia-oxidizing archaea were visualized by F420 autofluorescence false-colored in cyan (white arrow (A), cyan (B-E)). Contaminants were not autofluorescent (black arrows), except for minimal autofluorescence of the contaminants associated with aggregates of AOA of enrichment R2S (B, black arrow). The dashed white arrow in (B) indicates putative extracellular polymeric substances (EPS). Images are overlays of phase contrast light microscopy and false-colored F420 fluorescence microscopy using a standard DAPI filter cube. Enrichment Z3A was concentrated 10X times via centrifugation before imaging.

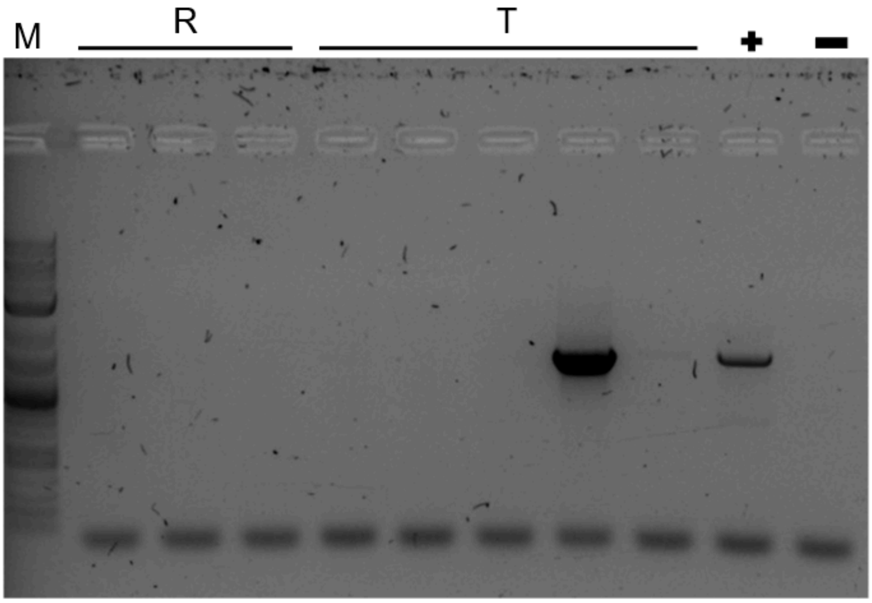


**Extended Data Fig. 2.** Agarose gel of bacterial 16S rRNA gene PCR of enrichments R2S and T1S. Multiple cultures of enrichments R and T were screened for the presence of bacterial contaminants via amplification of bacterial 16S rRNA genes (PCR, 33 cycles). Cultures free of bacterial contaminants were used as inoculum for future experiments. M: Marker, GeneRuler 1kb plus, +: positive control, -: negative control

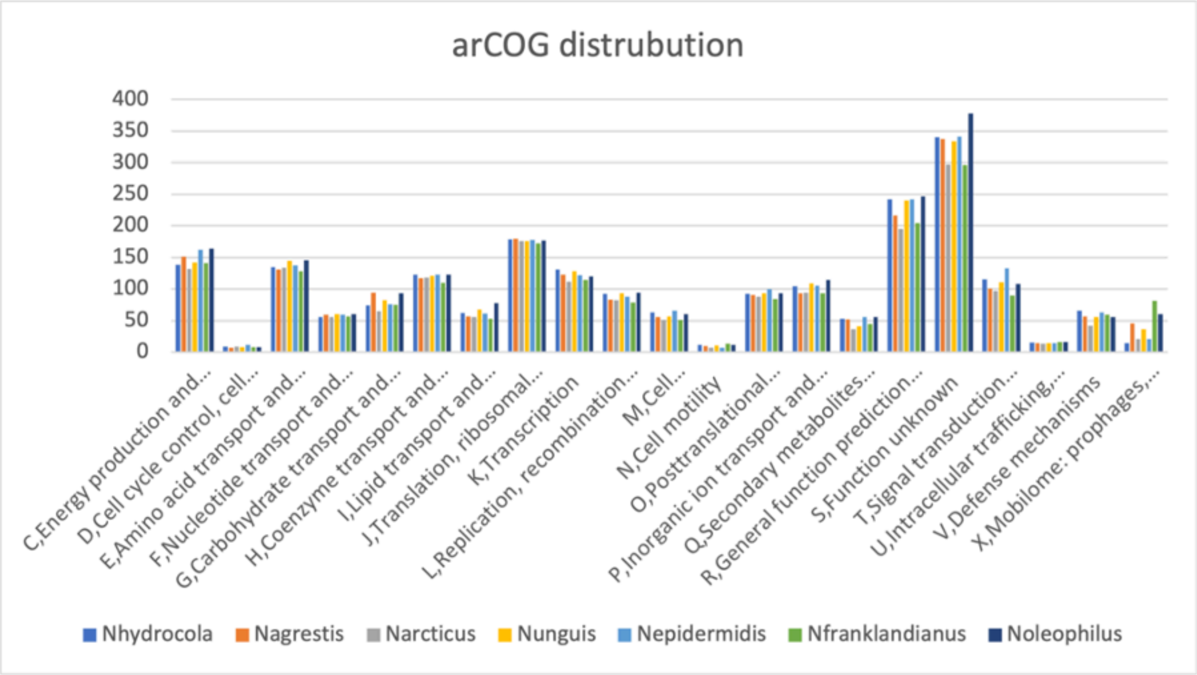


**Extended Data Fig. 3**: Distribution of arCOG functional categories among seven representative genomes of *Ca.* Nitrosocosmicus

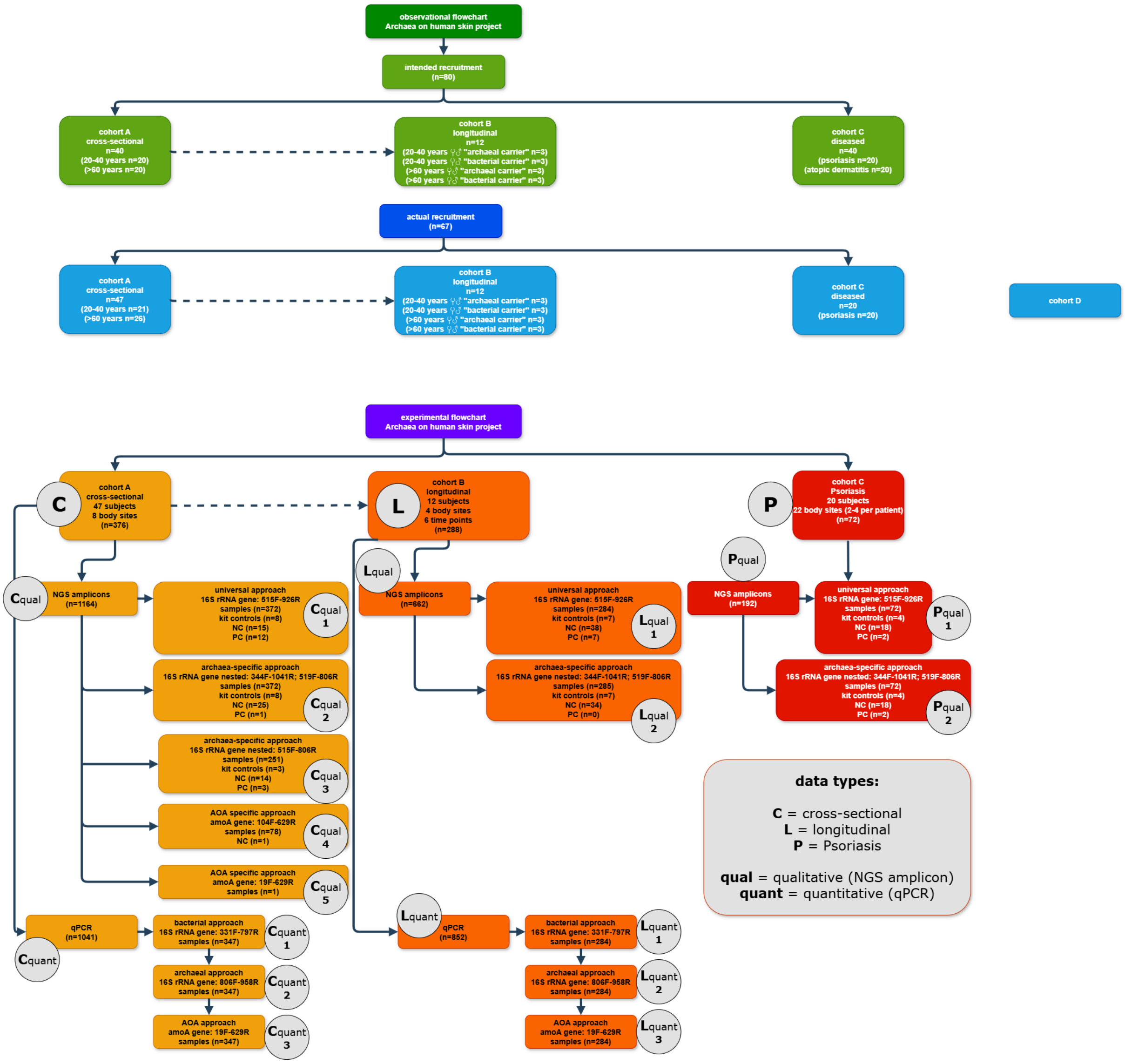


**Extended Data Figure 4** STORM chart: STORM flowchart of the project with identifiers for each different data type.

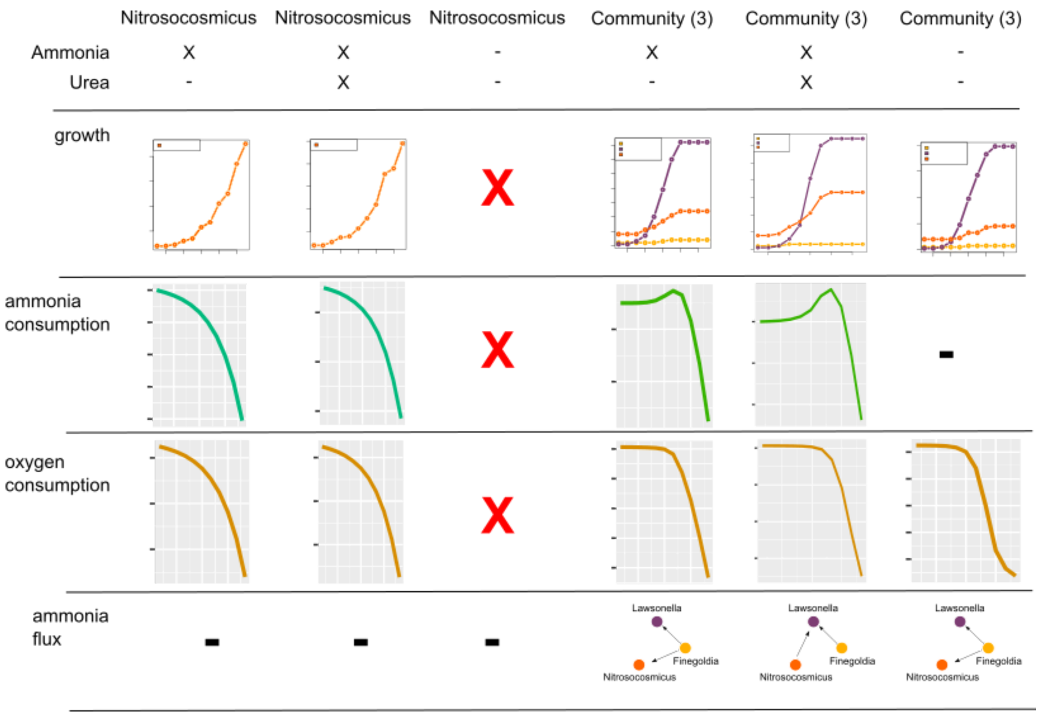


**Extended Data Fig. 5**: In silico metabolic modeling with BacArena showing modeled growth, ammonia consumption, oxygen consumption, and ammonia flux for *Ca.* Nitrosocosmicus alone and the co-occurring community of *Ca.* Nitrosocosmicus with Lawsonella and Finegoldia (see Fig. 5) with and without ammonia and urea as supplements in the provided media.

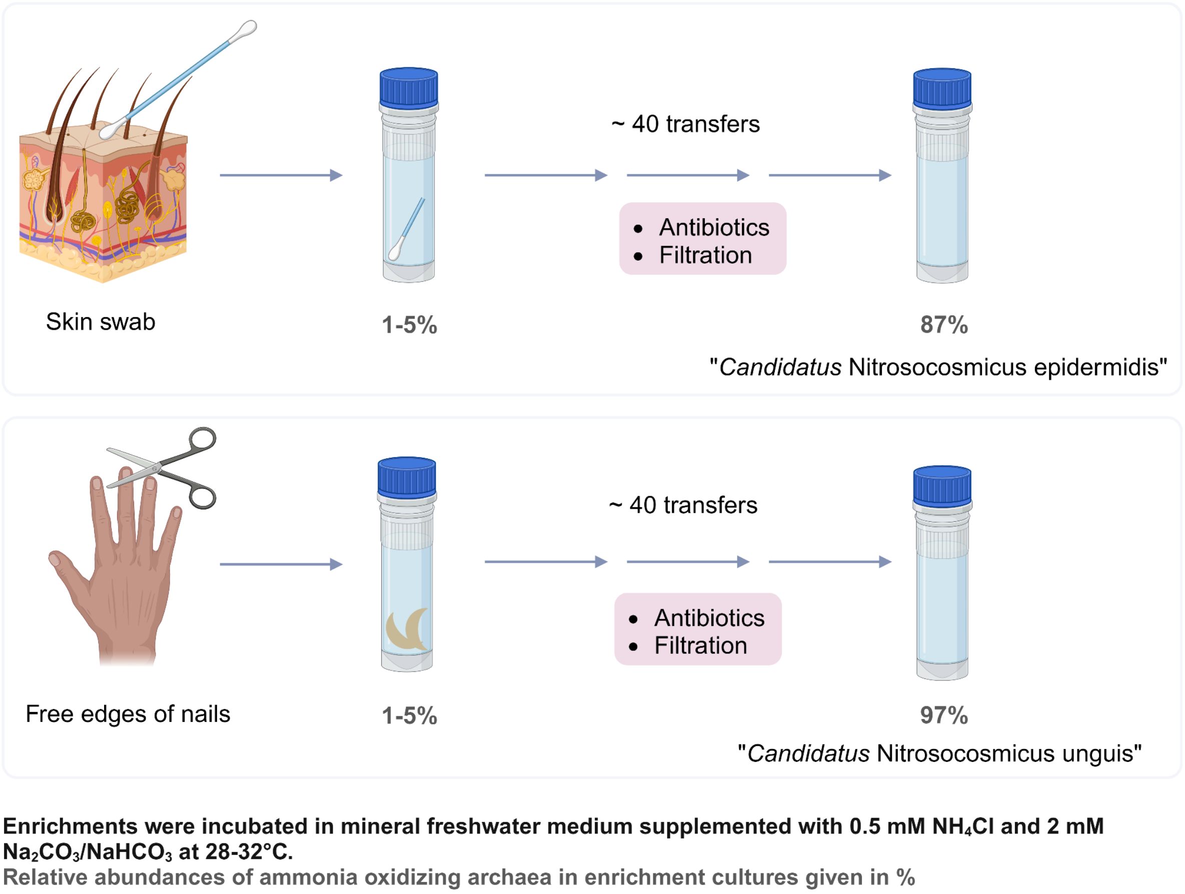


**Extended Data Fig. 6:** Enrichment scheme for skin-residing AOA. Created with BioRender.com.
